# CaBLAM! A high-contrast bioluminescent Ca^2+^ indicator derived from an engineered *Oplophorus gracilirostris* luciferase

**DOI:** 10.1101/2023.06.25.546478

**Authors:** Gerard G. Lambert, Emmanuel L. Crespo, Jeremy Murphy, Kevin Turner, Emily Gershowitz, Michaela Cunningham, Daniela Boassa, Selena Luong, Dmitrijs Celinskis, Justine J. Allen, Stephanie Venn, Yunlu Zhu, Mürsel Karadas, Jiakun Chen, Roberta Marisca, Hannah Gelnaw, Daniel K. Nguyen, Junru Hu, Brittany N. Sprecher, Maya O. Tree, Richard Orcutt, Daniel Heydari, Aidan B. Bell, Albertina Torreblanca-Zanca, Ali Hakimi, Tim Czopka, Shy Shoham, Katherine I. Nagel, David Schoppik, Arturo Andrade, Diane Lipscombe, Christopher I. Moore, Ute Hochgeschwender, Nathan C. Shaner

## Abstract

Measuring ongoing cellular activity is essential to understanding the dynamic functions of biological organisms. The most popular current approach is imaging fluorescence-based genetically encoded Ca^2+^ indicators (GECIs). While fluorescent probes are useful in many contexts, bioluminescence-based GECIs—probes that generate light through oxidation of a small-molecule by a luciferase or photoprotein—have several distinct advantages. Because bioluminescent (BL) GECIs do not use the bright extrinsic excitation light required for fluorescence, BL GECIs do not photobleach, do not suffer from nonspecific autofluorescent background, and do not cause phototoxicity. Further, BL GECIs can be applied in contexts where directly shining photons on an imaging target is not possible. Despite these advantages, the use of BL GECIs has to date been limited by their small changes in bioluminescence intensity, high baseline signal at resting Ca^2+^ concentrations, and suboptimal Ca^2+^ affinities. Here, we describe a new BL GECI, CaBLAM (<u>Ca</u>^2+^ <u>B</u>io<u>L</u>uminescence <u>A</u>ctivity <u>M</u>onitor), that displays much higher dynamic range than previous BL GECIs and has a Ca^2+^ affinity suitable for capturing physiological changes in cytosolic Ca^2+^ concentration. With these improvements, CaBLAM captures single-cell and subcellular resolution activity at high frame rates in cultured neurons and *in vivo*, and allows multi-hour recordings in mice and behaving zebrafish. This new advance provides a robust alternative to traditional fluorescent GECIs that can enable or enhance imaging across many experimental conditions.

## Introduction

Genetically encoded Ca^2+^ indicators (GECIs) play a central role in modern biomedical research, with much of our current understanding of systems neuroscience based on Ca^2+^ imaging. GECIs based on fluorescent proteins (FPs) have been the subject of intense engineering efforts for decades, leading to continual improvements to the practical properties of this class of biosensor^1–4^. The GCaMP family of GECIs is currently in its eighth revision^5^, with improvements in brightness, sensitivity, and response kinetics, and new families of FP-based GECIs continue to push the boundaries of performance^6–9^.

While powerful, fluorescence imaging has key limitations. Most importantly, obtaining fluorescent signals requires high intensity photon excitation. This bombardment typically causes photobleaching and high background autofluorescence, undermining sensitivity and spatial precision. Intense excitation can also cause photodamage, severely limiting the long-term (e.g., whole lifetime) imaging of cells. The optics required to deliver this light add hardware and fix the light path, challenges that are especially troublesome in experiments with freely moving animals. Bioluminescent (BL) GECIs remove these hurdles by generating light through an enzyme–driven reaction. The approach began with the natural *Aequorea victoria* photoprotein, aequorin^10–12^, and has since grown to include a number of luciferase-based BL GECIs^13–18^. Despite these advances, no available BL GECI has achieved imaging performance comparable to highly optimized fluorescent indicators such as the GCaMP series. Existing BL GECIs typically afford limited dynamic range *in vivo* owing to high baseline emission and Ca^2+^ affinities that reduce sensitivity in the physiological range, restricting their practical application to population–level recordings.

Here, we describe the development of SSLuc (“<u>S</u>ensor <u>S</u>caffold <u>Luc</u>iferase”), a variant of *Oplophorus gracilirostris*^19–21^ luciferase (OLuc) engineered to provide increased BL emission *in vitro* along with improved folding and performance in protein fusions. SSLuc is several-fold more active than NanoLuc *in vitro* using the widely employed furimazine (Fz) substrate and is less prone to aggregation in cells than NanoLuc when fused to other proteins. We found that SSLuc was highly amenable to sensor domain insertion, making it a favorable scaffold for engineering a new BL GECI, CaBLAM. Optimization of the C-terminal peptide sequence was a critical component of both luciferase and GECI engineering, and targeted mutations to the calmodulin domain of the sensor were particularly important for obtaining the high contrast currently unique to CaBLAM.

CaBLAM has similar maximum light output to other described BL GECIs in cells but displays ∼83-fold full-range contrast *in vitro* and up to 15-20-fold in live cultured cells in response to physiological changes in cytosolic Ca^2+^ levels. We further demonstrate that CaBLAM reliably reports the Ca^2+^ signal from a field stimulation equivalent to a single action potential when imaged at single-cell resolution at 10 Hz in cultured primary neurons on a typical widefield microscope and EMCCD camera. In mice, CaBLAM was also readily imaged at 10 Hz in head-fixed awake animals, where its ability to report physiologically relevant neural Ca^2+^ activity evoked by vibrissa stimulation at the single-cell level was superior to GCaMP6s under widefield single-photon (1-photon) illumination. In zebrafish, we demonstrate CaBLAM detection in several genetically targeted transgenic lines in awake head-fixed larvae, with sampling rates of 40 Hz using a photomultiplier tube (PMT) or 20 Hz using an intensified camera. In both cases, CaBLAM produced robust signals following large tail movements with kinetics tuned to cell type.

## Results

### Rational design and directed evolution of high-activity soluble OLuc variants

We initially set out to develop a variant of OLuc that could more efficiently utilize coelenterazine (CTZ), the native substrate of many marine luciferases including OLuc, reasoning that NanoLuc was only one of the possible endpoints achievable through directed evolution of this lineage of luciferases. Starting from the OLuc mutant eKAZ^22^, we performed multiple rounds of directed evolution consisting of alternating error-prone and site-directed mutagenesis libraries in *E. coli*. For each round, variants were selected based on higher BL emission and solubility. Subsequent engineering of the intact luciferase included the generation of mNeonGreen^23^ fusions for enhancement of emission quantum yield, incorporation of a subset of mutations found in NanoLuc^24^ and NanoBit^25^, optimization of the C-terminal peptide sequence, and fine-tuning interactions between the C-terminal peptide and the rest of the protein. The final clone, which we named **SSLuc** (“Sensor Scaffold Luciferase”, **Fig. 1A**), can be extracted with nearly 100% efficiency from *E. coli* cultures, retains activity *in vitro* over longer storage periods than NanoLuc, and appears to behave more like a monomer in mammalian cells than NanoLuc.

**Figure 1.**
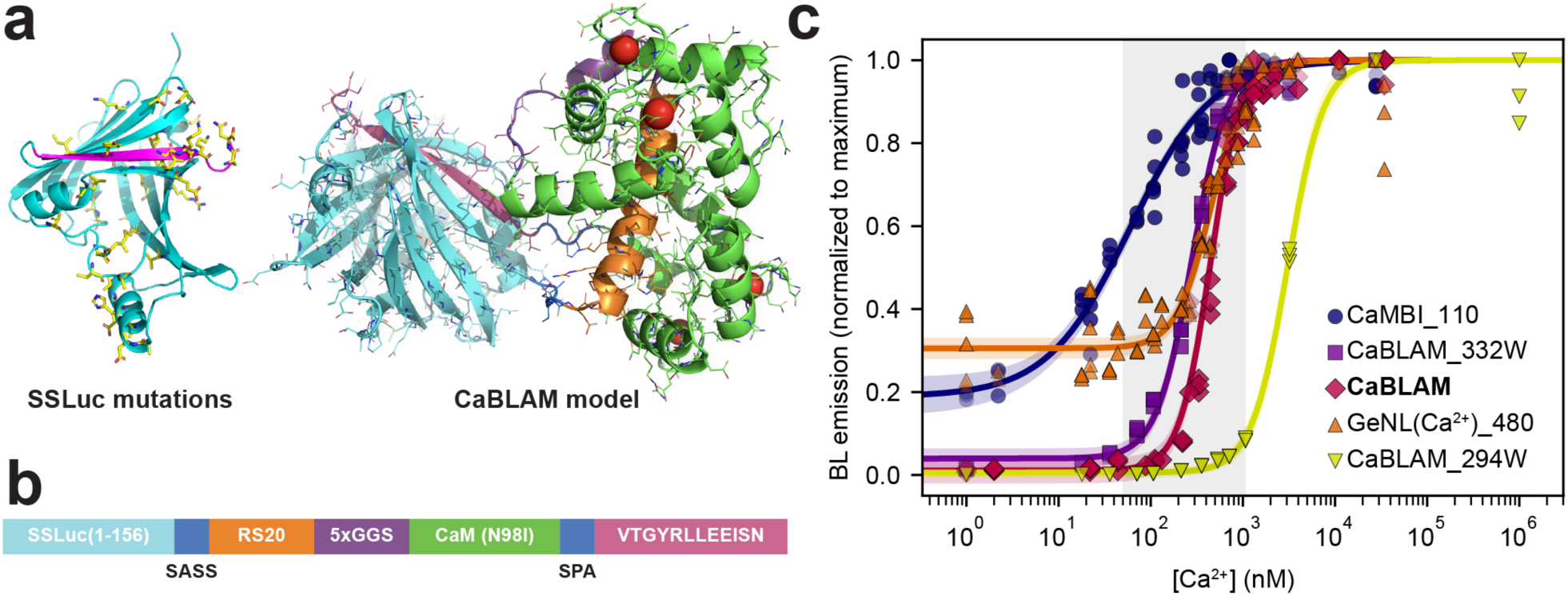
CaBLAM architecture and Ca^2+^ affinity. **a,** Structure models of eKL9h and CaBLAM. Side chain positions with mutations relative to eKAZ are depicted as yellow sticks in the luciferase model. The C-terminal peptide is depicted in magenta on both structure models. **b,** Architecture of the sensor component of CaBLAM (not to scale), with colors corresponding to the model structure backbone. Full-length CaBLAM includes the N-terminal mNeonGreen FRET acceptor. **c,** *In vitro* Ca^2+^ titration of CaBLAM (*n* = 6, diamonds) alongside GeNL(Ca^2+^)_480 (*n* = 8, triangles), CaMBI_110 (*n* = 6, circles), and two CaBLAM variants, 294W (*n* = 3, inverted triangles) and 332W (*n* = 3, squares), with altered Ca^2+^ affinities. The given *n* value in parentheses for each BL GECI represents the number of technical replicates of the full titration performed. All individual data points are shown. BL intensities were normalized to the maximum BL within each replicate dataset, pooled for each indicator, and globally fitted to a 3-parameter Hill model. Fit curves are shown for each with shaded areas representing 95% CI. The typical physiologically relevant cytosolic Ca^2+^ concentration range between ∼50 nM and ∼1 μM is shaded in grey.

Contrary to our original intentions, SSLuc performs best using Fz as its luciferin, though its activity with other CTZ analogs is also higher than we measure for NanoLuc. Interestingly, we found that SSLuc does not appear brighter than NanoLuc when expressed in mammalian cells, despite its high performance *in vitro*, suggesting that availability of the luciferin substrate (e.g., Fz) may limit SSLuc’s maximum brightness in cells. SSLuc proved to be a favorable luciferase scaffold from which to generate biosensors, and led us to develop a novel high-performance BL GECI, as described below. A detailed account of our development and characterization of SSLuc can be found in **Supplementary Results and Discussion, Supplementary Table 1, and Supplementary** Figures 1-3.

### Development and evaluation of a high-contrast BL GECI

To engineer a high-contrast bioluminescent GECI, we split SSLuc in the loop joining its final two β-strands and inserted Ca²⁺-sensing domains, testing circular permutations and direct fusions of RS20–CaM, CaM–RS20, and Troponin C (**Supp. Fig. 4**). All prototypes responded robustly *in vitro*, but the RS20–CaM topology (**Fig. 1A,B**) produced the largest BL change (**Fig. 1A,B**), echoing a hybrid NanoBit-GCaMP based design (GLICO^26^) while retaining SSLuc’s fixed N-terminal fluorescent partner for FRET enhancement^17^ and restricting Ca²⁺ readout to the BL channel. Early clones had very high Ca^2+^ affinity (*K_D_* < 10 nM), so we iteratively tuned affinity and contrast by swapping RS20/CaM sequences from fluorescent GECIs^27–29^, shortening or extending inter-domain linkers, and introducing rational EF-hand mutations (**Supp. Fig. 4)**. The optimal combination we identified utilized the GCaMP6s RS20/CaM pair combined with a naturally occurring CaM mutation, N98I^30^, that lowers the apparent Ca^2+^ affinity (**Fig. 1C**, **Table 1**), aligning the sensor’s most sensitive range with typical cytosolic Ca²⁺ levels observed in hippocampal neurons^31^. To maximize contrast, we next systematically evaluated variants in this framework using the collection of C-terminal peptides we had generated while developing SSLuc (**Supp. Results and Discussion, Supp. Table 1**), finding that the low-affinity sequence **VTGYRLFEEIL** (“pep114” ^25^) produced the greatest increase in BL emission between low and high Ca^2+^ concentrations. Based on these results, we redesigned the optimized C-terminal peptide from SSLuc to arrive at **VTGYRLLEEISN**, which preserves the catalytic Arg while replacing Lys residues with Glu to weaken peptide–enzyme binding and retain high contrast. This final architecture (**Fig. 1B, Supp. Fig. 4D**) was dubbed **CaBLAM**.

**Table 1.** Properties of selected intensiometric genetically encoded Ca^2+^ indicators.

| GECI Name | Type | $K_D$ (nM)<sup>a</sup> | Hill coefficient<sup>b</sup> | Contrast<sup>c</sup> | ref. |
| --- | --- | --- | --- | --- | --- |
| **GCaMP8s** | FP | 46 | 2.2 | 50 | 5 |
| **GCaMP8f** | FP | 334 | 2.1 | 79 | 5 |
| **NEMOf** | FP | 528 | 3.2 | 246 | 6 |
| **NEMOc** | FP | 557 | 3.8 | 422 | 6 |
| **CaMBI_110** | luciferase | $59 \pm 6$ (110) | $0.97 \pm 0.06$ (2.5) | $4.8 \pm 0.1$ (8) | 18 |
| **GeNL($\text{Ca}^{2+}$)_480** | luciferase | $457 \pm 20$ (480) | $2.28 \pm 0.21$ (3) | $3.6 \pm 0.3$ (5) | 17 |
| **CaBLAM** | luciferase | $439 \pm 14$ | $2.62 \pm 0.18$ | $83 \pm 9$ | this work |
| **CaBLAM_294W** | luciferase | $3067 \pm 95$ | $2.24 \pm 0.19$ | $624 \pm 39$ | this work |
| **CaBLAM_332W** | luciferase | $281 \pm 9$ | $2.26 \pm 0.14$ | $68 \pm 5$ | this work |

<sup>a</sup>Binding constant for $\text{Ca}^{2+}$ for FP-based GECIs as reported in the original publications; binding constant for $\text{Ca}^{2+}$ as measured *in vitro* for CaMBI, GeNL( $\text{Ca}^{2+}$ )\_480, CaBLAM, and CaBLAM variants and (in parentheses) as reported in the original publications. <sup>b</sup>Hill coefficient (cooperativity) for FP-based GECIs as reported in the original publications; Hill coefficient for luciferase-based GECIs calculated from sigmoidal curve fit to $\text{Ca}^{2+}$ titration series and (in parentheses) as reported in the original publications. <sup>c</sup>Contrast between $\text{Ca}^{2+}$ -free and $\text{Ca}^{2+}$ -saturated indicator *in vitro* for FP-based GECIs as reported in the original publications; contrast for luciferase-based GECIs measured in this work and (in parentheses) as reported in the original publications. Error margins represent 95% CI for $K_D$ and Hill coefficient and s.e.m. for contrast, as measured in this work.

We characterized purified CaBLAM protein *in vitro,* determining a *K_D_* of ∼439 nM for Ca^2+^ binding and an absolute contrast (i.e., the ratio of luminescence emission between saturating Ca^2+^ and zero Ca^2+^, typically reported for this class of sensor) of ∼83-fold (**Fig. 1C**, **Table 1**). This value places CaBLAM favorably relative to other BL GECIs published to date, all of which display substantially lower contrast than CaBLAM^14,17,18^. The Hill coefficient of CaBLAM is ∼2.6, similar to that of GeNL(Ca^2+^)_480^17^ (reported as 3 (^17^), measured at ∼2.3 in this study) and notably higher than that of CaMBI^18^, which displays a Hill coefficient of ∼1 in our hands (**Fig. 1C**, **Table 1**). We also characterized two additional EF-hand mutants of CaBLAM that retain high brightness and contrast but have higher and lower Ca^2+^ affinities (**Table 1, Supp. Data File 1**): 294W (N98W in CaM; *K_D_* ∼3 μM) and 332W (Q135W in CaM; *K_D_* ∼280 nM).

Since CaBLAM has a similar Ca^2+^ affinity to GeNL(Ca^2+^)_480 and shares the same mNeonGreen FRET acceptor, we chose these two BL GECIs as a matched pair for benchmarking in cells. In practice, when a GECI is expressed in a living system, its contrast typically drops dramatically relative to *in vitro* values, and so we next evaluated the behavior of CaBLAM in cultured cells in comparison with GeNL(Ca^2+^)_480 to determine how much of its promising *in vitro* behavior would translate to improved real-world performance.

### Characterization of CaBLAM in cells and benchmarking against GeNL(Ca^2+^)_480

CaBLAM markedly outperformed the NanoLuc-derived indicator GeNL(Ca²⁺)_480 across cell lines and primary neurons in terms of contrast. In U2OS cells treated with ionomycin, CaBLAM produced ∼5.4-fold higher contrast between resting and high Ca^2+^ versus GeNL(Ca²⁺)_480 (**Supp. Fig. 5A**), owing to CaBLAM’s low baseline luminescence and favorable Hill coefficient. CaBLAM’s performance in HeLa cells showed similar advantages, generating L-histamine-induced BL oscillations that were ∼3.3-fold larger than those for GeNL(Ca^2+^)_480 and displaying diverse oscillatory phenotypes that ranged from seconds to minutes (**Supp. Fig. 5B, D, Supp. Movie 1**). Primary rat cortical neurons exhibited ∼20-fold bioluminescence increase after KCl depolarization versus ∼4-fold with GeNL(Ca²⁺)_480 (**Supp. Fig. 5C**), leading us to begin exploring CaBLAM’s performance relative to fluorescent GECIs as well.

We compared the high-Ca^2+^ brightness (maximal brightness) of CaBLAM compared with GeNL(Ca^2+^)_480 by normalizing the BL signal in ionomycin-treated HeLa cells to the mNeonGreen fluorescence signal for each. However, as with SSLuc, we observed an unexpectedly weak correlation between fluorescence and BL brightness in all but the lowest-expressing cells for both GECIs, which we suspect is due to rate-limiting substrate diffusion across the plasma membrane. Analysis of only the cells with low expression for each sensor indicates that CaBLAM is ∼39% ± 19% as bright as GeNL(Ca^2+^)_480 on a per-molecule basis (95%CI). The uncertainty in this estimate is due to the weak linear correlation between fluorescence and BL intensity for both indicators, even at low expression levels (**Supp Fig. 6**).

### Comparison of CaBLAM to GCaMP8s in rat hippocampal neurons

We performed simultaneous Ca^2+^ imaging and electrical field stimulation (FS) to characterize the CaBLAM sensor (**Supp. Movie 2**). Benchmarking against the state-of-the-art fluorescent sensor GCaMP8s in rat hippocampal neurons. CaBLAM-expressing neurons showed low detectable photon counts during 10 Hz imaging, yet individual traces demonstrated detectable evoked responses across a range of electrical field stimulations (**Fig. 2A** and **C**). As expected for a fluorescent indicator, GCaMP8s was brighter overall and captured both spontaneous and evoked neural activity (**Fig. 2B and D**). These initial observations represent typical examples of the responses observed for each indicator. We calculated the evoked change in bioluminescence for CaBLAM, showing that it reliably reported changes in intracellular Ca^2+^, with large changes observed in the normalized (ΔL/L) signal (**Fig. 2C**). For GCaMP8s (**Fig. 2D**), the normalized change in fluorescence (ΔF/F) was reduced due to its high baseline fluorescence. Despite this, GCaMP8s still produced observable changes in fluorescence, consistent with previous reports.

**Figure 2.**
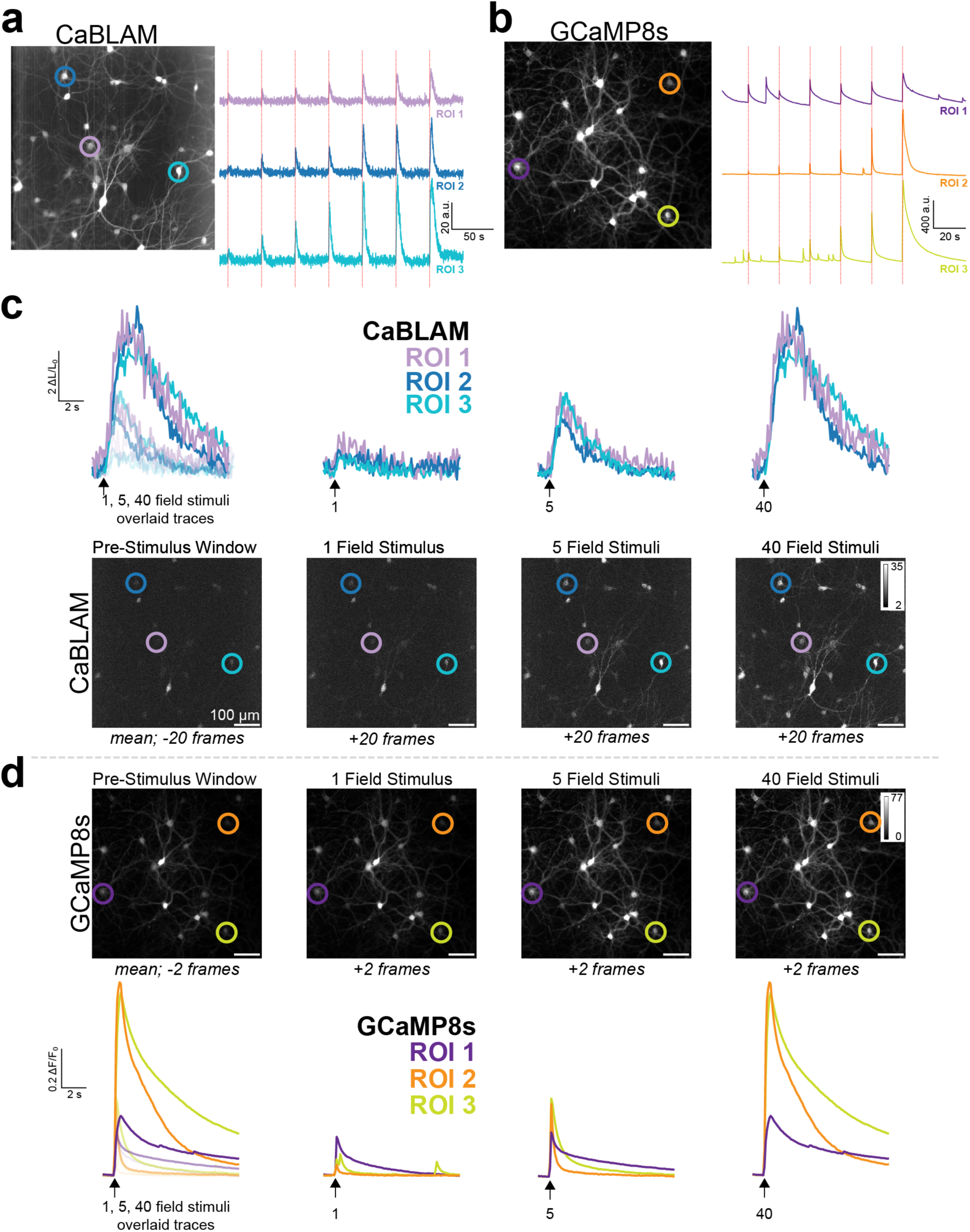
Overview of a typical Ca^2+^ imaging session comparing electrical field evoked responses in CaBLAM and GCaMP8s expressing rat hippocampal neurons. **a,** Representative CaBLAM bioluminescence imaging session. *left*: Mean z-stacked image of with regions of interest (ROIs) highlighted by colored circles (ROI 1 in light purple, ROI 2 in blue, and ROI 3 in teal). *righ*t: BL Ca^2+^ transients from ROIs 1-3 displaying responses evoked by multiple field stimuli (red dashed lines indicate field stimulation events). **b,** Representative GCaMP8s fluorescence imaging session. *left*: Mean z-stacked image with individual ROIs highlighted by colored circles (ROI 1 in dark purple, ROI 2 in orange, ROI 3 in yellow). *right*: Fluorescence Ca^2+^ transients from ROIs 1-3 displaying responses evoked by multiple field stimuli (red dashed lines indicate stimulation events). For both **a** and **b**, values are in arbitrary units (a.u.). **c,** CaBLAM Ca^2+^ responses to varied field stimuli. *top*: Individual BL Δ L/L Ca^2+^ response traces from ROIs 1-3 for each field stimulus condition. *bottom*: Mean z-stacked frames (20 frames per condition) for pre-stimulus, 1 field stimulus, 5 field stimuli, and 40 field stimuli conditions, corresponding to data shown in the top panel. **d,** GCaMP8s Ca^2+^ responses to varied field stimuli. *top*: Individual fluorescence ΔF/F Ca^2+^ response traces from ROIs 1-3 for each stimulus condition. *bottom*: Mean z-stacked frames (2 frames per condition) for pre-stimulus, 1 field stimulus, 5 field stimuli, and 40 field stimuli conditions, corresponding to data shown in the top panel. For **c** and **d**, arrows indicate field stimulus onset.

We next characterized stimulus-evoked bioluminescence and fluorescence signals in neurons expressing CaBLAM **(Fig. 3A**) and GCaMP8s (**Fig. 3B**). Consistent with our initial observation, electrical field stimulations (1, 5, and 40 pulses at ∼83 Hz, 1 ms pulse) generated higher detectable changes in ΔL/L with CaBLAM when compared to GCaMP8s ΔF/F due to the low baseline signal of CaBLAM (**Fig. 3C**, Wilcoxon signed rank sum test, two tailed, Bonferroni correction for multiple comparisons, CaBLAM vs GCaMP8s at 1 FS: *P* = 1.1 × 10^-10^, 5 FS: *P* = 2.45 × 10^-21^, 60 FS: *P* = 2.03 × 10^-2^). The response kinetics of CaBLAM are considerably slower than GCaMP8s, as expected, since CaBLAM has not yet been optimized for fast responses (**Fig. 3D**, two-tailed Kolmogorov-Smirnov test, *P* = 7.1 × 10^-51^, *n* = 96 trials/20-38 GCaMP8s expressing neurons vs 570 trials/88-276 CaBLAM expressing neurons).

**Figure 3.**
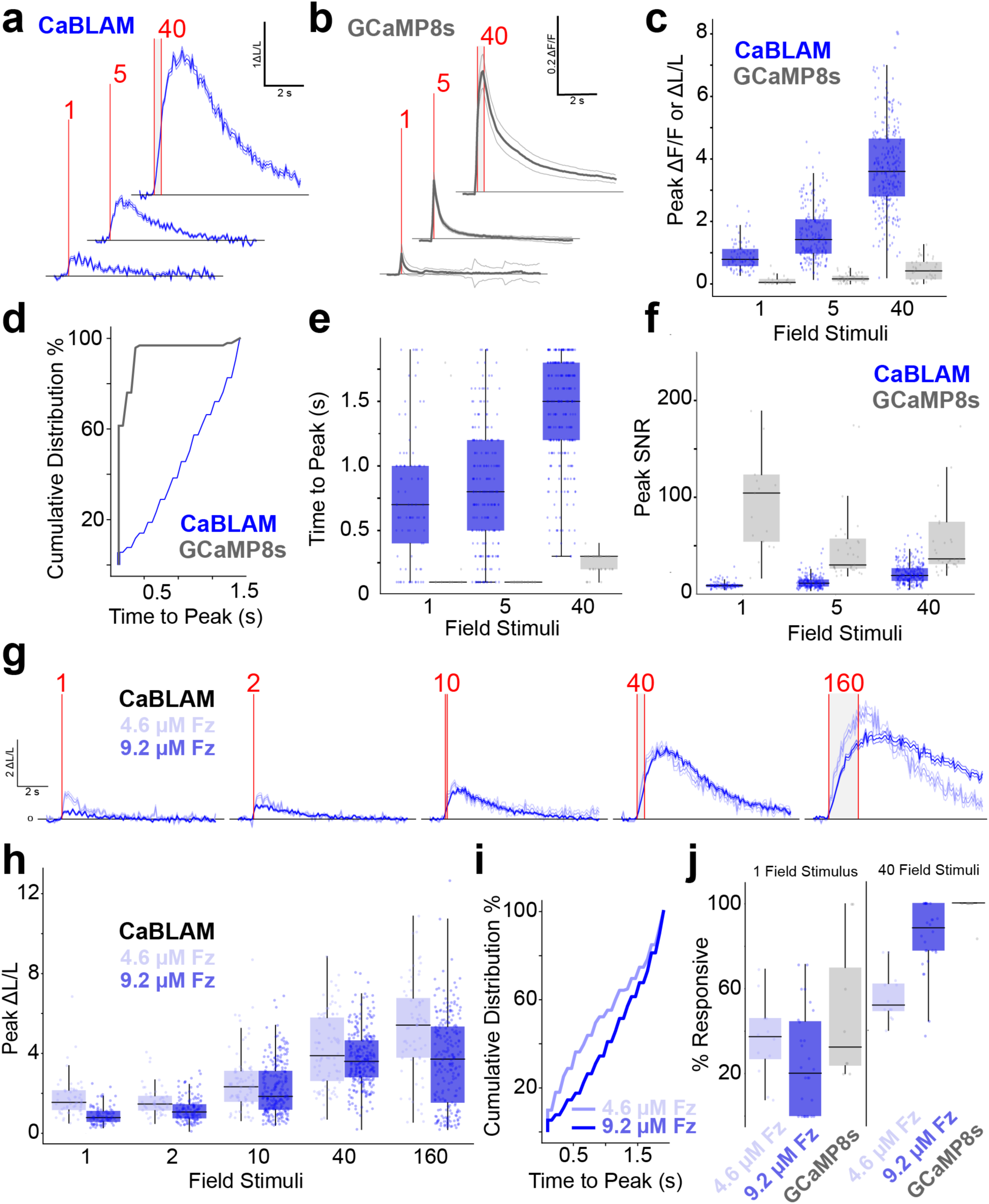
CaBLAM provides high-contrast reporting of stimulus-evoked neural activity in cultured neurons. **a,** Bioluminescence ΔL/L time-locked traces of CaBLAM Ca^2+^ responses to 1, 5, 40 pulses of 1ms field stimulations at 83 Hz (red dashed lines indicate field stimulation window). **b,** Same as panel **a**, but fluorescence ΔF/F time-locked traces for GCaMP8s. For panels **a** and **b**, data are shown as mean ± s.e.m. **c,** Peak stimulus evoked ΔF/F or ΔL/L across neurons across increasing field stimulations. **d,** Cumulative distribution of time to peak ΔF/F or ΔL/L responses. **e,** Time to peak ΔF/F or ΔL/L across neurons in response to increasing field stimulations. **f,** Peak signal-to-noise ratio (SNR) of ΔF/F or ΔL/L responses across neurons at increasing stimulation intensities. **g,** Overlaid bioluminescence ΔL/L time-locked traces of CaBLAM Ca^2+^ responses to 1, 2, 10, 40, 160 pulses of 1ms field stimulations at 83 Hz, at 4.6 μM or 9.2 μM Fz concentration. Red dashed lines indicate field stimulation window. Data are shown as mean ± s.e.m. **h,** Peak stimulus ΔL/L for CaBLAM across neurons elicited across increasing field stimulations. **i,** Cumulative distribution of time to ΔL/L responses between (includes statistical analysis). **j,** Proportion of neurons responding to 1 (left) and 40 (right) electrical field stimulations. All boxplots show the median, 25th and 75th percentiles (box edges), whiskers extending to the most extreme data points, and individual outliers plotted separately.

For both indicators, we measured the time to peak ΔL/L (CaBLAM) and ΔF/F (GCaMP8s) and observed distinct response timing. GCaMP8s reached peak responses around 100 ms (approximately 1 frame post-stimulation at our 10 Hz imaging rate) for 1 and 5 pulses, while the response extended to 300 ms (about 3 frames) for higher stimulation frequencies (**Fig. 3E**, Wilcoxon signed rank sum test, two tailed, Bonferroni correction for multiple comparisons, CaBLAM vs GCaMP8s at 1 FS: *P* = 1.93 × 10 ^-05^, 5 FS: *P* = 1.07 × 10 ^-16^, 60 FS: *P* = 1.24 × 10^−22^). In contrast, CaBLAM under constant Fz perfusion (9.2 μM final concentration), displayed median peak response times of approximately 700, 800, and 1500 ms post-stimulation for 1, 5, and 40 pulses, respectively. As these experiments did not include synaptic blockers, the true indicator kinetics are likely faster than measured. Next, we compared the peak signal to noise ratio (SNR) between CaBLAM and GCaMP8s. As we expected when imaging a monolayer of primary neurons in culture, GCaMP8s exhibited higher peak SNR in this scenario despite its lower contrast relative to CaBLAM (**Fig. 3F**, Wilcoxon signed rank sum test, two tailed, Bonferroni correction for multiple comparisons, CaBLAM vs GCaMP8s at 1 FS: *P* = 7.86×10⁻¹¹, 5 FS: *P* = 1.14×10⁻¹⁹, 40 FS: *P* = 1.41×10⁻¹⁴).

We then examined how varying Fz concentration influenced CaBLAM’s ability to report Ca^2+^ dynamics in this field stimulation benchmark. For analysis, we pooled all time-locked evoked responses across stimulations to analyze the average response and temporal kinetics of CaBLAM. Interestingly, neurons expressing CaBLAM imaged in a bath application of 4.6 μM Fz reported *larger* stimulus time-locked responses (ΔL/L) compared to those in 9.2 μM Fz at both lower and higher stimulation levels (**Fig. 3G, H**, Wilcoxon signed rank sum test, two tailed, Bonferroni correction for multiple comparisons, *P* = 6.63 × 10^−5^, 4.6 μM Fz: Mean= 3.24, Median= 2.49, SD=2.15, 95% CI= [3.01, 3.47], 9.2 μM Fz: Mean= 2.71, Median= 2.42, SD=1.85, 95% CI= [2.59, 2.83]), suggesting an improved ability to detect smaller changes in intracellular Ca^2+^ in at the lower Fz concentration. In parallel experiments using constant Fz perfusion (rather than bath application), the time to peak ΔL/L for CaBLAM after stimulation was lower with 4.6 µM Fz compared to 9.2 µM Fz, indicating a *faster* response at the lower Fz concentration (**Fig. 3I**, two-tailed Kolmogorov-Smirnov test, *P* = 5.61 × 10^-8^, *n* = 333 trials/148 neurons in 4.6 μM Fz vs 961 trials/322 neurons in 9.2 μM Fz). These findings are surprising, since reducing the luciferin concentration would typically decrease overall signal and make detection less efficient. We attribute this counterintuitive finding to saturation effects at substrate concentrations or near the *K*_m_ of the enzyme, as well as potential modulation of apparent Ca^2+^ affinity by substrate binding via unexpected allosteric effects.

We then proceeded to compare the *responsiveness* of neurons expressing CaBLAM under varying luciferin concentrations to those expressing GCaMP8s. There was no significant difference in the proportion of responsive neurons between GCaMP8s and CaBLAM, regardless of Fz concentration, in response to 1 ms electrical field stimulation (**left, Fig. 3J**, Kruskal Wallis test *χ*^2^(2) = 5.02, *P* = 0.081). This indicates that CaBLAM’s BL signal can reliably report single field stimulations, comparable to the state-of-the-art fluorescent indicator GCaMP8s. In contrast, delivery of 40 electrical field stimulations revealed a significant difference in the proportion of responsive neurons between CaBLAM-expressing neurons and those expressing GCaMP8s, with CaBLAM showing decreased responsivity at lower concentrations of Fz (**right, Fig. 3J**, Kruskal Wallis test: *χ*^2^(2) = 24.3, *P* = 5.23 × 10^-6^). Pairwise comparisons indicated that CaBLAM expressing neurons in the presence of 9.2 μM Fz had a significantly higher proportion of responsive neurons compared to 4.6 μM Fz (*P* = 4.79 × 10^-4^). The mean response rate was 98% for GCaMP8s, 84% for CaBLAM at 9.2 μM Fz, and 56% for CaBLAM at 4.6 μM Fz, suggesting that the 9.2 μM Fz concentration yields observed evoked responses similar to those of GCaMP8s (*P* = 0.109).

### Evaluation of additional CaBLAM substrates

The introduction of fluorofurimazine (FFz)^32,33^ and cephalofurimazine (CFz9, referred to hereafter as CFz)^34^ to the research community, shown to be superior *in vivo* substrates for NanoLuc-derived indicators, led us to investigate their performance with CaBLAM. We recorded BL from N2a cells transiently expressing CaBLAM over a 20 min time course after adding graded concentrations of CFz, FFz, or furimazine (Fz) and quantified the area under each curve (AUC) to generate dose–response relationships (**Supp. Fig. 7**). CFz and FFz responses were best described by a hybrid Hill + linear-decay model, yielding EC₅₀ values of 7.0 µM (*n_H_* = 0.86, R² = 0.97) and 8.6 µM (*n_H_* = 1.33, R² = 0.99), respectively, whereas Fz followed a simple Hill function with an EC₅₀ of 0.80 µM (*n_H_* = 1.10, R² = 0.96). High-speed EMCCD imaging of individual cells across representative substrate concentrations (0.01–1000 µM for CFz and FFz; 0.23–2.3 µM for Fz) confirmed that all three luciferins supported robust bioluminescence and large ionomycin-evoked signals; however, CFz and FFz produced markedly higher baseline brightness (L₀/F) and ionomycin-evoked contrast (maximum ΔL/L_0_) than Fz at their respective optimal concentrations (**Supp. Fig. 8**). These data establish CFz and FFz as effective alternatives to Fz for driving CaBLAM, with FFz providing the best overall combination of brightness and dynamic range in cultured cells.

### Comparison of CaBLAM and GCaMP6s *in vivo*

To characterize and compare the performance of CaBLAM to GCaMP6s *in vivo* we used an established model for sensor development^35^, neocortical interneurons labeled in the NDNF-Cre mouse line (**Fig. 4, Supp. Movie 3**). After selective viral expression and surgical preparation, we performed imaging either without external illumination (CaBLAM) or under 1-photon epifluorescent illumination (GCaMP6s). All mice were head-fixed and free to run on a wheel as tactile stimuli were delivered to their vibrissae **(Fig. 4A**). Imaging was performed through a cranial window and, in the CaBLAM expressing group, FFz was infused through a canula implanted at the edge of the imaging window. This approach allowed us to directly assess single-cell 1-photon BL and fluorescent signals evoked by tactile stimulation. All animals exhibited detectable activity in single interneurons (Total cell numbers: CaBLAM: Median = 42, IQR = 18; GCaMP6s: Median = 84, IQR = 74.25). Replicating recent findings, subsets of NDNF cells exhibited either positive (**Fig. 4E,G-I**) or negative responses to tactile stimuli^36^, and modulation by arousal state as indexed by their running speed (**Fig. 4K**; ^37^). The majority of responsive cells in both sensor groups were positively modulated by tactile stimuli (**Fig. 4F**, CaBLAM: 97% and GCaMP6s: 72% of all responsive cells), and as such we focused our analyses on these cells (See **Extended Data Fig. 1** for examples of negative responses).

**Figure 4.**
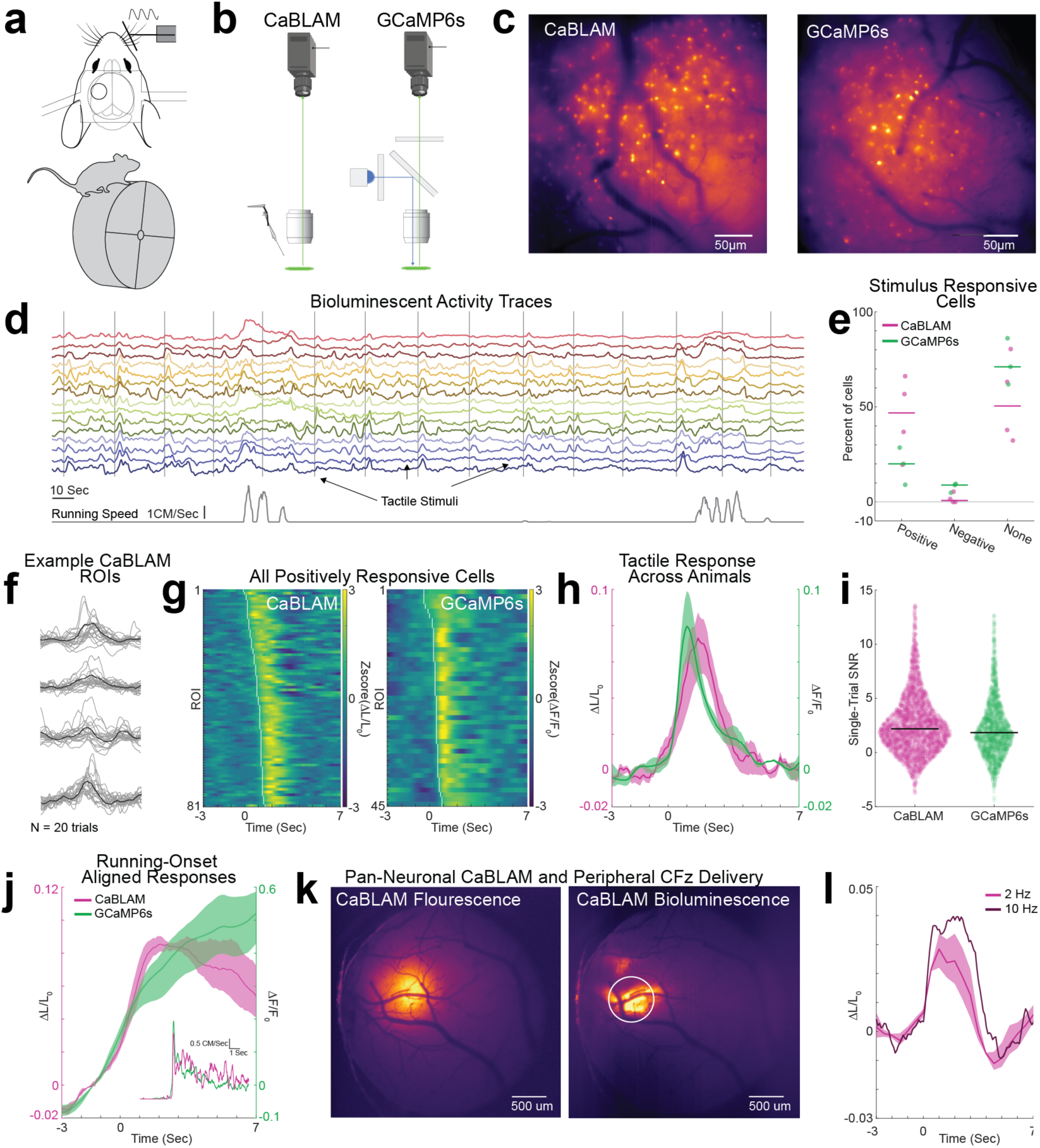
CaBLAM Shows SNR Comparable to GCaMP During *in vivo* Mammalian Imaging. **a,** The mouse imaging setup with location of the cranial window and head-fixation on a running wheel. **b,** The CaBLAM and GCaMP6s imaging configurations: CaBLAM was imaged in darkness in a light tight enclosure after administration of luciferin; and, GCaMP was imaged under epifluorescent illumination. **c,** Mean projection images of an NDNF-Cre CaBLAM and GCaMP6s FOV**. d,** CaBLAM activity traces from 15 neurons in one FOV. Vertical gray bars indicate onset of tactile stimuli, and the *bottom* trace is mouse running speed. **e,** Percentages of stimulus responsive cells across animals and sensor groups. Individual mice are represented by dots and group medians by horizontal bars. **f,** Example of 20 tactile stimulation trials for 4 cells from 1 FOV. Gray traces are individual trials, and black traces are the mean. **g,** Mean responses of all positively responsive cells for the NDNF CaBLAM (*left*) and NDNF GCaMP6s groups sorted in ascending order by latency of the signal at half maximal. **h,** Mean sensory response across mice in the two groups from positively responsive cells. Stimulation onset is at t = 0, indicated by the vertical dashed line. Opaque bars represent the jackknifed 95% CI of the means. Separate *left* and *right* y-axes for the CaBLAM and GCaMP signals, respectively. **i,** Single-trial SNR measurements across all positively responsive cells for the two sensor groups. Dots are individual trials and horizontal lines indicate the median. **j,** Mean running-onset waveforms across animals for the two sensor groups. Running onset is at t = 0, indicated by the vertical dashed line. *Inset*, the average running speed across mice time-locked to onset of running bouts. **k,** *Left*, epifluorescence image of a full cranial window expressing CaBLAM pan-neuronally. *Right*, the CaBLAM BL average projection in the same mouse after retro-orbital injection of CFz. **l,** Mean CaBLAM tactile response (*n* = 3 mice) after retro-orbital injections of CFz (sampled ROI shown by the *white circle* in **k**). Stimulation onset is at t = 0, indicated by the vertical dashed line. For 2Hz sampling rate (*Magenta*), opaque shading represents the jackknifed 95% CI of the mean. The *darker* trace shows an average tactile response from one animal imaged at 10 Hz.

The tactile evoked waveforms averaged across neurons and animals exhibited close correspondence between the two sensor groups and, although necessarily different measures, tactile evoked ΔF/F_0_ and ΔL/L_0_ were highly comparable in magnitude and overall temporal dynamics (**Fig. 4I**). We measured response onset latency by finding the half peak within the 0-2 s response window (**Fig. 4H**). With this metric, CaBLAM onset latencies occurred later than GCaMP6s onset latencies (Wilcoxon rank sum, *P* = 3.3 x 10^-8^, *z* = 5.52; CaBLAM: *n* = 82, Median = 1 s, IQR 0.7, GCaMP6s: *n* = 44, Median = 0.45 s, IQR = 0.25). This difference in response latency agrees with our *in vitro* data.

To compare the ability of each sensor to detect a signal when present, single trial SNR was computed across all cells as max 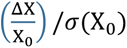, normalizing the maximum change from baseline 0 to 2 s post stimulus (numerator) by the standard deviation (SD) of the baseline period (denominator, -3 to 0 s). Single-trial CaBLAM SNRs were significantly greater than those of the GCaMP6s (Wilcoxon rank sum test, *P* = 1.89 x 10^-5^, z = 4.28; CaBLAM SNR: *n* = 3127, Median = 2.18, IQR = 3.22; GCaMP SNR: *n* = 1570, Median = 1.83, IQR = 3.20). To determine the components of the SNR that differ between the sensors, we ran two subsequent analyses comparing the single-trial peak values and single-trial baseline SD. We found that while signal peaks were typically larger in CaBLAM, this trend was not significant (*P* = 0.079, *z* = 1.76; CaBLAM Median = 0.096 ΔL/L_0_, IQR = 0.16; GCaMP6s Median = 0.088 ΔF/F_0_, IQR = 0.17). However, CaBLAM baseline was steadier, showing a significantly smaller SD (*P* = 1.23 x 10^-3^, *z* = -3.26; CaBLAM Median = 0.044 ΔL/L_0_, IQR = 0.049; GCaMP6s Median = 0.049 ΔF/F_0_, IQR = 0.048). The low background of BL signals provided their significantly better performance for signal detection.

We next sought to test that CaBLAM was effective in reporting sensory-driven dynamics with peripheral administration of luciferin, a common experimental design. We injected 200 μL of CFz retro-orbitally in mice expressing CaBLAM pan-neuronally (*n* = 3; CaBLAM expressed by injection of AAV 2/9 under control of the hSyn promoter). Imaging a wide field-of-view (3.23 mm across) at 2 Hz, we saw robust BL responses to tactile stimulation with rapid onset times. Calculated as the half-peak latency of the average response, all mice exhibited onset latencies at time 0, indicating that onset occurred in the 0.5 s window between stimulus onset and the first post-stimulus image. This rapid onset time precedes almost all local hemodynamic responses (e.g., dilation), a concern that has been shown to contribute to sensory responses in other BL GECIs (**Fig. 4L, M**, ^34^). Trial-to-trial SNR was significantly greater than 0 (one-sample Wilcoxon signed rank test, *P* = 2.64 x 10^-30^), if typically less than the single-cell NDNF response (Median = 1.83, IQR = 2.66).

To better define the rapid onset of this response, we subsequently imaged a mouse under the same conditions at 10 Hz and found a highly comparable sensory response with regards to shape and magnitude (**Fig. 4M**, dark maroon trace). At 10 Hz the trial-to-trial SNR of the animal’s sensory response was slightly higher than at 2 Hz (10 Hz SNR: Median = 2.91, IQR = 1.89 vs. 2 Hz SNR: Median = 2.36, IQR = 3.06) but not significantly different (Wilcoxon rank sum test, *P* = 0.52, z = 0.64). Additionally at 10 Hz the half-peak latency of the tactile response occurred at 0.20 s, further supporting the conclusion that the onset latencies of the 2 Hz imaged animals all occurred in the 0-0.5 s window post-stimulus. We also tested the efficacy of this administration route and concentration in an NDNF-Cre mouse expressing CaBLAM. At 0.5 Hz frame rate and under the same conditions used for direct neocortical infusions, BL signals were detectable from NDNF cells but proved too weak to capture Ca^2+^ fluctuations on a relevant time scale.

To test the viability of CaBLAM as a long duration sensor, FFz was applied directly to the exposed cortex of of a pan-neuronal CaBLAM mouse under anesthesia, and BL recorded at 10 Hz for > 5 hours. Imaging commenced < 1 minute after FFz addition, at which time the BL signal was already apparent. To evaluate the overall time course of BL intensity, as context for understanding the SNR of the evoked response, mean emission values from a circular ROI (**Extended Data Fig. 2A**, black circle) were binned at 5-minute intervals, reaching a peak at 25 minutes, slowly decreasing over the next 2 hours, then plateauing at around half the peak brightness in hours 3 to 5 (**Extended Data Fig. 2B**).

The tactile CaBLAM response, measured as ΔL/L_0,_ was stable across the recording (30 min bins, Median = 93, IQR = 10 trials/bin, **Extended Data Fig. 3A**). A Kruskal-Wallace test indicated a significant difference in the peak ΔL/L_0_ of single-trial responses across the 10 30-minute bins (*P* = 4.52 x 10^-6^; χ^2^ = 41.24, df = 9). Post-hoc comparisons (Tukey’s HSD) indicated that the waveform peaks within the first 30-minute bin were significantly smaller than the bins at later times in the range from 90 to 240 minutes (all *P*s < 0.02). Additionally, the peaks in the second 30-minute bin (30 to 60 minutes after FFz administration) were significantly smaller than those in the 180- to 210-minute bin (**Extended Data Fig. 3B**). In sum, the peak ΔL/L_0_ tactile responses within approximately the first hour after FFz administration were smaller than all later times, after which the peaks were statistically similar across the latter 4 hours. Interestingly, smaller contrasts (ΔL/L_0_) were observed during the time window of maximal BL emission (**Extended Data Fig. 2B**), suggesting that at high luciferin concentrations, the dynamic range of CaBLAM was reduced due to signal saturation. That said, the SNR was not significantly different across time bins (Kruskal-Wallace test, *P* = 0.82, χ^2^ = 5.11, df = 9).

This experiment was concluded while the BL signals and sensory-evoked response were still robust, >5 hours after one FFz application. This indicates that single, long-duration sessions are possible, and that the duration of CaBLAM imaging is likely limited primarily by the bioavailability of fresh luciferin. After the extended imaging session, in the same mouse, we were able to clearly image neural processes (**Extended Data Fig. 4**) at a 1 Hz imaging rate at 40x magnification. Given that these images were acquired 5 hours after FFz administration, at which point the signal had faded from its peak, indicating that imaging Ca^2+^ dynamics in subcellular compartments *in vivo* using CaBLAM for extended periods, such as across the time scale of biological learning, is feasible.

### CaBLAM-derived bioluminescent signals in astrocytes and neurons follow high-amplitude tail movements in head-embedded larval zebrafish

We generated a new transgenic line *Tg(UAS:CaBLAM)* that, together with existing driver lines, would let us express CaBLAM in astrocytes and neurons. We built a simple microscope to simultaneously measure bioluminescence using a photon-counting PMT and behavior using a machine vision camera. Larval zebrafish at 3-4 days post-fertilization (dpf) were immersed in either 1:100 or 1:1000 vivazine for 10 minutes before each session and showed bright basal bioluminescence relative to control animals (**Table 2**). Consequentially, slight changes in elevation or orientation relative to the PMT that accompanied swimming produced large changes in detected counts. We therefore embedded the head of the fish in agarose, leaving the tail free to move while allowing us to center the fish relative to the PMT. Even embedded, the PMT signal still showed detectable slight variations due to changes in tail pitch and roll. However, we noticed that after particularly large-amplitude movements, the bioluminescent signal persisted long after movement had ceased. We focused on quantifying these large-amplitude movements, as they allowed us to attribute changes in bioluminescence to changes in neural activity.

**Table 2.** Parameters for zebrafish CaBLAM experiments.

| **Genotype** | **Vivazine concentration** | **Average bioluminescence (counts / s)\*** | **Number of high-amplitude events\*** | **Experiment duration (min)\*** |
| --- | --- | --- | --- | --- |
| ***Is(nefma)*** | 1:100 | 16631<br>17818<br>13793 | 25<br>10<br>12 | 81<br>37<br>41 |
| ***Is(nefma)* sibling controls** | 1:100 | 3007<br>2393<br>2692 | 11<br>15<br>8 | 39<br>60<br>34 |
| ***Tg(GLAST)*** | 1:1000 | 113664<br>20347<br>15280 | 10<br>8<br>6 | 32<br>28<br>28 |
| ***Tg(hcrtr2)*** | 1:100 | 16525<br>22896<br>32134 | 16<br>95<br>38 | 29<br>103<br>45 |
| ***Tg(elavl3)*** | 1:1000 | 36712<br>87159<br>67205 | 57<br>22<br>5 | 125<br>41<br>57 |

\* Values shown are from three experimental runs with individual zebrafish larvae for each genotype.

We began by using a new driver line, *Tg(GLAST)* ^38^ to express CaBLAM in zebrafish astrocytes, (**Fig. 5A**). Small, short increases in bioluminescence were associated with smaller movements, and large, prolonged increases followed the largest movements (**Fig. 5B**). The video frames around the time of large movements (e.g. **Fig. 5C-E**) show that they are composed of strong, asymmetric, and uncoordinated tail flicks (**Supp. Movie 4**). Increases in bioluminescence lasted ∼10 s (**Fig. 5F**), and varied in amplitude (**Fig. 5G**), and all large movements we observed were accompanied with increases in bioluminescence. Our data suggests that large amplitude tail movements elicit CaBLAM-derived bioluminescence.

**Figure 5.**
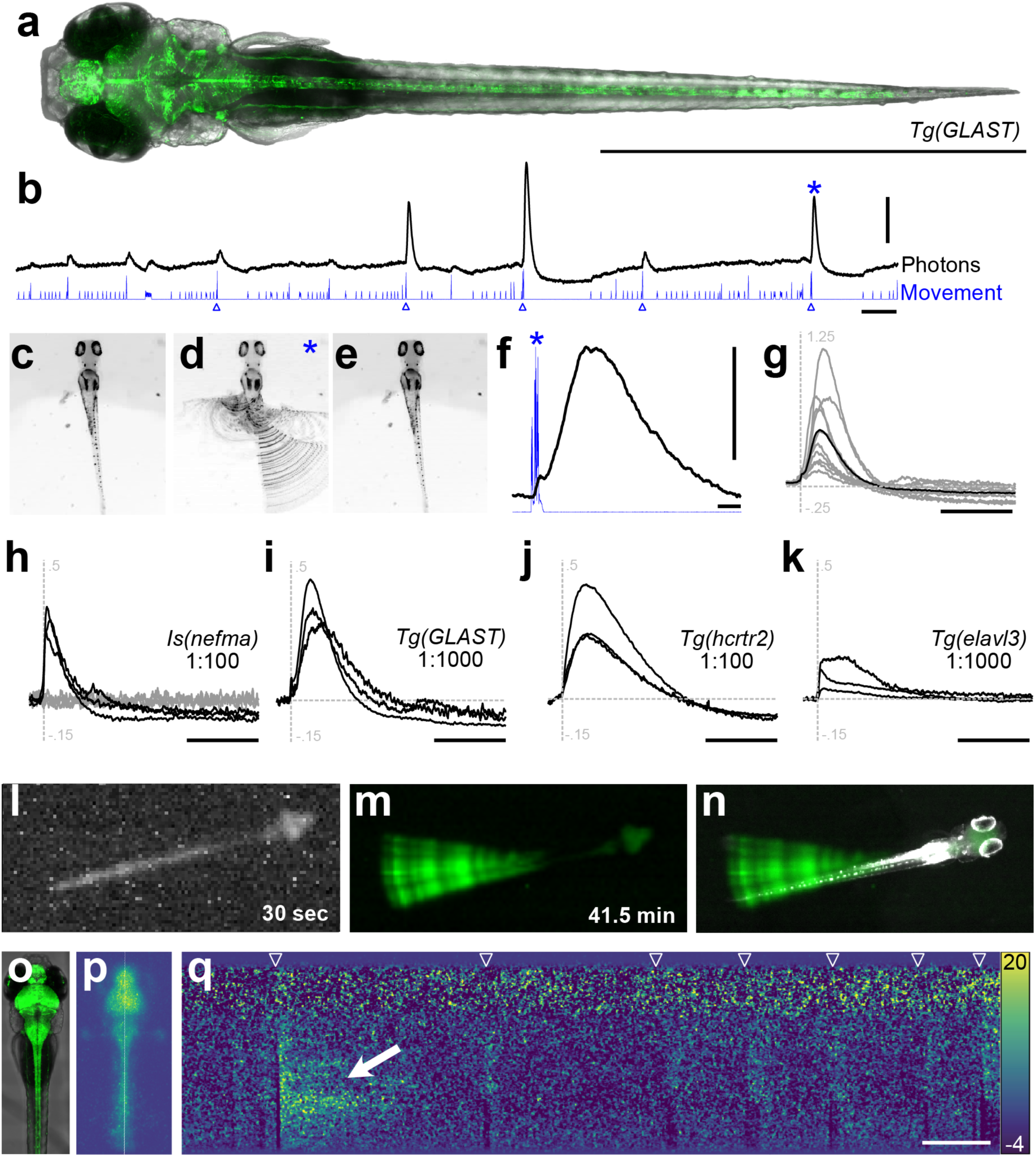
CaBLAM-derived bioluminescent signals in astrocytes and neurons follow high-amplitude tail movements in head-embedded larval zebrafish. **a,** Confocal image of a 5 dpf *Tg(GLAST)* larva showing expression of CaBLAM in astrocytes (green) and transmitted light (gray). Scale bar is 2 mm. **b,** A representative trace (80 s) of detected photons (black) and movement (blue). Blue triangles correspond to detected high-amplitude movements. Scale bars are 50,000 counts (vertical) and 10 s (horizontal). **c-e,** Summed frames from a 4 s video corresponding to a high-amplitude movement. **c** is the 200 ms before the movement, **d** is the 400 ms of movement, and **e** is the 3400 ms after the movement. Changes are only detectable during the movement, marked by strong uncoordinated and asymmetric tail flicks. **f,** A zoomed in view of photons and movement from c-e, marked by the asterisk in **b**. Counts begin during the movement and continue for seconds after the movement has ceased. Scale bars are 50,000 counts (vertical) and 1 s (horizontal). **g**, Normalized counts during detected high-amplitude movements (gray, n=10) over a full experiment. Black is the mean bioluminescent response. Dotted lines at t = 0 (vertical) and 0 counts (horizontal), vertical scale is -0.25 to 1.25, scale bar is 10 s. **h-k,** Normalized counts during high-amplitude movements for four different genotypes. Dotted lines at t = 0 (vertical) and 0 counts (horizontal), vertical scale is -0.15 to 0.5, scale bar is 10 s. **h**, *Is(nefma*) CaBLAM-expressing fish (black, n=3) and negative siblings (gray, n=3) with 1:100 vivazine substrate, 3-4 dpf, **i**, *Tg(GLAST)* at 1:1000, 3 dpf, n=3, **j**, *Tg(hcrtr2)* at 1:100, 3 dpf, n=3, **k**, *Tg(elavl3)* at a lower dose (1:1000), 3 dpf, n=3. l, A single 30 s exposure of a 5 dpf *Tg(elavl3)* fish at 1:100 vivazine. **m**, Pixel-wise standard deviation across a stack of 83 30 s exposures, **n,** A composite of the image in **n** and an infrared-illuminated image of the fish. **o,** Maximum intensity projection confocal stack of a 4 dpf *Tg(elavl3)* larva. **p**, Maximum projection of a 2 min intensified image of a 4 dpf *Tg(elavl3)* larva in 1:1000 vivazine. The vertical white line in the center of the image shows the location of the pixels used for analysis. **q,** 2 min kymograph of intensity changes over time in the pixels along the white line in **p**. White triangles indicate movements, the white arrow shows the elevated intensity that followed a large tail flip. Scale bar: 10 s

Different genotypes showed similar increases in bioluminescence following the largest movements, but with varied kinetics. Experiments lasted 28-125 minutes (**Table 2**). *Is(nefma)* labels descending midbrain/hindbrain premotor and spinal cord neurons^39^. CaBLAM-expressing *Is(nefma)* larvae (*n* = 3) displayed strong increases in bioluminescence with a fast rise followed by a decrease relative to baseline (**Fig. 5H**, black). Non-fluorescent siblings (*n* = 3) ran with identical 1:100 vivazine had baseline counts near the noise floor of the PMT (with IR illumination on) and showed no detectable fluctuations associated with the largest movements (**Fig. 5H**, gray). *Tg(GLAST)* larvae with CaBLAM expressed in astrocytes were similar to the example (**Fig. 5B-G**) with a slower rise time and decrease below baseline (**Fig. 5I**). *Tg(hcrtr2)* labels both neurons and large vacuolar cells in the notochord^40^ and shows a more prolonged response with a comparable decrease below baseline (**Fig. 5J**). Finally, *Tg(elavl3)* labels the majority of post-mitotic neurons^41^; we decreased the concentration of vivazine to 1:1000 and still observed changes in bioluminescence with a fast rise and minimal decrease below baseline (**Fig. 5K**). Average BL intensity, number of high-amplitude movements, and experiment duration for three head-fixed individuals from each genotype are detailed in **Table 2**. Taken together, our data support the inference that large amplitude movements are accompanied by prolonged Ca^2+^ flux in both neurons and astrocytes.

We performed two proof-of-principle experiments to explore imaging approaches that preserve spatial information about CaBLAM-derived bioluminescence. First, we used the machine vision camera (IMX174 sensor, SONY) in our microscope to image a 5 dpf *Tg(elavl3)* fish in 1:100 vivazine. A 30 s exposure, with 4×4 binning clearly reveals bioluminescence in the brain and spinal cord (**Fig. 5L**). We could remove the background noise by measuring the pixelwise standard deviation across repeated exposures (*n* = 83, 41.5 min), revealing the slight shifts in basal tail orientation across this particular experiment (**Fig. 5M**, **Supp. Movie 5**). Overlaying an IR-illuminated image of the larva (**Fig. 5N**) shows the alignment of fish and bioluminescence. We conclude that even a low-cost, uncooled, and un-intensified camera is sufficient to resolve CaBLAM-derived bioluminescence.

Second, we used an intensified high-speed camera (HiCAM FLUO, Lambert Instruments) to measure bioluminescence in a head-fixed 4 dpf *Tg(elavl3)* fish in 1:1000 vivazine (**Fig. 5O-P**). We could image a head-embedded larva at 20 fps, which was sufficient to resolve large and small amplitude tail movements (**Supp. Movie 6**). A kymograph of 2 min of imaging the midline of the fish was sufficient to resolve individual swims as the tail moved (**Fig. 5Q**). For one large movement, we observed a strong increase in intensity along the tail; qualitatively, caudal regions stayed bright for longer than rostral regions, consistent with a traveling Ca^2+^wave. Taken together, our data support the conclusion that CaBLAM-derived bioluminescence can be used to monitor Ca^2+^flux in neurons and astrocytes in larval zebrafish. Here, we use CaBLAM to reveal intense and prolonged increases in Ca^2+^flux along the tail following large-amplitude tail movements in head-fixed fish.

## Discussion

Here, we described the development of a powerful new Ca^2+^ indicator, CaBLAM. This BL GECI provides much higher resolution in cultured systems, including the identification of single action potential dynamics. *In vivo*, CaBLAM provides single neuron precision at high frame rates, allowing measurement of differentiated tuning and evolving dynamics in multiple cells with distinct tuning. These benefits were found in multiple leading model systems (zebrafish motor behavior and mouse vibrissa sensory processing).

With these new advances, CaBLAM is well-positioned to enable activity monitoring across a breadth of new applications. The ability to record well-resolved signals from many distinct neurons across hours in head-posted mice enables robust imaging without the photodamage and bleaching endemic to 1-photon fluorescence imaging. Further, studies of the neural dynamics supporting complex behavior are being conducted in free behavioral settings, that better match the context the vertebrate brain evolved to function within. Tracking activity in these contexts necessitates indicators that do not require fixing the position of the experimental subject, and minimize the additional implanted hardware required. To capture truly naturalistic behavior, environmental illumination is ideally kept low. Unlike fluorescent tools, BL GECIs do not require input of photons. As such, the complexity, weight and size of implanted hardware for imaging in freely moving preparations (e.g., GRIN lens-based or fiber photonic) can be cut approximately in half^42^ by using a BL GECI in place of a fluorescent GECI. Further, if net activity levels in a given target are the sought after metric, a BL GECI enables dynamic imaging *without any implanted systems*, as in our zebrafish studies and, in the future, mammalian model systems. Future studies using color-shifted variants of CaBLAM could also allow completely non-invasive imaging of the output of multiple brain areas and/or organ systems during such behavior.

While the first known GECIs were, in fact, naturally-occurring BL proteins^10–12^, fluorescent small molecule dyes and FP-based GECIs are currently the most popular tools for reporting Ca^2+^ activity in living systems. The primary limitation of BL probes relative to fluorescent probes has been their lower rate of photons emitted per indicator molecule at saturating Ca^2+^ concentrations. This peak emission intensity can be increased linearly over several orders of magnitude for fluorescent probes simply by increasing the intensity of the excitation light that is delivered to the sample, but the maximum intensity is limited by enzyme kinetics in BL probes. At the same time, all fluorescent probes undergo photobleaching during imaging which worsens as excitation intensity increases. In intact tissues, autofluorescence from endogenous molecules limits detection of weak fluorescent signals, supplying excitation photons becomes a major technical challenge, and confounding effects such as heating from intense excitation are of major concern. BL probes hold promise for circumventing these and other limitations of fluorescence, especially when used in deeper tissues.

While the signal-to-noise ratio advantage afforded by elimination of autofluorescent background partially compensates for the low light output of luciferases in imaging, required exposure times have historically been in the tens of seconds, far from what is necessary to observe meaningful Ca^2+^ dynamics and other fast cellular processes. Imaging changes in intracellular Ca^2+^ at high speeds is possible only when the contrast of an indicator is well above the electronic noise in the camera. BL GECIs derived from GeNL imaged at up to 60 Hz in iPS-derived cardiomyocytes^17^, which display very large changes in cytosolic Ca^2+^ concentration relative to most cells, displayed a dynamic range of less than 2-fold despite *in vitro* characterization suggesting much higher contrast. We observe that in HeLa cells displaying smaller swings in Ca^2+^ concentration when responding to histamine stimulation, the high baseline luminescence of the GeNL (Ca^2+^) sensor family limits the achievable dynamic range (**Fig. 1C**). In neurons, CaBLAM reliably reports responses to single action potential stimulation like GCaMP8s when imaged at 10 Hz and can be imaged indefinitely without photobleaching or phototoxicity when provided a steady supply of luciferin substrate. Most importantly, CaBLAM enables imaging of Ca^2+^ flux in awake mice at single neuron levels with superior SNR to GCaMP6s under 1-photon illumination. Future improvements to CaBLAM’s BL brightness are expected to further improve SNR and increase practically achievable frame rates and imaging depths, while future improvements in substrate bioavailability, blood-brain barrier permeability, and stability will continue to simplify substrate delivery and allow fully non-invasive free-behavior experiments.

Phototoxicity and photobleaching are distinct but interrelated phenomena. Exposing cells to low levels of blue light for periods as short as 4 minutes has been shown to alter their normal physiology^43^. In whole organism imaging, light exposure at typical FL GECI levels can even alter the typical development of non-neural structures, such as bone^44^. Photobleaching is typically approached as a limitation of the sensor that should be minimized for practical purposes, but the presence of photobleaching necessarily indicates the presence of phototoxic cellular damage, while the presence of photobleaching is not a prerequisite for phototoxicity^43,45^. The nature and extent of alterations in cell physiology caused by the illumination source in FL GECI imaging remains minimally documented, particularly in the context of *in vivo* imaging. Experiments that assess functional changes across several imaging sessions, such as those investigating learning and memory, may be particularly susceptible to these issues since slow changes in cellular activity due to the actual mechanisms of interest are essentially indistinguishable from changes induced by phototoxicity and photobleaching. *In vivo* fluorescent photobleaching is typically assessed along periods in the tens of minutes, thus longer duration FL GECI imaging (e.g., > 30 min), investigating changes in cellular dynamics on similar timescales are necessarily accompanied by phototoxicity. We demonstrate *in vivo* that imaging across at least 5 hours is possible using CaBLAM with stable SNR and very slow degradation in intensity, something that would currently be far beyond what is achievable with any fluorescent GECI.

Translucence makes fish species like *Danio* and *Danionella* particularly attractive models for BL imaging of Ca^2+^ flux in neurons and glia, but technical challenges have hindered progress. Earlier work established a first-generation reporter, Aequorin-GFP^46^ in zebrafish^12,47^, but, in contrast to cardiology^48^ and cancer biology^49^, BL has not been adopted in neuroscience, likely due to the strikingly low flux and challenges of dealing with off-target (i.e. muscle) expression^50^. In our experiments, we routinely saw 2-3 orders of magnitude more flux than the 400-3000 photons/s in rigorously screened GFP-Aequorin lines^47^. While the lines tested here likely drive broader expression, and we cannot rule out the possibility of low levels of muscle expression, we nonetheless see both strong BL that persists long past the movement and kinetics that are specific to particular transgenic driver lines. Thus, we view CaBLAM as an enabling tool for future experiments to monitor activity in genetically targeted populations of neurons and glia in freely behaving fish over long time periods.

Biochemical indicators using BL as their output modality hold the promise of enabling superficial or non-invasive deep imaging of these events than is possible with fluorescent sensors. CaBLAM will facilitate this type of imaging because of its low baseline signal, preventing a washout of true signal from scattered out-of-focus baseline emission that is common for both fluorescent and BL GECIs in thick tissues. Another motivation for BL indicators is their potential to be partnered with optogenetic tools, for example to allow simultaneous monitoring of neural activity and light-controlled neural activation, with the lack of fluorescence excitation allowing maximal flexibility in the choice of optogenetic channel. We also plan to use CaBLAM to *drive* optogenetic elements such as channelrhodopsins and light-sensitive transcription factors, making them response to changes in intracellular Ca^2+^, building on previous work fusing bioluminescence and optogenetics^51–57^.

Structure-guided engineering and directed evolution can generate large improvements in luciferase solubility and toxicity while maintaining high catalytic activity, and these highly active and soluble luciferases can be used to engineer biochemical indicators with large signal changes in cells. The scaffold and design principles we have identified will be useful in the design and optimization of bright, high-contrast BL sensors for ***other biochemical activities*** as well^58^. In the future, development of red-emitting SSLuc and CaBLAM variants, either through improved red-emitting FRET acceptors (such as optimized long Stokes-shift FPs) or engineering of high-activity, red-emitting substrates, will further enhance the deep-tissue imaging potential of these probes. Optimization of CaBLAM kinetics for detection of fast spiking events is another clear target for future development. Continuing efforts also focus on further improving the light production rate of this and other classes of luciferase. Looking forward, if *K*_m_ values are maintained at ∼10 μM, a diffusion-limited luciferase (i.e., *k*_cat_/*K*_m_ = 10^8^-10^9^ M^-1^s^-1^) should be capable of generating 10^3^-10^4^ photons per second per molecule, which leads us to hope that there is ample room left for future improvements to these enzymes. Given our current observations, efforts to engineer stable caged substrates with oral bioavailability and to increase the delivery rate of these engineered luciferins to the cytoplasm of target cells will be central to realizing the full potential of BL imaging probes.

## Materials and Methods

### General molecular biology

*E. coli* strain NEB 10-beta (New England Biolabs) was used for all cloning, library screening, and recombinant protein expression. PCRs were performed using Phusion DNA polymerase (New England Biolabs) and the supplied ‘GC buffer’ under manufacturer-recommended conditions. DNA oligonucleotide primers were purchased from IDT. Restriction enzymes were purchased from New England Biolabs and restriction digests followed the manufacturer’s protocols. All plasmids were constructed using Gibson assembly^59^ of PCR or restriction digest derived linear DNA fragments as previously described^23,60^. Unless otherwise noted below, chemicals were purchased from Sigma-Aldrich.

### Construction of *E. coli* expression plasmids and libraries

All *E. coli* expression plasmids were constructed using the pNCST backbone as previously described^23,60^. A linear DNA fragment of the pNCST vector was generated by PCR (primers, **Supp. Data File 2**). Designed luciferase and novel GECI coding sequences were initially prepared as synthetic genes (IDT), as were many designed variants during directed evolution in cases where site-directed mutagenesis would be impractical. Mammalian expression plasmids encoding GeNL(Ca^2+^)_480^17^ (plasmid #85205) and CaMBI^18^ (plasmid #124094) were obtained from AddGene and amplified by PCR (primers, **Supp. Data File 2**) for insertion into the pNCST expression vector. Error-prone mutagenesis was performed using the GeneMorph II kit (Agilent Technologies) as previously described^23^ with a target mutation rate of ∼5 base changes per kilobase of amplified product. Site-directed libraries were constructed by Gibson assembly of PCR fragments amplified to introduce mixed-base codons as previously described^23^. Selected clones from library screening were sequenced using standard Sanger sequencing (Eton Biosciences). Libraries were introduced to *E. coli* via electroporation with plating at densities between 10^3^ and 10^5^ colonies per 10cm petri dish of LB/Agar supplemented with 100 μg/mL carbenicillin (Fisher BioReagents) and incubated overnight at 37 °C. All plasmid sequences are provided in the **Supplementary Sequences** text file.

### Luciferin substrates

All substrates and stock solutions were stored at -80 °C. Stock solutions of native coelenterazine (CTZ, NanoLight, cat# 303) were prepared by dissolving in NanoFuel^®^ Solvent (NanoLight, cat# 399) to a final concentration of 10 mg/mL (23.6 mM). Furimazine (Fz) stock solution is a component of the Nano-Glo® Luciferase Assay System (Promega, cat# N1110 or N1120). Because Promega does not disclose the Fz concentration in this stock solution, we measured absorbance spectra of samples of the stock solution diluted in methanol, then calculated the concentration of the stock solution using the extinction coefficient of Fz (21,000 M^-1^cm^-1^ at 254 nm) from published patent application US2020/0109146A1, giving a concentration of 4.6 mM for the Promega stock solution. From this, we calculated the final Fz concentrations from each dilution ratio used (typically a 1:500 or 1:1000 dilution of the Fz stock solution for live imaging and 1:100 for *in vitro* assays) and provide this value in the text and figures.

Water soluble h-coelenterazine (hCTZ, Nanolight, cat# 3011) was dissolved in sterile saline (4.55 μg/μL). Nano-Glo® Fluorofurimazine *In Vivo* Substrate (FFz, Promega, cat# N4110) was dissolved in 525 μL of phosphate buffered saline per vial (4.6 μmol/525 μL). Nano-Glo® Cephalofurimazine *In Vivo* Brain Substrate (CFz9, Promega, cat# CS3553A01) was dissolved in 1 mL of 0.2 M Tris buffer (pH 8.0) per vial (4.2 μmol/1 mL). As needed for FFz and CFz, the lyophilized solids were broken up into aliquots and stored at – 80 °C; for use they were dissolved at the concentrations detailed above. For all live imaging experiments, luciferase substrates were dissolved just before use.

The caged Fz derivative Nano-Glo® Vivazine Live Cell Substrate (Promega, cat# N2580) was diluted 1:100 or 1:1000 into aquarium water immediately prior to pre-incubation of zebrafish larvae (described below). We did not attempt to measure the stock concentration of vivazine, and report concentrations as dilution factors.

### Luciferase library screening

Error-prone mutagenesis and site-directed libraries of luciferases were screened by spraying a sterile solution of 50 μM CTZ or 46 μM Fz (1:100) in 50 mM Tris-HCl (pH 7.5) onto individual plates using a small atomizer. For ‘manual’ screening, the entire screening process was performed in complete darkness or very dim far-red LED light, with care taken to dark-adapt vision prior to starting. Sprayed plates were examined by eye, and the brightest colonies were immediately marked. For image-based screening, plates were placed immediately into a light-tight chamber and imaged with a PRIME 95b sCMOS camera (Photometrics), followed by identification of the brightest colonies using ImageJ and manual marking on the plates. Marked colonies were grown individually in 5 mL 2xYT medium supplemented with 100 μg/mL carbenicillin with shaking at 250 rpm at 37 °C overnight.

### Protein Expression, Purification, and *in vitro* experiments

The pNCST plasmid is useful for protein library screening and expression because it does not require an inducer in most strains of *E. coli*. For characterization of selected luciferase and GECI clones, 1ml of each overnight liquid culture was pelleted by centrifugation and the supernatant was discarded. The bacteria were then resuspended once in ultrapure water to remove residual culture medium and pelleted again by centrifugation, discarding the supernatant. The washed pellet was then resuspended in 400 μL B-PER (Thermo) reagent and incubated at room temperature for 20 minutes with gentle rocking to extract soluble proteins, followed by a final centrifugation to pellet insoluble components and yield a clear lysate. Clarified B-PER lysates were used directly for characterization experiments in most cases, since we observed rapid degradation of some proteins when subjected to additional purification procedures. In all cases, proteins prepared as clarified B-PER lysates were assayed within 6 hours of extraction.

For experiments using purified recombinant protein, B-PER lysates were bound to equilibrated cobalt beads as previously described^23,60^, washed, and eluted in 50 mM Tris-HCl (pH 7.5), 150 mM NaCl, 500 mM imidazole, then buffer-exchanged into 50 mM Tris-HCl (pH 7.5), 150 mM NaCl using Zeba desalting columns (Pierce). Purified proteins were stored at 4 °C.

For brightness comparisons *in vitro*, B-PER lysates of mNG-fused luciferase clones were diluted in 50 mM Tris-HCl (pH 7.5), 150 mM NaCl to a peak mNG absorbance value < 0.05 and 200 μL samples were loaded into a clear-bottom black 96 well plate (Corning) along with B-PER lysates from one or more controls (e.g., GeNL). Fluorescence emission spectra were recorded for each sample with excitation at 480 nm (5 nm bandwidth) using a Tecan M1000Pro Infinite multimode plate reader. To normalize enzyme concentrations based on the fluorescence, a dilution factor was calculated for each well based on the peak fluorescence emission. Samples were then diluted into new wells to a total volume of 50-100 μL and were measured again to verify that fluorescence intensities varied by ≤ 5% over all wells. The plate was then transferred to a ClarioStar multimode plate reader (BMG) to measure luminescence kinetics. Luciferin solutions at 50 μM (CTZ) or 46 μM (Fz, 1:100) were prepared in the same Tris buffer as described above and injected in volumes of 100-150 μL individually into wells with concurrent measurement of luminescence intensity for a total measurement time of 50 s per well. These sample and injection volumes were chosen to reduce variability in measurements due to uneven mixing, which can occur if the injection volume is smaller than the sample volume. Peak and steady-state intensities were then normalized by dividing by the previously measured fluorescence of each well to account for remaining small variations in concentration between wells, and these normalized values were used for brightness comparisons between clones.

### Ca^2+^ titrations in vitro

B-PER lysates of mNG-fused GECIs were diluted ∼100-fold in 30 mM 3-(N-morpholino)propanesulfonic acid (MOPS) (pH 7.2), 100 mM KCl, 1 mM MgCl_2_ and then dispensed into 3 rows (36 wells) of a 96-well plate, 5 μL per well. EGTA-buffered Ca^2+^ solutions with free Ca^2+^ concentrations ranging between 0 and 39 μM were then prepared from commercial stock solutions (Invitrogen cat# C3008MP), also in 30 mM MOPS (pH 7.2), 100 mM KCl, and supplemented with MgCl_2_ to a final concentration of 1 mM. The presence of Mg^2+^ provides more physiologically relevant conditions for this assay, but slightly alters free Ca^2+^ concentrations in EGTA-based buffers, so we calculated final free Ca^2+^ for each condition using parameters determined by Tsien & Pozzan^61^ and validated these calculations using the small-molecule Ca^2+^ dye fluo-4 (ThermoFisher cat# F14200). To obtain a fuller titration curve when titrating the low-affinity CaBLAM_294W variant, we prepared an additional unbuffered solution of 100 μM CaCl_2_ in 30 mM MOPS (pH 7.2), 100 mM KCl, 1 mM MgCl_2_.

For the titration, 100 μL of Ca^2+^ buffer solution was added to each well containing sample to produce three or more identical rows of the same concentration series. A solution of 46 μM Fz (1:100) in the same Mg^2+^-containing MOPS buffer was prepared and injected in volumes of 150 μL individually into wells with concurrent measurement of luminescence intensity for a total time of 20 s per well. The area-under-the-curve (AUC) luminescence values were calculated for each kinetic curve and normalized per replicate series. Custom python code was then used to fit titration curves using a three-parameter Hill equation: *y* = *y_min_* + (1 - *y_min_*) / (1 + (EC_50_/*x*)^*n_H_*), where y_min_ is the minimum normalized emission, EC_50_ is the Ca^2+^ concentration at half-maximal response, and *n_H_* is the Hill coefficient. Confidence intervals were calculated using the full covariance matrix to account for parameter correlations. Contrast ratios were calculated by taking the reciprocal of the mean of all data points at Ca^2+^ of < 10 nM, after normalization, for all sensors except CaBLAM_294W, for which we used all points at Ca^2+^ of < 50 nM. Plots were generated using custom Python scripts with matplotlib. Marker transparency was automatically adjusted based on local density to improve visibility of overlapping data points. Color schemes were selected to maximize contrast and accessibility. All plots use logarithmic Ca^2+^ concentration scales and linear normalized emission scales (0-1).

### Construction of mammalian expression plasmids

Mammalian expression constructs were mainly generated by Gibson assembly of PCR-amplified coding sequences (primers, **Supp. Data File 2**) into restriction-digested pC1 (CMV promoter, modified from pEGFP-C1, Clontech) or pCAG (CMV enhancer fused to chicken beta-actin promoter) vector fragments. When vector fragment preparation by restriction digest was impractical, we PCR-amplified vector fragments instead (primers, **Supp. Data File 2**). We observed qualitatively less variable and longer-lasting expression using the pCAG vector in most cases. Both vectors were suitable for all constructs tested. Mammalian expression plasmids encoding GeNL(Ca^2+^)_480^17^ (plasmid #85205) and CaMBI^18^ (plasmid #124094) were obtained from AddGene. The plasmid pCAG_EGFP_GPI (plasmid #32601) was obtained from AddGene and digested with *Sac*I and *Xho*I to generate a vector fragment for insertion of GeNL and GeNL_SS for GPI-anchored extracellular display. All plasmid sequences are provided in the **Supplementary Sequences** text file.

### Cell line culture, Transfection, and Imaging

HeLa and U2OS cell lines were purchased from ATCC. Cells were maintained under standard culture conditions with incubation at 37 °C and 5% CO2. Growth medium for HeLa cells was high-glucose DMEM (Gibco) supplemented with 10% fetal bovine serum (FBS) (Gibco), and for U2OS was McCoy’s 5a (Gibco) with 10% FBS. For transfection and imaging, cells were plated at a density of 1-2x10^5 cells/mL on coverslip-bottom 35mm dishes (Mattek) or 13mm round coverslips (BioscienceTools) placed in 6-well culture plates and incubated overnight. The following day, cells were transfected using polyethyleneimine (PEI) as previously described^60^. Cells were imaged 24-48 hours post-transfection.

Prior to imaging non-sensor luciferase constructs and fusions, cells were gently rinsed with fresh medium, leaving 500 μL of medium in each dish for Mattek dishes. For cells plated on coverslips, slips were transferred to a low-profile coverslip chamber (BioscienceTools) with a silicone gasket and overlaid with 500 μL medium. In either case, cells were transferred to the stage top environmental chamber and allowed to equilibrate for at least 5 minutes prior to imaging.

Image acquisition was performed in a stage-top environmental enclosure (37 °C, 5% CO2; Okolab) on a Nikon Ti-E microscope and an Andor iXon Ultra 888 EMCCD camera. Cells expressing BL constructs were imaged with a Plan Apo λ 20x Ph2 DM 0.75NA objective (Nikon). For reference fluorescent images in constructs containing mNeonGreen, green fluorescence was excited with a Spectra X LED source (Lumencor) using the 475/28nm channel and a FF495-Di03 dichroic (Semrock) and emission was selected with a FF01-520/35 filter (Semrock). The camera was set to 30 MHz at 16-bit horizonal readout rate, 2x pre-amplifier gain, and an electron multiplication (EM) gain setting of 3 for fluorescence acquisition. For BL signal recording, the illumination source was turned off and emission was collected either with no filter to maximize light collection efficiency or with the FF01-520/35 filter to minimize background light leakage. The camera was set to 30 MHz at 16-bit horizonal readout rate, 2x pre-amplifier gain, and EM gain of 300, with either 1×1 or 2×2 binning. Exposure times were set to between 50 and 500 ms for bioluminescence time series and between 500 ms and 1 s for single images.

For Ca^2+^ indicator time series imaging, cells were only plated in Mattek dishes and were not rinsed, a handling method we found prevented induction of extraneous Ca^2+^ signals in the cytosol. Medium was carefully removed to leave 500 μL volume remaining in each dish, and 500 μL of medium from the dish was reserved and used to dilute the Fz substrate to eliminate small changes in ionic strength and osmolarity that can arise due to evaporation during culture. Baseline images were collected for at least 30 s in the dark prior to luciferin addition. Fz diluted in culture medium was then injected to a final concentration of 4.6 μM (1:1000) and the BL signal was recorded for 60s. Next, 50 μL of concentrated ionomycin was injected to give a final concentration of 20 μM and images were recorded for an additional 60-120 s for determination of the maximal contrast in cells. The concentration of Ca^2+^ in the culture medium was ∼1 mM (but likely buffered to some degree by other medium components) and therefore expected to produce a “maximal” physiological Ca^2+^ concentration in the cytosol in the presence of ionomycin without additional supplementation. For imaging induced Ca^2+^ oscillations, ionomycin injection was replaced by injection of 50 μL of concentrated L-histamine to give a final concentration of 50 μM, and images were recorded for an additional 10-20 min.

### Primary Rat Cortical Neuron Culture, Transfection, and Ca^2+^ Imaging

Animal procedures were approved by the Institutional Animal Care and Use Committee of UC San Diego. Cortices were dissected out from P2 Sprague-Dawley rats and neurons were dissociated using papain as previously reported^62^. Transfection of neuronal cells with pCAG-CaBLAM and pCAG-GeNL(Ca^2+^)_480 was done by electroporation using an Amaxa Nucleofection Device (Lonza) at day-in-vitro 0 (DIV0). Neurons were cultured on poly-D-lysine (PDL) coated 35 mm coverslip-bottom dishes (Mattek) in Neurobasal A medium (Life Technologies) supplemented with 1X B27 Supplements (Life Technologies), 2 mM GlutaMAX (Life Technologies), 20 U/mL penicillin, and 50 mg/mL streptomycin (Life Technologies) for 2-3 weeks prior to imaging, refreshing half medium every 2-3 days.

For Ca^2+^ imaging with KCl-mediated depolarization, Mattek dishes were treated similarly to cell lines, with removal of all but 500 μL medium, equilibration in the stage-top incubation chamber on the microscope, and imaging following the same basic protocol as described above for ionomycin and L-histamine experiments. Dark images were recorded for ∼30s, followed by injection of Fz diluted in 1000 μL medium at a final concentration of 4.6 μM (1:1000) and continuous imaging for 60 s. Finally, all neurons were rapidly depolarized by injection of 500 μL of medium supplemented with 120 mM KCl to give a final concentration of 30 mM KCl and images were collected for an additional 30-60 s.

### Primary Rat Hippocampal Neuron Culture, AAV Transduction, and Ca^2+^ imaging

Primary E18 rat hippocampal neurons were prepared from tissue shipped from BrainBits (Transnetyx) following the vendor’s protocol. Neurons were seeded on PDL-coated 18 mm glass coverslips (Neuvitro) in 12-well tissue culture plates (1 x 10^5^ neurons per well) and grown in Gibco Neurobasal Media supplemented with 2% B27, 0.1% gentamycin and 1% GlutaMAX (all from Invitrogen). The next day, day-in-vitro 1 (DIV1), neurons were transduced with AAV9-Syn (5 x 10^9^ gc per well) encoding jGCaMP8s (AddGene 162374) or CaBLAM (in-house prep). AAV9-Syn-CaBLAM was generated by triple lipofection of HEK293-FT cells and harvesting viral particles as previously described^63^. Neurons were imaged between DIV 20 – 30.

For Ca^2+^ imaging with electrical stimulation, cover slips were washed three times and imaged in artificial cerebrospinal fluid, containing: 121 mM NaCl, 1.25 mM NaH_2_PO_4_, 26 mM NaHCO_3_, 2.8 mM KCl, 15 mM d(+)- glucose, 2 mM CaCl_2_, 2 mM MgCl_2_ (maintained at ∼37 °C, pH 7.3-7.4, continuously bubbled with 95% O_2_/5% CO_2_ vol/vol). All imaging was conducted in the RC-49MFSH heated perfusion chamber where the temperature was continuously monitored (Warner Instruments). Only cultures where neurons were evenly distributed across the entire coverslip. In addition, only coverslips with minimal neuronal clumping/astrocytic growth were included for Ca^2+^ imaging. A Teensy 3.2 microcontroller was utilized to synchronize the camera frame number with electrical stimulation using BNC cables connected to EMCCD camera and the SIU-102 stimulation isolation unit (Waner Instruments). Custom written scripts were developed in order to generate analog signals for precise control current stimulation as previously described^64^. Briefly, current field stimulation with a 1 ms pulse width at 40 mA and 83 Hz was utilized. The length of time for each stimulation period was varied (12, 36, 60, 120, 240, 960, 1920 ms) in order to achieve the corresponding number of action potentials (1, 3, 5, 10, 20, 80, 160 action potentials).

For fluorescent and BL Ca^2+^ imaging, a 20x objective lens (0.75 NA) and iXon Ultra 888 EMCCD (Andor Technology) camera were used. Furimazine (Fz) (Promega, cat# N1120) was diluted at 1:1000 or 1:500 in bubbled ACSF to give final concentrations of 4.6 μM or 9.2 μM, both of which provided sufficient BL signal for imaging at 10 Hz. Image acquisition parameters were as follows: 0.09 s exposure time, 4.33 μs vertical pixel shift, normal vertical clock voltage amplitude, 10 MHz at 16-bit horizonal readout rate, 1x pre-amplifier gain, 2×2 binning. The EM Gain was set to 300 for BL image acquisition and it was not enabled during fluorescent imaging. ACSF was continuously perfused in the heated chamber prior to imaging and during bright field and fluorescence imaging to determine a field of view before beginning BL Ca^2+^ imaging. During BL Ca^2+^ imaging, fresh ACSF was utilized to dilute the Fz and kept at 37 °C in a water bath. BL imaging included a 60 s initial period of ACSF perfusion without Fz followed with ACSF/Fz at the designated concentration for the remainder of the imaging session. Fz concentration in the imaging chamber typically reached equilibrium within 60 s after initiating ACSF/Fz perfusion, as judged by baseline BL signal in the neuron cell bodies.

### Image processing and data analysis

All image processing was performed using ImageJ (versions 1.54a through 1.54p). It is important to note that processing of bioluminescence imaging data requires compensation for the average dark pixel offset arising from the detector’s dark current characteristics and by small light leaks in the optical path. For each acquisition setting, dark time series were acquired under identical conditions but without illumination or addition of luciferin substrates. Dark time series with at least 60 frames were used to generate single-frame ‘dark’ images for each acquisition setting, containing the median value for each pixel over the time series. For single bioluminescence images of fusion proteins, the appropriate dark image was subtracted from the raw image. For time series, the appropriate dark image was subtracted from each frame of the time series. These dark-subtracted time series were summed using the ‘Z project’ function in ImageJ to create low-noise images for generation of regions of interest (ROIs) for analysis. Additional settings used for image display (e.g., thresholding, output scaling, lookup tables, etc.) are described for individual images in their corresponding figure legends.

To extract data from time series for downstream analysis, ROIs corresponding to individual whole cells were identified from summed time series images using the Cellpose v3.0.10^65^ ‘cyto’ model followed by manual curation. An additional ‘background’ ROI was drawn manually for each time series in an area devoid of cells. Mean intensities were then measured for each ROI at each time point from the dark-subtracted time series and data transferred to Excel for subsequent steps. The ‘background’ ROI value was subtracted from each cell ROI at each time point to account for any variable background signal arising from changes in light leakage (for example, when the injection port is uncovered) as well as any bioluminescence generated by extracellular luciferase/GECI molecules that escape from damaged or dead/lysed cells. Note that in cases where cells can be rinsed, the extracellular bioluminescence signal is typically negligible, but baseline shifts from changes in light leakage are inevitable, in our experience, making this step critical for accurate quantitation. For each ROI (whole cell) in a time series, we additionally calculate the mean dark value during the first 30 s prior to Fz injection and subtract this value from all time points to bring the dark baseline as close to zero as possible, compensating for any remaining offset in the data not captured by previous processing steps.

For Ca^2+^ indicators, the baseline luminescence emission intensity was determined for each cell by taking the average value of time points in the steady-state phase after initial Fz injection but before ionomycin or L-histamine injection, typically observed between 30 and 40 s after Fz injection (following a small transient cytosolic [Ca^2+^] increase) in most cells. The indicator signal at each time point was then calculated as the change in luminescence relative to the baseline luminescence (ΔL/L_0_) for each cell.

### Dose-response determination in cultured cells

*Luminescence dose response assay and AUC quantification.* N2a (Neuro 2a) cells (P30-P45) were plated 3 x 10^5^ cells/well in a six well plate and transfected with 2 μg of pcDNA3-CMV-CaBLAM using Lipofectamine 2000 (Invitrogen cat# 11668027). 48 hours later, cells were harvested, resuspended in external solution (ES) (135 mM NaCl, 10 mM HEPES, 2 mM CaCl_2_, 2 mM KCl), and distributed in equal volumes to a 96-well plate. Cephalofurimazine (CFz9, Promega cat# CS3553A01) was kept on dry ice until immediately before use and resuspended in 1 mL of 0.2 M Tris according to manufacturer’s instructions. Fluorofurimazine (FFz) (Promega cat# N4110) was resuspended in 525 μL of sterile phosphate buffered saline (PBS), aliquoted, and placed in -80 °C until use. Furimazine (Fz) (Promega cat# N1150) was kept on ice until use. A second 96 well plate was prepared with dilutions of CFz, FFz, and Fz, and substrates were transferred to the plate containing cells immediately before luminescence measurements. Luminescence was quantified with a Synergy HTX Multi-Mode Microplate Reader (BioTek) and Gen5 3.11 acquisition software using a top-positioned optic, 135 gain, 0.01 s integration time, and 1 mm read height at 33 s intervals for 20 min.

*Dose–response curve fitting and plotting.* Microplate reader data were imported from raw CSV files, and the area under the curve (AUC) was computed for each well using the trapezoidal rule using custom Python code (Python v3.11.7). Each CSV file included a row specifying substrate concentrations and a column for acquisition time in minutes. To visualize the dose-response curves, AUC values were grouped by concentration, and the mean ± standard error of the mean (SEM) was calculated. For each substrate, either a three-parameter Hill function or a hybrid model combining a Hill function with a linear decay term was fitted to the mean response values. Curve fitting was performed using non-linear least squares optimization (scipy.optimize.curve_fit) (scipy v.1.11.4 & matplotlib v3.8.0), with bounds applied to constrain biologically implausible parameter estimates.

The Hill-only model took the form:

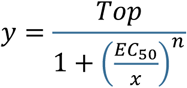

Where *n* is the Hill slope.

While the hybrid model added a linear decay term:

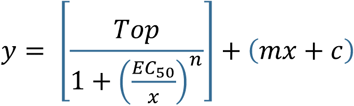

Where *m* is the slope of the decay and *c* is the offset. Goodness-of-fit was quantified using the coefficient of determination (R²).

*Bioluminescence imaging.* N2a (Neuro 2a) cells (P30-P45) were plated on 18 mm coverslips coated with 0.1 mg/mL poly-D lysine (Gibco A389040) at 1 x 10^5^ cells/dish in 35 mm dishes in DMEM + 1% FBS. Cells were transfected with 0.25 μg CaBLAM using Lipofectamine 2000 (Invitrogen #11668027). 24 hours after transfection, media was changed to DMEM + 1% FBS + 20 μM retinoic acid (Sigma #R2625). 24 hours later, cells were imaged with an iXon Ultra 888 EMCCD camera mounted on an Eclipse FN1 microscope (Nikon) with a 16X immersion objective (NA 0.8, WD 3.0, Nikon #MRP07220). Coverslips were rinsed with ES and placed on a Quick Change Imaging Chamber (Warner Instruments, #RC-41LP) with 500 μL ES. A reference fluorescence image was taken prior to bioluminescence acquisition at 30 MHz at 16-bit horizontal readout rate, 1x pre-amplifier gain, EM gain of 3, and exposure time of 0.1 s. Bioluminescence timeseries were acquired with Andor Solis 64 bit v4.32 at 10 Hz with the camera set to 0.09 s exposure time, 4.33 μs vertical pixel shift, normal vertical clock voltage amplitude, 10 MHz at 16-bit horizonal readout rate, 1x pre-amplifier gain, and 2×2 binning. Images were collected for 1 min in the dark prior to adding luciferin. Luciferins were diluted in ES and added to the coverslips to reach the final concentrations described in the figures. After 10 minutes, ionomycin was added to a final concentration of 2 μM and images were collected for another 30 minutes.

*Data processing and analysis.* Image stacks were processed using ImageJ as described above and data was exported from ImageJ as .csv files. The rest of the analysis was completed using custom Python code (Python v3.11.7, scipy v.1.11.4 & matplotlib v3.8.0) Traces from individual cells were first smoothed using a Bessel filter, (4th order, cutoff = 0.1 Hz) The baseline luminescence (L_0_) was calculated for each cell by taking the mean value of time points in the steady-state after luciferin injection, which was typically 9.3 – 9.4 minutes after luciferin injection. The maximum luminescence (L_max_) was calculated as the peak luminescence after ionomycin injection. Both L_0_ and L_max_ values were normalized to the mNeonGreen fluorescence values collected from the reference fluorescence image for each ROI.

### *In vivo* Ca^2+^ imaging in mice

#### Animals

Seven NDNF-Cre mice (2 female/5 male; 23-35 weeks old on imaging day; JAX stock #030757) were used for *in vivo* imaging to selectively express CaBLAM or GCaMP6s in neuron-derived neurotrophic factor (NDNF) expressing cortical layer 1 interneurons^66^. Three additional NDNF-Cre mice (0 female/3 male, each 14 weeks old on imaging day) were injected with a pan-neuronal CaBLAM to test peripheral luciferin delivery.

Mice were housed in a vivarium on a reversed light-dark cycle and had free access to food and water. All procedures were conducted in accordance with the guidelines of the National Institute of Health and with approval of the Animal Care and Use Committee of Brown University.

#### Surgical procedures

For the infusion experiments, four mice were injected with an AAV vector encoding a floxed version of the CaBLAM sensor (AAV9-ef1a-DIO-CaBLAM) and another three mice were injected with a floxed version GCaMP6s (AAV2/1-CAG-FLEX-GCAMP6s). For the peripheral luciferin delivery experiments, three mice were injected with an AAV encoding CaBLAM pan-neuronally (AAV2/9-hsyn-CaBLAM). Each animal was anesthetized (1-2% isoflurane), fitted with a steel headpost, and injected with viral constructs in a 3 mm craniotomy centered over left SI barrel cortex (-1.25 A/P, 3.5 M/L relative to bregma).

Viral injections were performed through a glass pipette in a motorized injector (Stoelting Quintessential Stereotaxic Injector, QSI). A glass window was then placed over the open craniotomy and cemented with dental cement (C & B Metabond). Mice received a single injection of a given construct at a location within 1mm of the center of our SI coordinates at depths of 500 μm. Each injection was 500 nL in volume and delivered at a rate of 100 nL/min. The glass injection pipette was then allowed to rest for an additional 10 minutes.

Mice receiving direct cortical infusions of luciferin were fitted with a cannula (Plastics One, C315DCS, C315GS-4). The tip of the canula was inserted below the dura at the very edge of the craniotomy. The canula and cranial window were then cemented in place together^67–69^.

#### Image acquisition

Images were acquired using an Andor Ixon Ultra 888 EMCCD camera and a 16x 0.8NA objective (Nikon CFI75) using Andor Solis data acquisition software (Andor Solis 64 bit, v4.32). Imaging data of the CaBLAM mice were acquired under no illumination in a light shielded enclosure (512 x 512 pixels after 2×2 binning) at a frame rate of 10 Hz (0.0955 s exposure) and an electron multiplication gain of 300. The GCaMP6s mice were imaged under epi-illumination from a X-Cite 120Q mercury vapor lamp (Excelitas Technologies) using an EGFP filter set (Chroma 49002, excitation filter: ET470/40x, dichroic: T495lpxr, emission filter: ET525/50m). The imaging parameters were otherwise the same as in the CaBLAM mice with the exception that the electron multiplication gain was set to 0.

The pan-neuronal CaBLAM mice were imaged using the same settings as above, with the exceptions of the use of a 4x 0.13NA objective (Olympus 1-U2B5222) at rates of 2 and 10 Hz (0.495 and 0.0955 s exposures). One of these mice was additionally imaged for an extended period (∼5 hours) under anesthesia after direct application of FFz. In this experiment images were acquired at 10 Hz as above, with all other parameters unchanged with the exception of the use of a 10x 0.30NA objective (Olympus 1-U2B5242). At the end of the 5-hour imaging session, in order to determine if bioluminescent subcellular neural processes can be identified using CaBLAM, we imaged at 1 Hz (no binning, 300 EM gain) using a 40x 0.8NA objective (Olympus 1-U2M587). Image stacks were acquired from multiple ROIs for ∼1 minute each.

#### Experimental procedures

Experiments were conducted 3 to 8 weeks post-surgery. For all animals, throughout data collection, a tactile stimulus was delivered to the right mystacial vibrissa pad using a piezo bender (Noliac). Tactile stimuli consisted of asymmetric sinusoidal deflections (10 ms rise and 15 ms fall time, 5 repetitions, 125 ms total stimulus length) delivered at random intervals between 20 and 25 s. Each animal received a minimum of 20 stimulus presentations. Animals were head fixed and allowed to freely run on a wheel.

During the direct cortical infusions of luciferin, the dummy cannula was carefully removed and replaced with an infusion cannula (Plastics One, C315IS-4) attached to a length of tubing connected to a 5 μL syringe (Hamilton 87930). FFz was reconstituted in sterile water (8.76 mM; 4.6 μmol/525 μL) and infused at a rate of 50-200 nL/min using a motorized injector (WPI UMP3) for a total volume of 500 nL. The infusion cannulas were kept in place for the duration of the imaging session.

For peripheral luciferin delivery, CFz9 (Promega) was reconstituted in 0.2M Tris buffer (8.40 mM; 8.40 μmol/1 mL) and 200 μL of the solution was injected retro-orbitally immediately prior (<1 min) to the start of imaging. In order to assess the viability of long duration imaging using CaBLAM, we removed the cortical window from one of the pan-neuronal CaBLAM mice under anesthesia (1-2% isoflurane). The mouse remained under anesthesia for the duration of the experiment that lasted ∼5 hours. 50 μL of FFz, reconstituted as above, was pipetted into a saline well over the craniotomy. Tactile stimuli we administered as above throughout the recording. The health and anesthetic depth of the mouse was monitored by breathing rate and toe pinch, checked at intervals of ∼20 minutes, during which the recording was paused.

#### Data analysis

Offline analyses of the *in vivo* imaging data from both CaBLAM and GCaMP6s were performed identically in Python 3.9.22 and Matlab R2024b (Mathworks) using custom scripts as well as the Matlab toolbox SUPPORT (https://github.com/NICALab/SUPPORT, accessed June 4, 2025, ^70^) and the Python package Suite2p (https://github.com/MouseLand/suite2p, v0.14.0, ^71^). The raw images were first spatially and temporally denoised using the SUPPORT toolbox using a pre-trained model available on the authors’ GitHub (“bs3.pth”). Using Suite2p, the denoised data was then motion corrected, automatically segmented into ROIs (followed by manual curation), and neuropil masks were created. The raw ROI traces were then imported into Matlab. The neuropil masks were used to apply a neuropil correction,

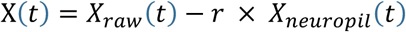

Where *X_raw_* is the fluorescent or bioluminescent signal of the ROI, *X_neuropil_* is the signal from the surrounding neuropil mask, and *r* is the decontamination factor, which was set to 1^72^. The resultant time series were smoothed using a 5-point moving average window.

To analyze stimulus evoked activity, we extracted -3 to 7 s windows centered on stimulus onset for each stimulus presentation and all ROIs. The pre-stimulus baseline period -3 to 0 s prior to stimulus onset was then used to calculate ΔF/F_0_ or ΔL/L_0_ for the fluorescent and bioluminescent data, respectively. We then identified responsive cells as those whose bootstrapped 95% CIs across all stimulus presentations in at least one of two response windows (0-1 S and 1-2 s) fell outside those of the cross-trial bootstrapped 95% CI of the baseline period. This yielded both positive and negatively responsive cells relative to baseline^36^.

### *In vivo* Ca^2+^ imaging in zebrafish

#### Fish husbandry and sample preparation

All procedures involving larval zebrafish (*Danio rerio)* were approved by the Institutional Animal Care and Use Committee (IACUC) at New York University Grossman School of Medicine. Adult zebrafish were maintained at 28.5 °C under a standard 14/10-hour light/dark cycle. Embryos were raised at densities ranging from 20 to 50 in 10 cm diameter petri dishes, each containing 25 to 40 mL of E3 medium with 0.5 ppm methylene blue added. At 1 day post-fertilization (dpf), larvae were kept in E3 medium without methylene blue. Larvae were screened for expression using a fluorescent stereomicroscope (Leica M165FC). Larvae were soaked in an E3 solution containing either 1:100 or 1:1000 vivazine (Promega Nano-Glo N2580) for 10 minutes and then mounted in 2% low melting point agarose. The tail (posterior to the pectoral fins) was then freed, and larvae remained in vivazine for the duration of the experiment.

#### Zebrafish lines

All larvae used were on the *mitfa -/-* background to remove pigment cells. Existing driver lines were *Tg(-6.7Tru.Hcrtr2:GAL4-VP16)* ^40^, called *Tg(hcrtr2)*, stl601Tg^39^, called *Is(nefma)*, and psi1Tg^41^, called *Tg(elavl3)*. In addition, two new transgenic lines were generated for this study.

First, the expression construct pTol2_slc1a3b:KalTA4 was generated using Gateway cloning to recombine p5E_slc1a3b^38^, pME_KalTA4^73^ and p3E_pA with pDestTol2CG2 from the Tol2Kit^74^. The resulting construct was microinjected into fertilized zebrafish eggs (using 1 nL of an injection solution containing 20 ng/µL DNA, 50 ng/µL Tol2 transposase mRNA and 10% phenol red). Injected F0 animals were outcrossed to wildtypes and F1 offspring was screened for germline transmission to obtain *Tg(slc1a3b:KalTA4)*, called *Tg(GLAST)*.

Second, the mNeonGreen-CaBLAM sequence was optimized for zebrafish translation using CodonZ^75^. The codon-optimized sequence was then synthesized (VectorBuilder) into a zebrafish Tol2 expression vector following 5xUAS before a SV40 PolyA tail. The expression plasmid was injected into *Is(nefma)* embryos at 1-cell stage, screened for green fluorescence, and raised to adulthood. Injected fish were outcrossed to wild-type fish to screen for founders. F2 and F3 embryos were used for experiments. All fish used for experiments were monoallelic for both the driver and CaBLAM.

#### Imaging and analysis

Anatomical imaging was done on a confocal microscope (Zeiss LSM 800) with a 20x/1.0NA water-dipping objective. Acquisitions were tiled to cover the entire length of the fish.

Simultaneous measurements of bioluminescence and behavior were made with a custom microscope that consisted of a high-sensitivity machine vision camera (Ximea MC023MG-SY with a SONY IMX174 sensor), a zoom lens (Navitar Zoom 7000, 1x at f/5.6), a custom chamber made from a magnetic mount (ThorLabs CP44F) and a glass-bottom dish (WPI FluoroDish FD3510), illuminated by a strip of 850 nm LEDs. Flux was measured with a photon-counting PMT (Hamamatsu H11890-110, 8 mm window) behind a 25 mm bandpass filter (ThorLabs BG-39) to block infrared light. The entire microscope was enclosed in a light-tight double-walled enclosure; when the IR light strip was off the PMT was at its noise floor (100 counts/s after 15 min). The microscope was controlled using custom software (LabView 2021) to acquire images (600x600 pixels, 250 fps, 1 or 2 ms exposure time) and sample from the PMT (40 Hz). Movement amplitude was defined as the number of pixels that changed intensity over a noise threshold (15/255) from frame to frame.

All analyses took place using custom code written in MATLAB (2024b, Mathworks Natick MA). To identify high-amplitude movements, we first defined a threshold that would reliably identify the largest events. To account for differences in imaging settings and variation in IR-reflectivity of each fish, we set the threshold at either the

99.9th percentile (1 ms exposure time) or 99.9975th percentile (2 ms exposure time) of movements for each fish. The first threshold-crossing event in a given second was defined to be the beginning of each high-amplitude, and events were spot-checked to ensure that they were comparable (i.e. prolonged and uncoordinated large-amplitude tail movements) by post-hoc examination of saved videos. We processed the vector of counts from the PMT by interpolating with a spline fitting algorithm to match the timebase of behavior and then smoothing the vector with a 0.25 s square window. For each fish, we extracted a response defined as the counts 2 s before and 30 s after a threshold-crossing event. The baseline was defined as the mean of the first 2 s of the response. To facilitate comparison across genotypes, we subtract the baseline from the response and divide the result by the baseline i.e. (counts(t) - baseline) / baseline.

Widefield imaging of bioluminescence was performed in two ways. First, slow images (30 s exposures) were taken with the machine vision camera above with the aperture set at f/2.8. Images were binned 4×4, and a single image was generated by taking the standard deviation across pixels in the resulting stack (83 frames), and then up-sampling 4×4 to overlay with an infrared-illuminated reference image. Second, high-speed imaging was performed using an intensified camera (HiCAM Fluo, Lambert Instruments) through with a Cousa 10x/0.5 NA air objective^76^. A sequence of images was captured over 2 minutes at 20 frames per second. Raw images were processed by subtracting a dark-count background image, smoothed with 1) a 3-frame rolling window 2) a 2×2 Gaussian blur and 3) a 3-pixel median filter in (x,y,t). A single-frame average projection was then generated to represent basal intensity. The final result was generated by subtracting the basal intensity from each image slice.

## Supporting information

Supplementary Results and Discussion, Figures

Supplementary Data File 1 (peptide sequences)

Supplementary Data File 2 (primer list)

## Acknowledgments

We wish to thank several individuals who supported this project as allies: Dr. Stephen Adams for many years of insightful discussions about sensors, Ca^2+^ buffers, and other subjects; Dr. Pauline Wales for her invaluable help with camera and objective selection and optimization; Dr. Kelly Monk for providing zebrafish lines; Dr. Matthew Lovett-Barron for helpful advice on zebrafish experiments; and Dr. Leona Flores and Dr. Michael Oberholzer for invaluable scientific discussions and moral support through very challenging times. We would also like to thank all members of our Bioluminescence Hub (http://www.bioluminescencehub.org/) laboratories for their feedback, discussions, and thoughtful comments throughout the progression of this work.

## Competing interests

The authors declare no competing or financial interests.

## Author contributions

Conceptualization, GGL, ELC, DL, CIM, UH, and NCS; Validation, GGL, ELC, JM, SV, DC, DL, CIM, UH, NCS; Formal Analysis, GGL, ELC, JM, DS, NCS; Investigation, DB, JH, SL, DKN, BNS, MOT, SV, DC, RO, DH, ABB, ATZ, AH, JC, RM, HG, EG, YZ, MK, DS, NCS; Data Curation, GGL, ELC, JM, KIN, DS, SS, YZ, NCS; Writing—Original Draft, GGL, ELC, NCS; Writing—Review, Revision, & Editing, GGL, ELC, JM, MC, BNS, DL, CIM, UH, JJA, AA, YZ, SS, DS, NCS; Visualization, GGL, ELC, JM, DS, NCS; Supervision, DL, AA, CIM, UH, TC, SS, DS, KIN, NCS; Project Administration, UH, JJA, TC, KIN, DS, NCS; Funding Acquisition, DL, CIM, UH, AA, BNS, JC, KIN, DS, NCS.

## Data and reagent availability

Raw and processed data sets from experiments performed in this study are freely available via the Brown Digital Repository (https://doi.org/10.26300/7sg5-w257) and as input data for reproducing analysis with the custom code developed in this study (https://github.com/Shaner-Lab/CaBLAM). Bacterial and mammalian expression plasmids encoding SSLuc, GeNL_SS, and CaBLAM will be deposited for distribution by AddGene (pending); prior to availability from AddGene, plasmids will be shared with non-profit researchers upon request to NCS.

## Code availability

Custom code used in data collection, processing, and analysis is freely accessible on GitHub (https://github.com/Shaner-Lab/CaBLAM).

## Funding

This work was supported by the National Institutes of Health (R21EY030716, R21MH101525, R01GM121944, R01CA279813, R01NS120832, R21EY036659, R34DA059500, R21NS115437, R01EY035691, R01MH124811, F32NS134617, and U01NS099709), the National Science Foundation (CBET-1464686, DBI-1707352, and DBI-2208914), a Research Seed Award from Brown University’s Office of the Vice President for Research (OVPR), and the Allen Discovery Center for Neurobiology in Changing Environments, funded by the Paul G. Allen Frontiers Group. The funders had no role in study design, data collection and analysis, decision to publish, or preparation of the manuscript.

---

**Extended Data Figure 1.**
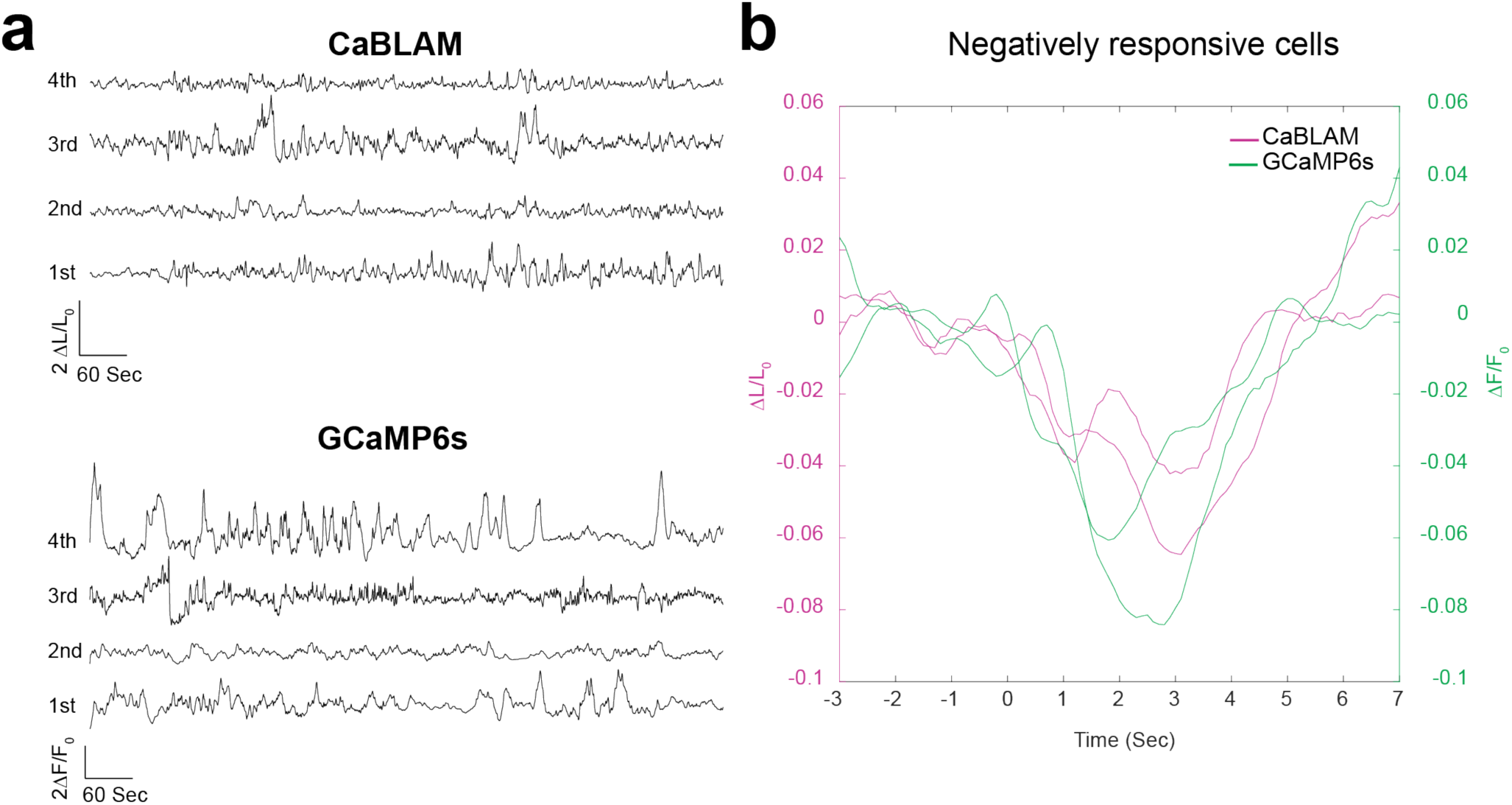
Classifying cell responses to vibrissa stimulation. **a,** Example 8-minute CaBLAM and GCaMP6s traces. In both, positively responsive cells were sorted by SNR. From bottom to top, traces from the 1^st^, 2^nd^, 3^rd^ and 4^th^ SNR quartiles **b,** Example of average response from 2 negatively responsive cells in the CaBLAM and GCaMP6s groups. Note the left and right y-axes.

**Extended Data Figure 2.**
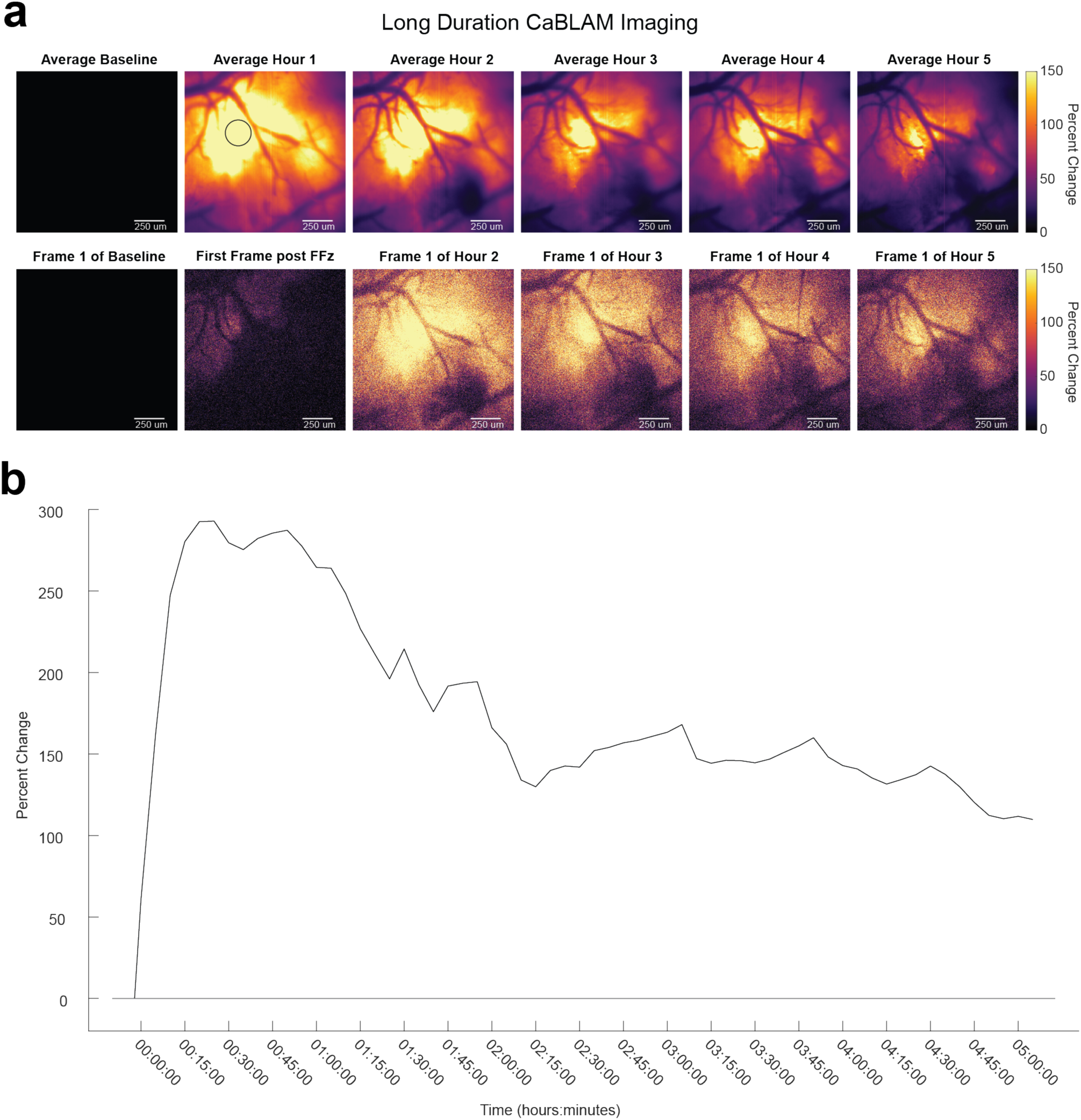
Bioluminescence time course with a single direct cortical application of FFz in a pan-neuronal CaBLAM mouse. **a,** *top row*, average images from each hour of recording after direct cortical application of FFz in one pan-neuronal CaBLAM mouse*; bottom row*, first single frames from each hour after FFz administration. **b,** Bioluminescence from a circular ROI (black circle in **a**, top row, second from left) binned at 5-minute intervals expressed as percent change from a 1-minute (dark) baseline prior to FFz administration.

**Extended Data Figure 3.**
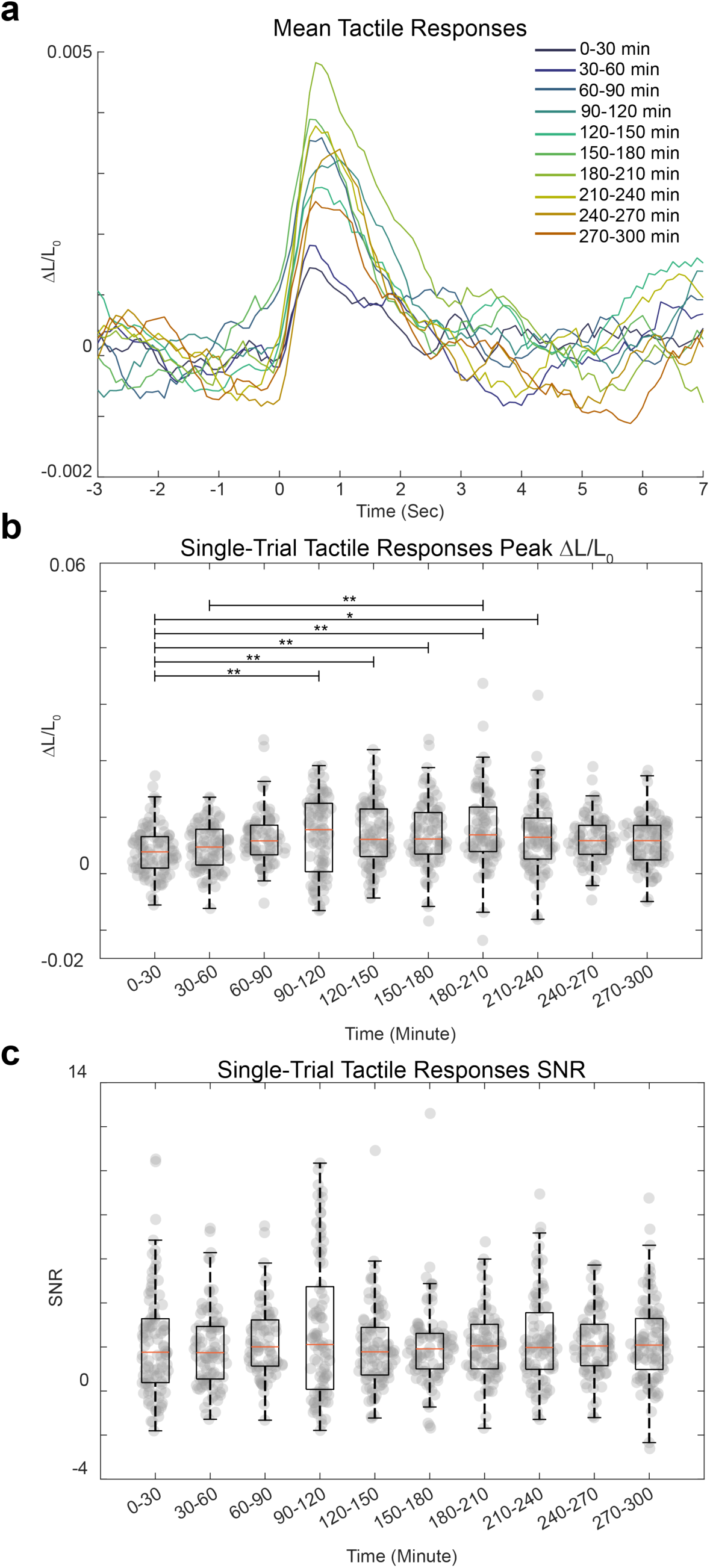
Tactile responses over time in an anaesthetized pan-neuronal CaBLAM mouse with a single direct cortical application of FFz. **a,** Average CaBLAM tactile response from ROI marked in Extended Data Fig. 2A, binned at 30-minute intervals after the administration of FFz. **b,** Box plot of the single-trial peak ΔL/L_0_ binned at 30-minute intervals. Orange lines indicate the median, boxes enclose the middle 2 quartiles of the data, whiskers extend 1.5 IQRs above and below. Gray dots are single data points. Horizontal lines at top indicate significant differences between intervals, where * indicates *P* < 0.05 and ** indicates *P* < 0.01. **c,** box plot of SNR same conventions as in B).

**Extended Data Figure 4.**
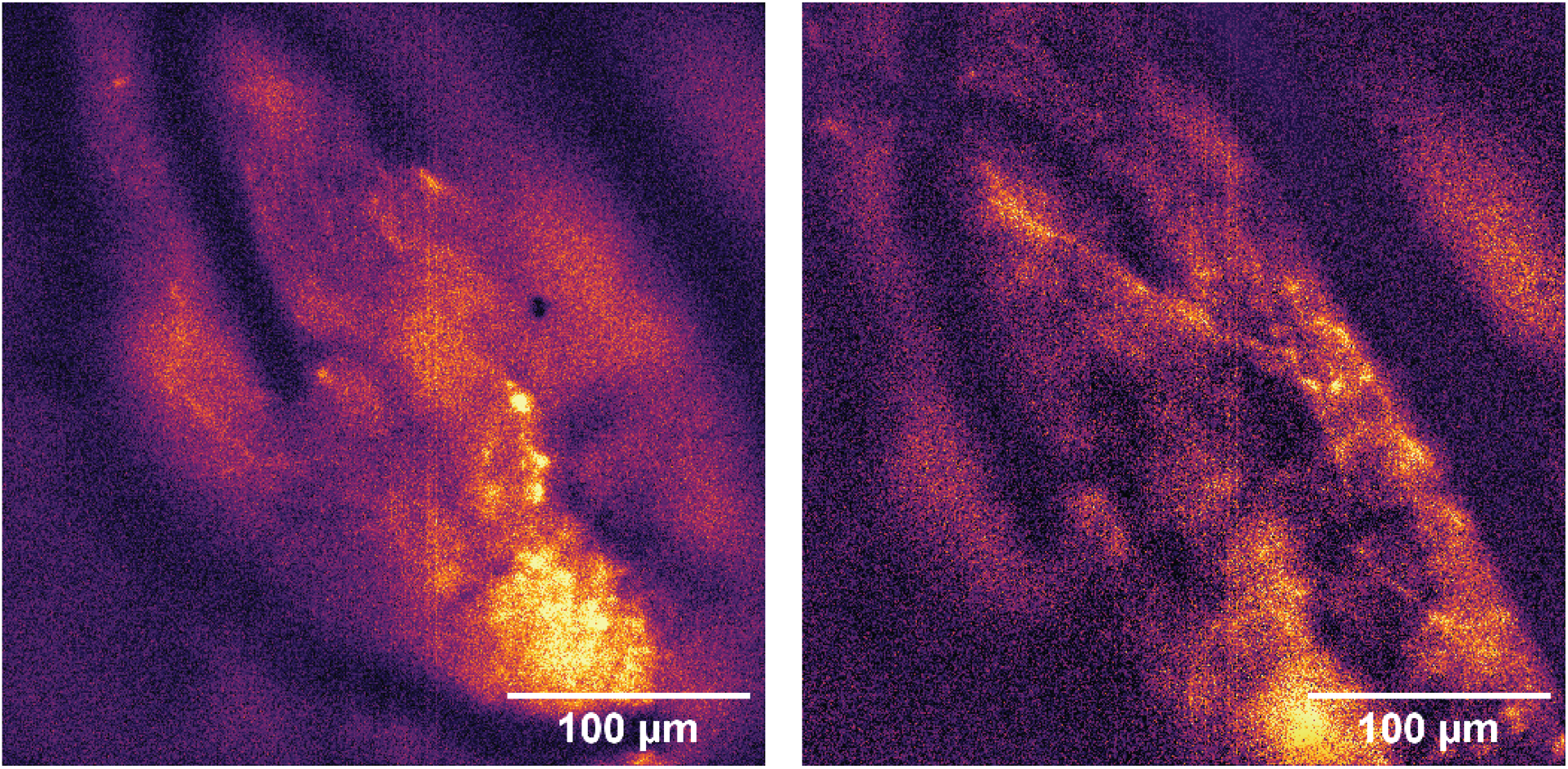
Representative average images of neural processes acquired at 40x *after* the 5-hour recording from a pan-neuronal CaBLAM mouse. Images were acquired at 1 Hz with no binning at 300 EM gain.

## Notes

https://github.com/Shaner-Lab/CaBLAM

https://doi.org/10.26300/7sg5-w257

### Competing Interest Statement

The authors have declared no competing interest.

### Summary of Updates

Key changes include comprehensive characterization of multiple substrates (fluorofurimazine, cephalofurimazine, and furimazine) with dose-response curves and brightness comparisons; three CaBLAM variants with different calcium affinities (KD ~439 nM, ~3 μM, and ~280 nM); expanded in vivo imaging capabilities including awake, head-fixed mice with multi-hour recordings (>5 hours), peripheral substrate delivery methods, and subcellular resolution imaging; and new zebrafish applications using transgenic lines with behavioral correlation and high-speed imaging. The methods section has been enhanced with detailed substrate preparation protocols, improved in vitro characterization using validated free Ca^2+^ calculations, and advanced data analysis including SUPPORT toolbox denoising and Suite2p motion correction. Complete data and code are now available via Brown Digital Repository and GitHub.

