## Supplementary Results and Discussion, Figures for "CaBLAM! A high-contrast bioluminescent Ca^2+^ indicator derived from an engineered *Oplophorus gracilirostris* luciferase"

### Rational design and directed evolution of high-activity soluble OLuc variants

While NanoLuc displays very high light output utilizing the proprietary furimazine (Fz) substrate, we initially intended to generate a variant derived from *Oplophorus* luciferase that would perform equally well with coelenterazine (CTZ), which is substantially less costly and more readily available. To this end, we started by evaluating the OLuc variant "eKAZ," which was originally described as having higher activity than NanoLuc when utilizing CTZ[1]. We found that the *in* vitro activity of eKAZ appeared highly sensitive to buffer conditions (particularly detergents) and that the activity of purified eKAZ also degraded rapidly upon storage, making it nearly impossible to quantitatively characterize its substrate preference or brightness relative to other luciferases.

To help enhance the luminescence signal of eKAZ by increasing the quantum yield of bioluminescent emission through Förster resonance energy transfer (FRET) to a bright acceptor chromophore[2], we tested several fusions to mNeonGreen[3], using an approach similar to that described by Saito et al. in their design of the original NanoLantern probe[4] (an optimized fusion of the YFP variant mVenus[5] to RLuc8[6]) and later for the GeNL probe[7] (an optimized fusion of mNeonGreen to the N terminus of NanoLuc[8]). We found that a fusion of mNeonGreen to the N terminus of eKAZ, with a 7 amino acid C-terminal deletion to mNeonGreen and proline-proline linker (**Supp. Fig. 1A**), produced a high FRET efficiency (**Supp. Fig. 1B**) and slightly enhanced luminescence intensity compared with eKAZ alone, though at this stage it was difficult to quantify the brightness improvement. We used this construct as the basis for further directed evolution to improve the luminescence activity of this variant, with the expectation that mutations conferring increased CTZ oxidation rates at the expense of luciferase photochemical quantum yield would still display bright luminescence output via FRET coupling.

Because of these clear limitations of eKAZ, we sought to introduce mutations that would improve both its brightness and stability. Following our traditional directed evolution approach, we performed multiple rounds of random and site-directed mutagenesis of the eKAZ variant (fused to mNeonGreen as described above). We screened libraries simply by spraying a sterile solution of CTZ onto agar plates in a dark room and identifying the brightest colonies either by eye or using a homebuilt whole-plate imaging setup. The original clone was already active enough to observe by eye in *E. coli* colonies constitutively expressing the protein from the pNCS expression vector[3]. We immediately identified several key locations in the protein that led to marked increases in colony brightness relative to the original clone. Three mutations (L30Q, S37F, and L92M) were located at an opening to the putative substrate binding pocket[9], one (T13I) was on the first beta strand proximal to the C-terminal peptide, and one (I56T) was internal and possibly part of the binding pocket (numbering based on published NanoLuc crystal structures[9,10]). We targeted positions 56, 92, and 93 in a directed library and identified the mutant L92M/P93E as the most improved clone, designated "eKAZ-L6."

Random mutagenesis of eKAZ-L6 (fused to mNeonGreen as described above) revealed relatively few improved clones, with the mutation A33D present in the brightest variants. We constructed a directed library targeting position 33 along with previously identified position 30 and positions 68 and 72 on the neighboring alpha helix. We identified the top clone from this library containing the mutations L30V/A33T/F68D/L72H, designated "eKAZ-L9," that displayed both higher luminescence emission and improved soluble protein yield. We suspect that this improvement in solubility arises from substitution of hydrophilic side chains in place of two external hydrophobic side chains (Phe68 and Leu72). At this stage, the *in vitro* activity of eKAZ-L9 had improved substantially compared to the original eKAZ clone[1] and the enzyme had gained a considerably higher tolerance for varied buffer conditions but still did not display activity levels comparable to NanoLuc using either CTZ or Fz as the substrate.

We reasoned that at this stage of development, merging our improved eKAZ mutations with some mutations from related luciferases such as NanoLuc could be beneficial. Several groups have described the development of split NanoLuc complementation assays and sensors[7,11–14]. The most widely adopted bimolecular version, dubbed "NanoBit," was optimized for low affinity between the split pieces to avoid self-complementation[11]. The small fragment ("SmBit") of the NanoBit pair comprises the last beta strand and is only 11 amino acids long. NanoBit was developed by optimizing the large fragment ("LgBit") by directed evolution with selection for the highest BL signal when the C-terminal peptide was added, along with the identification of several functional C-terminal peptides with lower and higher affinities to LgBit (**Supp Table 1**). *We reasoned that the activity- and folding-enhancing mutations present in NanoBit could be useful in improving eKAZ-L9*, and that substitution of the last 11 amino acids with the highest-affinity peptide identified (**VSGWRLFKKIS**, designated as "86" in the NanoBit publication[11]) could further enhance the stability of the folded protein.

In an effort to preserve the substrate specificity of eKAZ-L9 for CTZ, we designed a variant that included only the external NanoBit mutations (several of which are carried over from NanoLuc), reasoning that specificity for Fz was likely to be determined by internal mutations influencing the substrate binding pocket. The resulting designed mutant, designated "eKAZ-11S-L9," contains 22 mutations relative to eKAZ, including the 6 mutations we previously identified through directed evolution, 12 external mutations from NanoBit, and the remaining 4 non-aligned residues arising from substitution of the high-affinity C-terminal peptide "86" in place of the original C terminus. We then tested eKAZ-11S-L9 by constructing a new fusion to mNeonGreen identical in design to GeNL[7], which uses mNeonGreen with a 10 amino acid C-terminal truncation and a Gly-Phe linker before the luciferase with a 5 amino acid N-terminal truncation. We assayed this mutant *in vitro* in parallel with GeNL using *E. coli* extracts, normalizing to the fluorescence intensity (since mNeonGreen is fused to the luciferase, it can act as an internal standard). These experiments revealed that the mNGΔ10C-GF-Δ5NeKAZ-11S-L9 construct was already apparently ~5-fold more active than GeNL on a per-molecule basis using Fz, and ~4-fold using CTZ.

At this stage, since activity remained higher with Fz than CTZ even after omitting internal mutations expected to retain preference for CTZ, we reasoned that additional mutations borrowed from NanoLuc might help improve the enzyme further and might additionally enhance activity with both Fz and CTZ. To test this hypothesis, we generated a new variant by introducing most of the internal mutations present in NanoLuc and/or NanoBit that we had previously excluded in our design: A4E/G15A/Q18L/L27V/K43R/I76V/I90V/P115E/Q124K, in addition to altering the C-terminal peptide to **VSGWRLFEKISN** to bring its sequence midway between that of NanoLuc and NanoBit. The resulting variant was designated "eKL9h" and retained its high activity with CTZ while also displaying increased activity with Fz and other CTZ analogs. When expressed in *E. coli,* the mNG FRET fusion to eKL9h could be fully extracted with B-PER lysis reagent, while GeNL (the equivalent mNG FRET construct with NanoLuc) was largely insoluble (**Supp. Fig. 2**), suggesting that eKL9h might also perform more reliably as a fusion partner to other proteins.

To further increase the brightness of eKL9h, we focused on the C-terminal peptide, individually constructing and characterizing 8 different peptides selected from those characterized during the original development of NanoBit (**Supp. Table 1**). Peptides were chosen to represent the full range of NanoBit affinities, with the expectation that there would be an optimum affinity for activity in an intact OLuc variant. We found that the C-terminal peptide we had originally chosen was already superior to all others tested, though many performed similarly, and we observed no clear correlation between peptide affinity and luciferase activity. While characterizing these C-terminal peptide variants, we serendipitously identified one aberrant clone that displayed approximately 2-fold higher activity than eKL9h. Sequencing revealed that this clone had a frame shift in the C-terminal peptide that led to the unintended sequence **GYRLLKKSCDV**. Based on this sequence, we generated variants with truncations of one to four amino acids from this novel C terminus as well as a set of variants with a GYRLLxK or GYRLxxK leading sequence followed by a variety of rationally designed variations on the trailing sequences (**Supp. Table 1**). We identified the peptide **GYRLLKKISN** as the most optimal of those tested, leading to the luciferase clone eK-ISN.

Based on subtle correlations between sequence and activity in C-terminal peptide variants, we speculated that the lysine side chains at positions 168 and 169 in this new C-terminal peptide might be important for high activity of this lineage of OLuc variants. We explored substitutions to other residues with side chains that could potentially interact with these lysine side chains. We constructed a model of the eKL9h protein structure by homology modeling with a NanoLuc X-ray structure template[10] in Rosetta[15] and identified positions 9, 10, 13, and 146 to target for randomization based on their proximity to the key lysine side chains. We constructed single point mutants for all possible amino acids at these four positions and then measured the luminescence intensity for each clone in comparison with the standards (GeNL and the GeNL-style mNG fusion to eK-ISN) with CTZ, *h*-CTZ, *bis*-CTZ, and Fz substrates. Several substitutions at positions 13 (original residue Thr, replaced by Glu, Ile, His, or Leu) and 146 (original residue Asp, replaced by Glu, Asn, His, or Ser) produced clones with activity equal to or better than that of eK-ISN. We next shuffled these favorable substitutions, producing all possible double mutants. Of these, eK-ISN with the additional mutations T13I and D146E was ~2-fold brighter *in vitro* than eK-ISN, leading to the clone eK-ISN-IE, which we used for subsequent detailed characterization *in vitro* and in mammalian cell lines. Since this clone later proved to be well suited to insertion of sensor domains, we gave this clone the "official" name of Sensor Scaffold Luciferase, or SSLuc.

SSLuc contains 37 amino acid substitutions relative to the active domain from wild-type OLuc, 34 substitutions relative to eKAZ, and 26 substitutions relative to NanoLuc, comprising between 15% and 22% of the total amino acids in this small protein. Peptide sequences of SSLuc, related published OLuc variants, and notable intermediate variants from SSLuc directed evolution are provided in **Supplementary Data File 1**.

Interestingly, Inouye et al[16] recently investigated substrate specificity in NanoLuc and demonstrated that positions 18 and 27 were important in determining substrate preference, with reversion of these positions to the wild-type residues increasing specific activity with *bis*-CTZ relative to Fz. We did not exhaustively characterize the substrate preference for each mutant generated during the directed evolution of SSLuc in this study, but these intermediate variants may be useful for future studies of structural determinants of substrate preference in this enzyme lineage.

***In vitro* characterization of SSLuc**

We initially compared the properties of NanoLuc and SSLuc using GeNL-style mNeonGreen (mNG) fusions (i.e., mNeonGreenΔ10C-GF-Δ5NSSLuc, hereafter abbreviated to GeNL_SS). The FRET efficiency of SSLuc to mNG in the GeNL_SS construct is slightly lower than that of NanoLuc to mNG in GeNL (**Supp. Fig. 1B**), suggesting that the orientation of the luciferase and FP domains in these two fusions is slightly different. The use of mNG fusions provided an internal fluorescent standard, enabling accurate dilution of the two proteins to equal concentrations for each measurement using either the optical absorbance or the fluorescence emission to measure protein concentration (relative to a purified mNG standard at known concentration). Fusion to mNG is not expected to influence the kinetics or substrate specificity of either luciferase. We found that the substrate specificity of SSLuc was similar to that of NanoLuc, with 20±3-fold higher light output when utilizing native CTZ, 3.8±0.3-fold higher with h-CTZ, 4±1-fold higher with *bis*-CTZ, and 10±1-fold higher output with Fz relative to NanoLuc when measured *in vitro* (**Supp. Fig. 3A**), based on steady-state signals from each enzyme. Both SSLuc and NanoLuc produced large initial intensities that decayed rapidly when using *h*-CTZ as a substrate but did not produce a steady-state signal *in vitro* (**Supp. Fig. 3B**), with SSLuc displaying a larger area under the curve than NanoLuc. These observations suggest that the oxyluciferin produced from *h*-CTZ may be a potent inhibitor of both luciferases and/or that oxidation of *h*-CTZ may lead to covalent inactivation of the active site, as seen by others in *Gaussia* luciferase[17]. In our hands, both CTZ and Fz produced steady-state luminescence at all but the highest GeNL_SS concentrations, with Fz producing only slightly higher peak intensity but longer steady-state ("glow") emission (total light output ~2-fold higher) compared with CTZ.

When expressed in *E. coli*, GeNL_SS produced a much higher fraction of soluble protein than GeNL (**Supp Fig. 2**), as measured by the relative mNG fluorescence in the pellet and lysate, suggesting that the folding efficiency and/or solubility of GeNL_SS was superior to GeNL, at least in this expression context. Because mNeonGreen folds robustly in fusion proteins in most cases, total fusion protein concentrations inferred from mNG fluorescence were expected to be accurate, while the active/functional enzyme concentration *could differ substantially if the enzyme itself folds poorly in the fusion*. We suspect this is likely the case with the GeNL fusion (NanoLuc), which could also explain its tendency to form insoluble aggregates in *E. coli.* We additionally found that recombinant NanoLuc and GeNL typically remain stable for ~12 hours following extraction (with or without further purification) but frequently lose luciferase activity rapidly thereafter, while SSLuc and GeNL_SS appear much more shelf stable. We initially suspected that this effect could be due to oxidation of the single cysteine present in the luciferase but found that addition of DTT or β-ME to purified protein stocks did not prevent loss of activity. We now speculate that the GeNL fusion does not maintain a stable fold of the NanoLuc portion of the protein, leading to a slow decay in activity relative to stable mNG fluorescence after extraction from *E. coli,* possibly due to self-aggregation of misfolded NanoLuc domains. By contrast, SSLuc and GeNL_SS retain stable luciferase activity under prolonged storage at 4C, even in clarified *E. coli* lysate without further purification.

As a result of these solubility and stability differences, accurate and repeatable comparative assays of luciferase activity could only be made when proteins were extracted and assayed on the same day; *if the proteins were stored for longer times, we consistently measured an unrealistically high relative brightness of SSLuc/GeNL_SS relative to NanoLuc/GeNL.* Taken together, these observations suggest that differences in solubility and folding between SSLuc and NanoLuc when expressed in *E. coli* are likely responsible for the observed *in vitro* performance improvements in SSLuc, and that the $V_{max}$ of correctly folded SSLuc may in fact still be lower than that of NanoLuc. Unfortunately, the context-dependent folding of NanoLuc greatly limited our ability to quantitatively compare the two enzymes, even in mammalian cells (see below).

***In cellulo* characterization of SSLuc**

We initially attempted to compare the brightness of NanoLuc and SSLuc in mammalian cells by constructing plasmids for extracellular membrane-bound expression under the rationale that this would most closely replicate *in vitro* assay conditions, especially with respect to substrate availability. Ultimately, we chose to compare cytosolic constructs rather than extracellular ones, as discussed in more detail below. For extracellular targeting, NanoLuc and SSLuc full-length coding sequences alone were directed to the plasma membrane by an N-terminal signal peptide derived from *Gaussia* luciferase followed by the B7 hinge alpha helix[18], fused on the C terminus to mNeonGreen. This construct was expected to place the luciferase on the *extracellular* side of the membrane and the fluorescent protein on the *cytosolic side* of the membrane. We chose this configuration after observing that ***neither*** GeNL or GeNL_SS could be cleanly localized to the membrane when placed entirely on the extracellular side of the B7 hinge sequence or a GPI-anchor sequence, suggesting that either the mNeonGreen component or the luciferases may interfere with trafficking.

Cells expressing the extracellular NanoLuc and SSLuc constructs also displayed major differences in localization that prevented us from accurately quantifying brightness differences. While SSLuc localized well to the plasma in this construct, NanoLuc remained trapped in the ER, Golgi, and nuclear envelope without *any* clear localization to the plasma membrane. We also observed significantly higher signal with SSLuc versus NanoLuc—far higher than expected from the *in vitro* characterization of the two enzymes. We suspect that the reduced solubility of NanoLuc relative to SSLuc and/or the presence of an unpaired external cysteine side chain on NanoLuc leads to misfolding and aggregation that both eliminate luciferase activity and prevent extracellular display of the enzyme. Because of the substantial differences in trafficking between the NanoLuc and SSLuc proteins in these constructs, we could not quantitatively compare the two enzymes in this assay.

For cytosolic expression, we used the GeNL and GeNL_SS coding sequences driven by a CMV or CAG promoter. We found that the fluorescence brightness of individual cells was generally not correlated linearly to the luminescence brightness for either cytosolic construct, as would be expected in a luciferase-FP fusion construct. Our interpretation of this observation is that delivery of the luciferin across the cell membrane is rate-limiting when these luciferases are expressed in cytosol with the substrates we tested (CTZ, *h*-CTZ, *bis*-CTZ, and Fz). We concluded that cytosolic GeNL and GeNL_SS expression was also not a suitable test system for quantitative brightness comparisons in mammalian cells.

Finally, to assay the *in cellulo* brightness of SSLuc versus NanoLuc without the potential complications of using a fusion protein, we finally constructed plasmids to express mNeonGreen and each full-length luciferase separated by a self-cleaving T2A peptide[19] to provide 1:1 stoichiometry of each protein individually. This allowed us to maintain the internal fluorescent standard while measuring the luciferase output independently, which we hypothesized would minimize fusion-dependent misfolding of NanoLuc. Imaging these constructs expressed in U2OS cells revealed that only cells expressing very low levels of recombinant protein displayed an apparently linear relationship between fluorescent and luminescent intensity. This observation supports our hypothesis that delivery of luciferin (Fz) across the cell membrane is rate-limiting. Restricting analysis to the linear portion of each curve, we find that SSLuc appears to produce approximately half the photons of NanoLuc when not fused to a FRET acceptor, suggesting that it is indeed "dimmer" on a per-molecule basis when both proteins are properly folded.

**Subcellular imaging of GeNL_SS-tagged proteins**

We fused GeNL_SS to several proteins to evaluate its utility in subcellular localization imaging in living cells. Fusions to H2B, LifeAct, and TOMM20 displayed expected localization in cells for both GeNL and GeNL_SS fusions when expressed at moderate levels, but both failed to localize well in the highest-expressing cells. This phenomenon is common to even the most well-behaved fluorescent proteins, and so we note that researchers should *always* be wary of tagged protein localization in the brightest subset of transfected cells. Interestingly, when we compared U2OS cells expressing identical GeNL-H2B and GeNL_SS-H2B fusions, we noticed that the toxicity of the GeNL fusion appeared quite high, with most expressing cells dying within 24h of expression onset, while cells expressing the GeNL_SS fusion remained viable and were able to divide successfully. When considered along with its poor localization in extracellular plasma membrane-directed fusions, these observations support the hypothesis (as noted above) that GeNL undergoes a substantial amount of misfolding in mammalian cells just as it does in *E. coli*, especially when targeted to oxidizing compartments, possibly in part because of the unpaired cysteine near its C terminus which is absent in SSLuc/GeNL_SS. It remains possible that NanoLuc alone (without the mNG FRET acceptor) could be a suitable fusion tag for at least some proteins in mammalian cells, but because we would be unable to easily compare expression levels without the fluorescent component, we chose not to evaluate such fusions in this study.

**Oligomeric state of GeNL and GeNL_SS**

We constructed fusions of GeNL and GeNL_SS to the C terminus of the CytERM peptide used for the OSER assay, a standard assay of "monomeric behavior" for fluorescent proteins expressed in mammalian cells. ***Neither protein*** displayed OSER/NE ratios consistent with currently accepted values for monomeric fluorescent proteins[20] , suggesting some amount of self-association. Qualitatively, ER morphology was more evenly labeled in CytERM-GeNL_SS expressing cells compared with CytERM-GeNL, and OSER structures appeared more often in cells with lower expression level (as measured by fluorescence intensity) of CytERM-GeNL compared to CytERM-GeNL_SS, suggesting that GeNL_SS may self-associate to a lesser degree than GeNL. We conclude that while self-association of both GeNL and GeNL_SS appears to be low—neither appears to be a *strong* dimer in this assay—neither probe performs as well as a monomeric fluorescent protein tag for imaging fusion protein localization.

CytERM fusions to NanoLuc and SSLuc alone (without the mNG acceptor) were too dim to obtain images with a signal to noise ratio suitable for quantitative analysis, and as a result we were unable to determine whether the mNG fusions are more or less soluble and monomeric than the untagged luciferases. Additional characterization of the oligomeric state of NanoLuc and SSLuc under physiological conditions will be necessary to clarify the differences between these two luciferases. Taking all data and observations into account, SSLuc appears to have higher solubility, more efficient folding, lower toxicity, and less tendency to self-associate than NanoLuc, but has several areas that would benefit from future engineering to improve catalytic rate, performance as a fusion tag, and overall "inertness" in host cells.

**Development and evaluation of a high-contrast BL GECI**

We next explored the design of new BL GECI architectures based on SSLuc. Since the related split luciferase NanoBit[11] was specifically developed as a *reversible* complementation system, we chose the loop between the last two beta strands of the luciferase domain of SSLuc as our split point for insertion of sensor domains, with the goal of generating indicators with a large change in BL output between low and high cytosolic $Ca^{2+}$ concentrations (i.e. [$Ca^{2+}$] between ~100nM and ~1µM). We trialed several topologies for the sensor, including circularly permuted variants and simple sensor domain insertions of RS20-Calmodulin (RS20-CaM), CaM-RS20, and Troponin C (**Supp Fig. 4**). All versions of the initial GECI displayed readily measurable and fairly large changes in BL between $Ca^{2+}$-free and high-$Ca^{2+}$ conditions *in vitro*, with the highest contrast consistently observed from variants with RS20 followed by CaM inserted before the C-terminal luciferase peptide (**Fig 1A, B**). Contemporaneously with our exploration of all these topologies, an analogous RS20-CaM configuration was also identified by Farhana et al[21], in which the insertion is an entire GCaMP sensor domain, including the circularly permuted GFP component. In our designs, the FP, if present, remains fixed at the N terminus of the luciferase as a FRET partner to enhance the quantum yield of emission[2,4,7,22], and the $Ca^{2+}$-sensitive signal is restricted to the BL channel, not both channels, unlike GLICO's bimodal signal. Despite this substantial difference and the use of the SSLuc scaffold rather than NanoBit, we found that the tripeptide linker sequences flanking the sensor domains of GLICO were suitable for our new GECI designs as well, and so we adopted these in our final design.

Unexpectedly, the initial clones of our novel BL GECI lineage displayed $Ca^{2+}$ affinities far too high to be practical in living cells, often with $K_D$ < 10nM [$Ca^{2+}$]. We explored several ways of altering the affinity, including the use of alternate RS20 and CaM sequences derived from existing fluorescent GECIs[23–25] and lower-affinity designed variants of the RS20/CaM pair[26], altering the linker length and composition between the RS20 and CaM, altering linker lengths and compositions between the luciferase and sensor domain, and rational directed mutations to the CaM domain's EF-hand motifs. This combination approach allowed us to select for the brightest clones while also determining which design factors influenced the $Ca^{2+}$ affinity. Ultimately, the best balance of brightness, affinity, and Hill coefficient was achieved using the CaM and RS20 sequences from GCaMP6s coupled with a single point mutation (N98I in full-length calmodulin) chosen from among several naturally occurring mutations in human calmodulins leading to heart rhythm defects[27], producing apparent $Ca^{2+}$ binding affinities around 500nM. This $K_D$ target was selected to align the most sensitive part of the GECI's response curve with a physiologically relevant range of $Ca^{2+}$ concentrations—for example, $Ca^{2+}$ concentrations in the cytosol of rat CA1 hippocampal pyramidal neurons in slices have been measured using small molecule dyes[28] and are estimated to range from 32-59nM when resting and up to >300nM in response to a single action potential. The process of evaluating the many variables contributing to the calcium-binding behavior of this BL GECI architecture produced multiple additional CaBLAM variants suitable for future optimization and engineering for specific biological use cases requiring higher and lower $Ca^{2+}$ affinities (**Fig. 1C, Table 1**).

To maximize the BL signal contrast between inactive and active cytosolic $Ca^{2+}$ concentrations, we systematically evaluated variants of the C-terminal SSLuc peptide, initially choosing peptides described from the development of NanoBit[11] as we had previously done during SSLuc optimization. We found that the clones displaying the highest contrast in luciferase activity between low and high [$Ca^{2+}$] contained an I13T substitution (reversion) in the SSLuc N-terminal fragment coupled with the low-affinity peptide **VTGYRLFEEIL** ("pep114"). We then designed peptides based on this core sequence, drawing inspiration from the best-performing peptides identified during the evolution of SSLuc (**Supp. Table 1**). Ultimately, this led us to choose the peptide **VTGYRLLEEISN** for the final variant, which we named "CaBLAM." The rationale for the design of this peptide, which contains two glutamate residues substituted for the preferred lysines in SSLuc, was to maintain enzymatic activity (largely from the arginine side chain that forms part of the putative active site[9]) while also reducing affinity for the large piece of the luciferase to increase the contrast in luciferase activity between the unbound and $Ca^{2+}$-bound states. We speculate this low affinity allows the C-terminus of the luciferase to disengage from the enzyme more fully under low $Ca^{2+}$ concentrations, decreasing luciferase activity at baseline.

**Supplementary Tables**

**Supplementary Table 1.** C-terminal peptide sequences.

| Peptide name | Sequence |
|---|---|
| eKAZ/Oluc | VTGWRLCENILA |
| NanoLuc | VTGWRLCERILA |
| eKL9h | VTGWRLFEKISN |
| pep101 | VTGYRLFEKES |
| pep104 | VEGYRLFEKIS |
| pep114 | VTGYRLFEEIL |
| pep128 | VTGYRLFEKIL |
| pep79 | VTGYRLFKKISN |
| pep86 | VSGWRLFKKIS |
| pep99 | VTGYRLFEKIS |
| pepNP | VTGWRLFERILA |
| LKKIS | VTGYRLLKKIS |
| LEKIS | VTGYRLLEKIS |
| LEKISN | VTGYRLLEKISN |
| LEEIS | VTGYRLLEEIS |
| LEKS | VTGYRLLEKS |
| LEKSS | VTGYRLLEKSS |
| LERILA | VTGYRLLERILA |
| CEKISN | VTGYRLCEKISN |
| IKKS | VTGYRLIKKS |
| VKKS | VTGYRLVKKS |
| CKKS | VTGYRLCKKS |
| IEKIS | VTGYRLIEKIS |
| VEKIS | VTGYRLVEKIS |
| KKSCDV | VTGYRLLKKSCDV |
| KKSCD | VTGYRLLKKSCD |
| KKSC | VTGYRLLKKSC |
| KKS | VTGYRLLKKS |
| KK | VTYYRLLKK |
| LKKISN | VTGYRLLKKISN |
| LEEISN | VTGYRLLEEISN |

Supplementary Figures

A

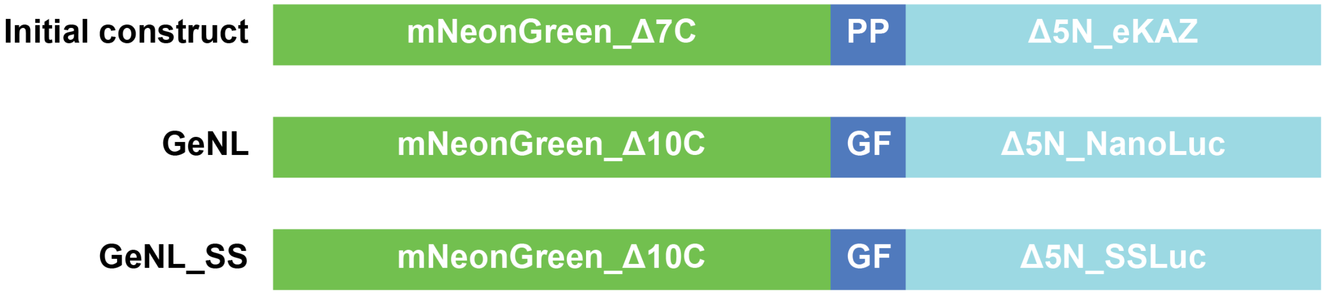

B

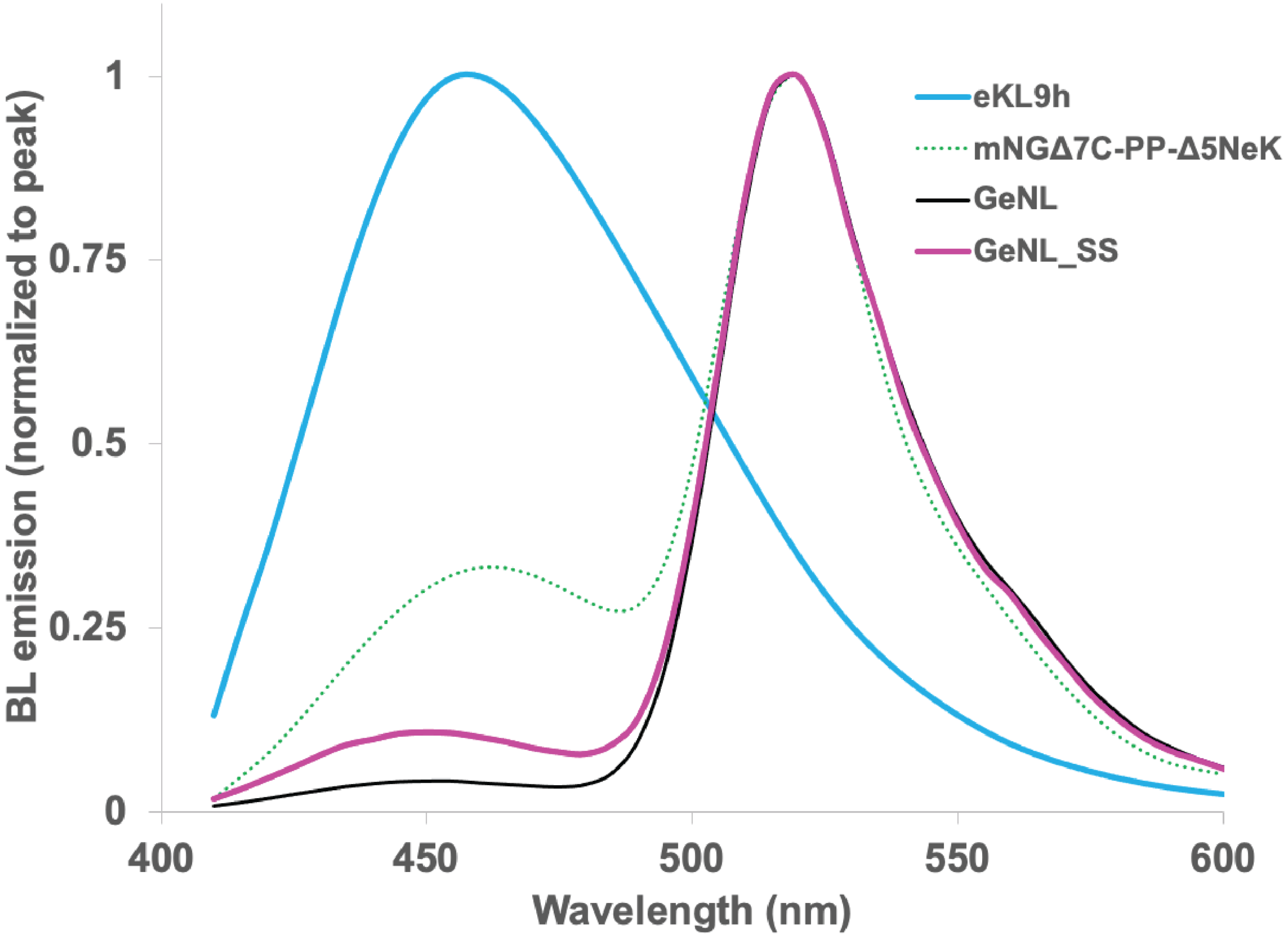

**Supplementary Figure 1.** Constructs and emission spectra. (A) Architecture of the initial mNeonGreen FRET construct with eKAZ, GeNL, and GeNL_SS. (B) Bioluminescence emission spectra for eKL9h (identical to both NanoLuc and SSLuc), initial FRET construct mNGΔ7C-PP-Δ5NeKAZ, GeNL, and GeNL_SS.

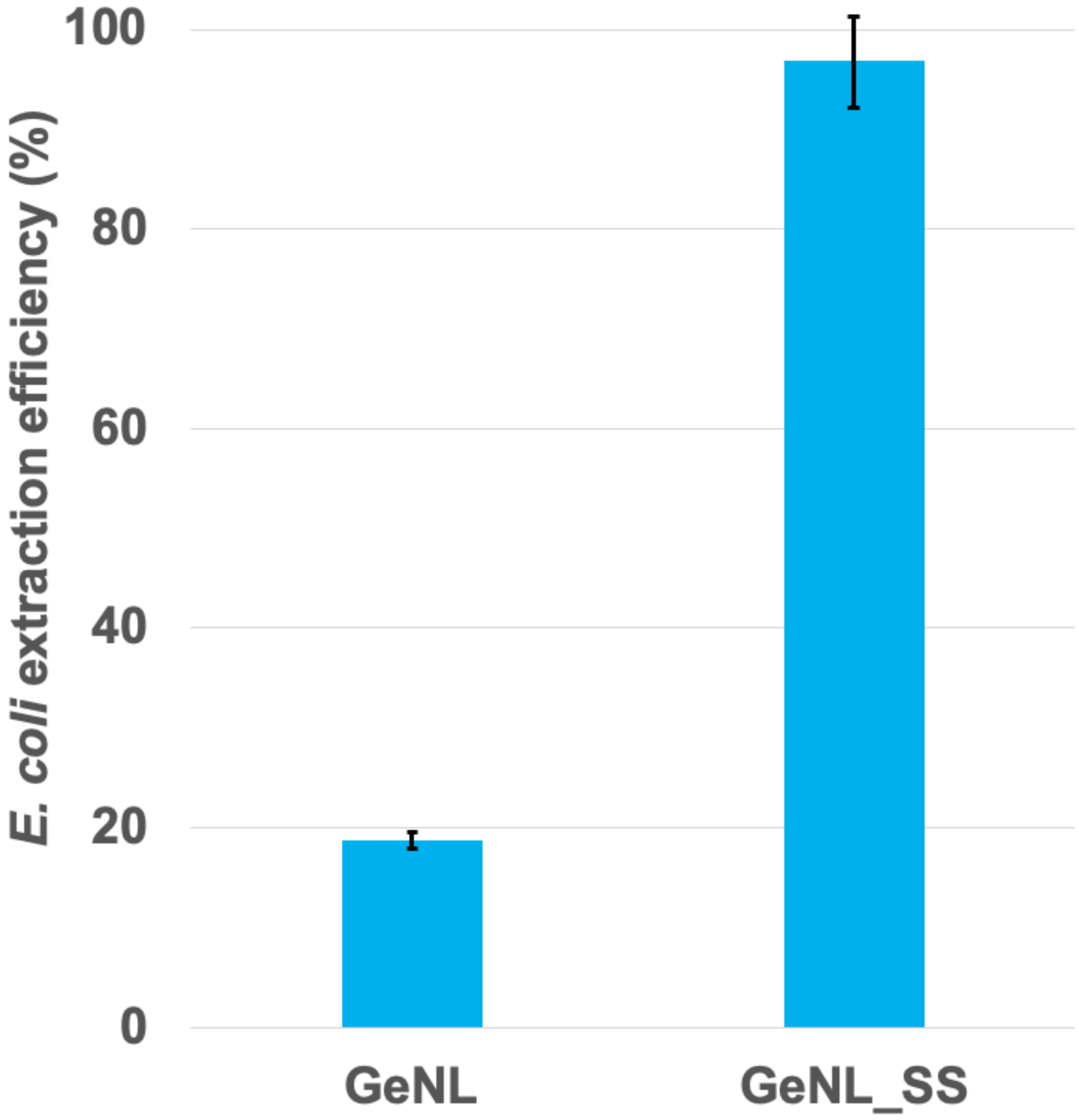

**Supplementary Figure 2.** Extraction efficiency of GeNL and GeNL_SS when expressed from the pNCST vector in *E. coli* and extracted with B-PER reagent. Values shown are the ratio between extracted fluorescence and total pellet fluorescence. Error bars represent 95%CI.

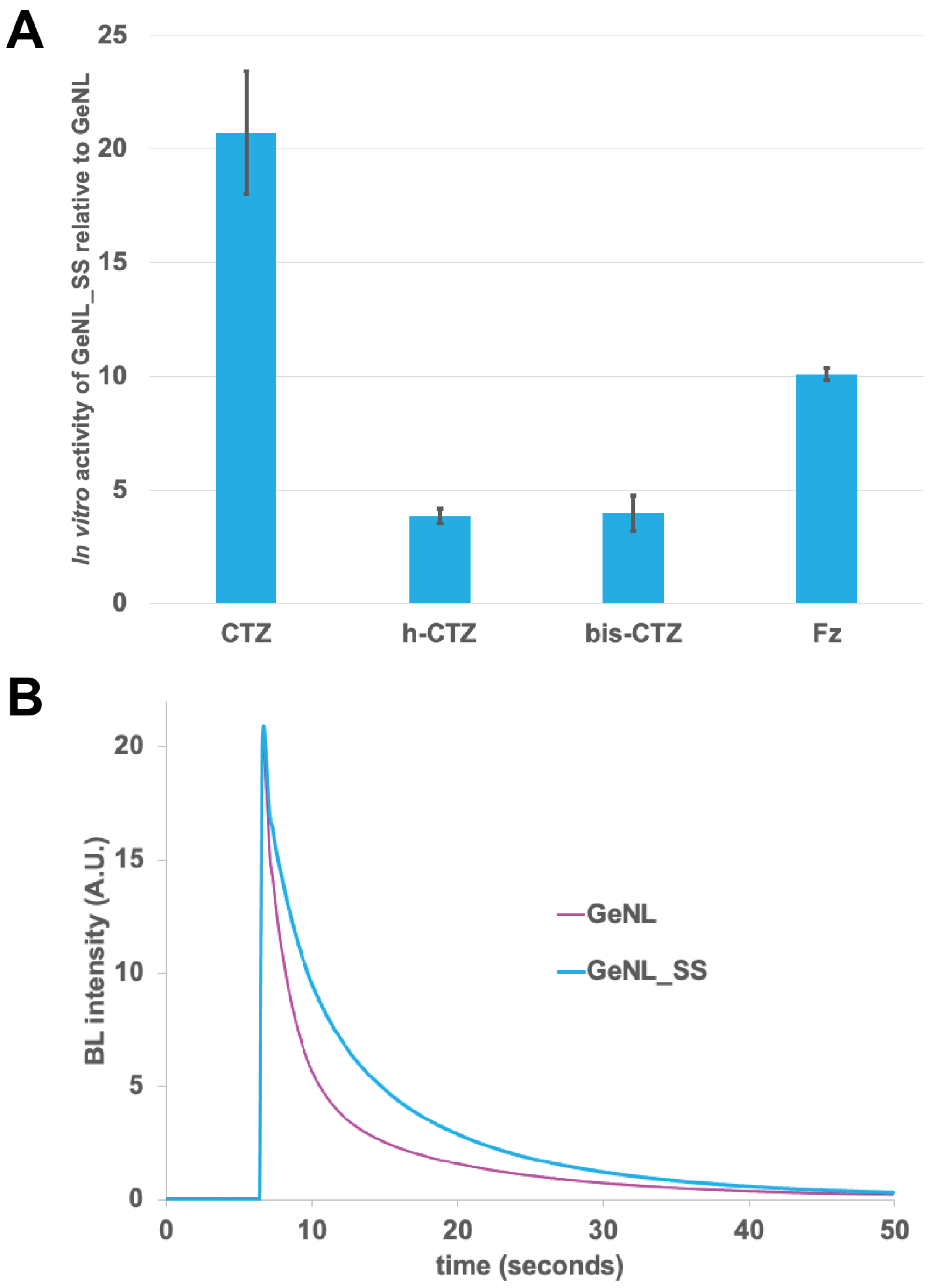

**Supplementary Figure 3.** *In vitro* characterization of SSLuc substrate preference and relative activity. (A) Relative activity with different substrates of GeNL_SS compared to GeNL. Bars represent fold difference of GeNL_SS steady-state intensity relative to equimolar GeNL in *E. coli* expression lysates. Error bars represent 95%CI. (B) Kinetic behavior of GeNL and GeNL_SS with *h*-CTZ.

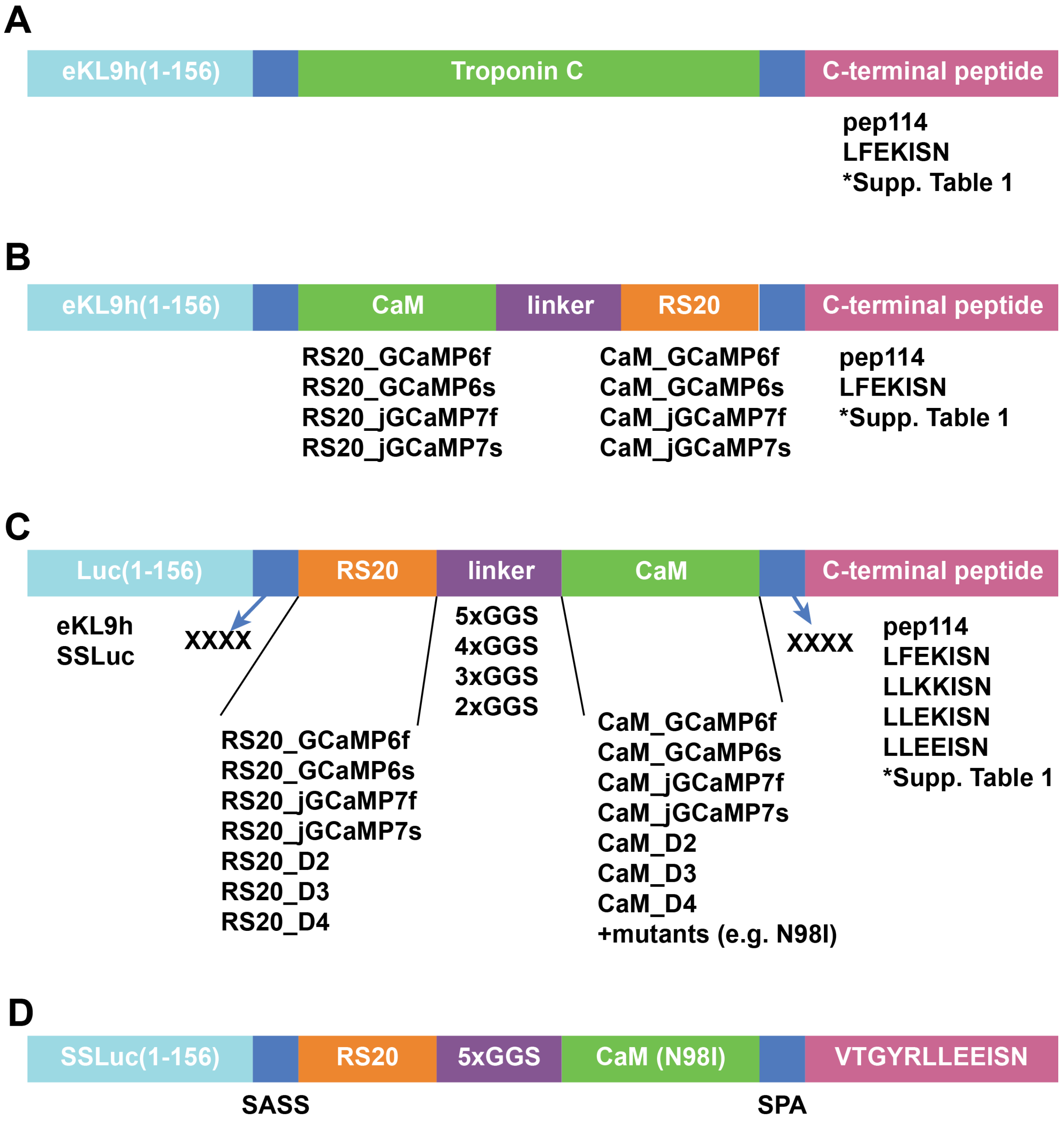

**Supplementary Figure 4.** BL GECI architectures investigated in this work. (A) Insertions of Troponin C domains produced working indicators with low contrast. (B) A calmodulin-linker-RS20 insertion into the luciferase was also functional but had low brightness regardless of the combination of components. (C) An RS20-linker-calmodulin insertion into the luciferase generated the highest brightness and contrast with tunable $Ca^{2+}$ affinity, ultimately leading us to the final CaBLAM sensor module design (D). Note that full-length CaBLAM includes the N-terminal mNeonGreen FRET acceptor, like GeNL_SS.

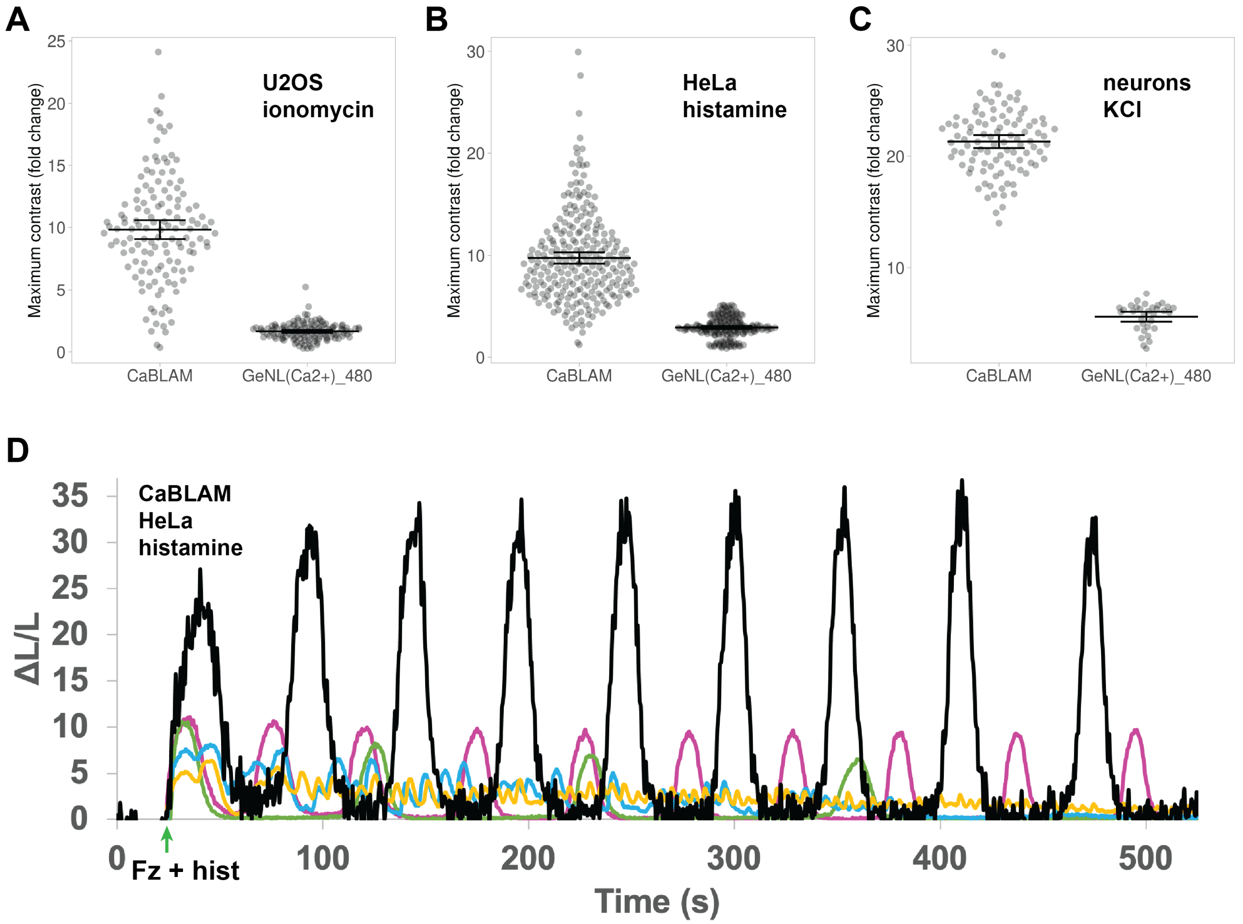

**Supplementary Figure 5.** Characterization of CaBLAM in cultured cell lines and rat cortical neurons. (A) In U2OS cells treated with ionomycin and 1mM external $Ca^{2+}$, CaBLAM consistently generated large changes in BL with ionomycin treatment with a dynamic range 5.4±0.1-fold higher (95% confidence interval (95%CI)) than that of GeNL($Ca^{2+}$)_480. (B) In HeLa cells treated with L-histamine, the maximal contrast was range 3.3±0.1-fold (95%CI) higher in CaBLAM vs. GeNL($Ca^{2+}$)_480. (C) Rat cortical neurons depolarized with KCl produced a ~20-fold increase in BL with CaBLAM, 3.8±0.1-fold higher than observed for GeNL($Ca^{2+}$)_480. For (A)-(C), Data points represent contrast (fold change) between steady-state resting BL GECI signal and peak high-$Ca^{2+}$ GECI signal after treatment. Each point corresponds to a single cell. Mean contrast is shown for each condition with error bars representing the 95%CI for each. (D) Representative CaBLAM signals in HeLa cells treated with L-histamine. Luciferin substrate (Fz) was injected along with L-histamine, indicated by the green arrow. Each colored line represents an individual cell in the same imaging field.

A

B

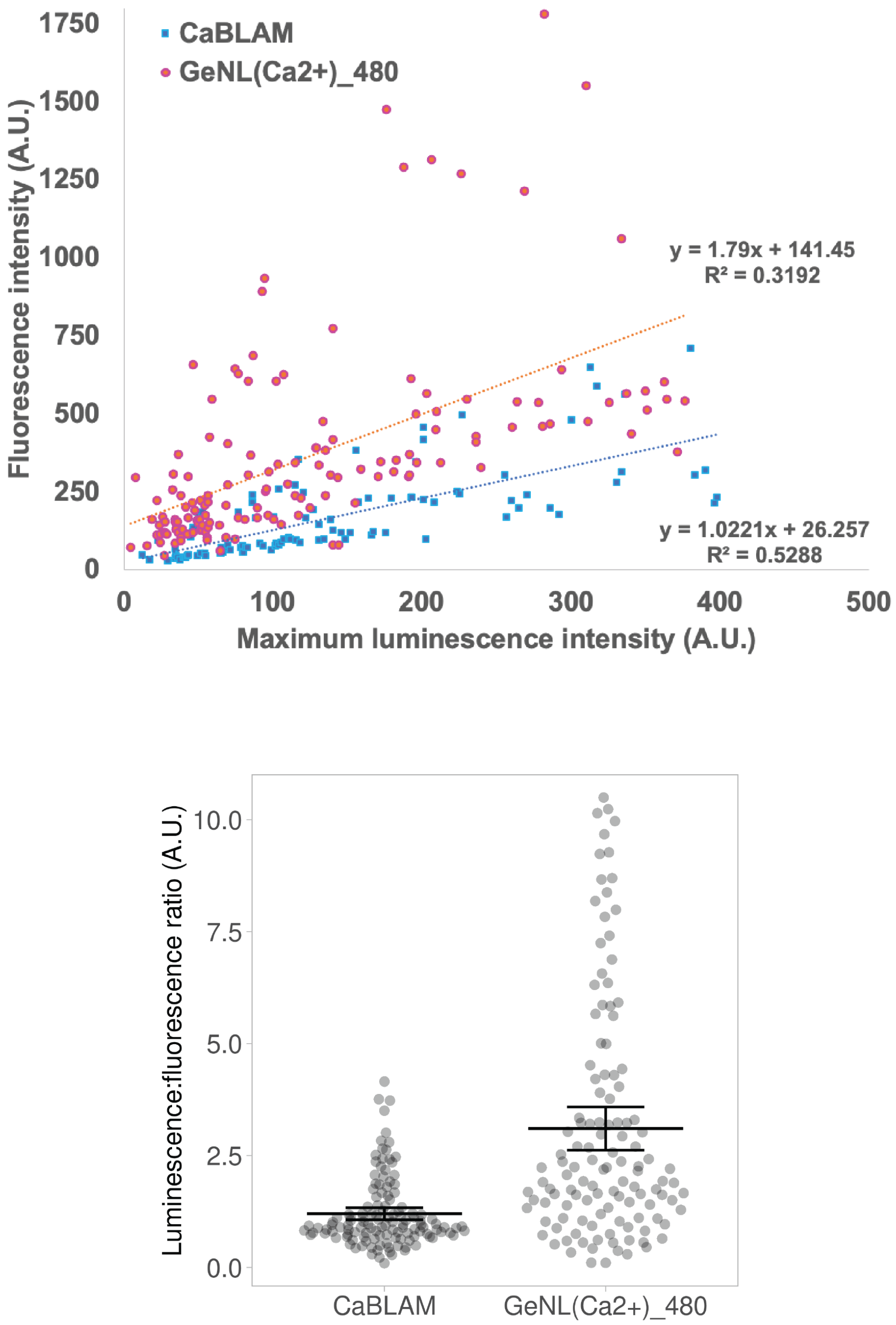

**Supplementary Figure 6. Maximal brightness comparison of CaBLAM and GeNL($Ca^{2+}$)_480 in HeLa.** (A) We observed a poor linear correlation between fluorescence intensity and maximal bioluminescence emission in cells expressing the lowest levels of each sensor, leading to a broad spread in fluorescence:bioluminescence ratios per cell (B). However, these data strongly suggest that CaBLAM is ~40% as bright as GeNL($Ca^{2+}$)_480 when both are $Ca^{2+}$-saturated.

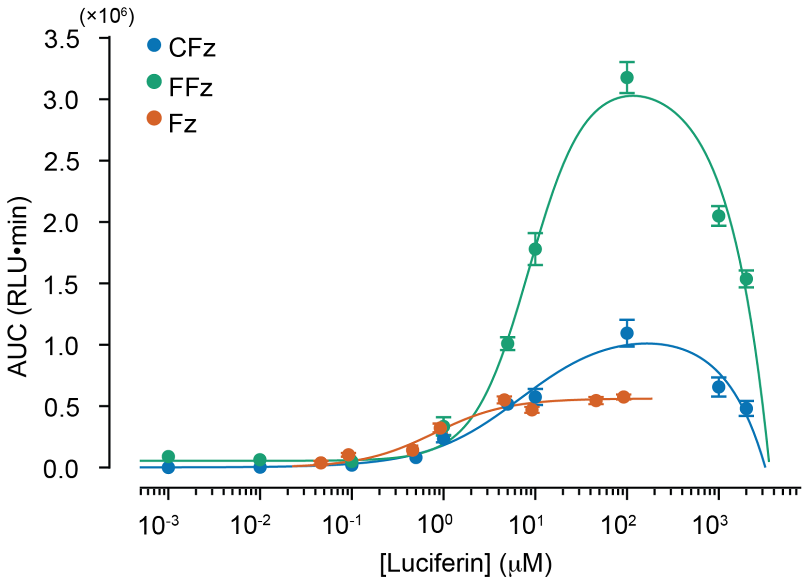

**Supplementary Figure 7. CaBLAM luminescence in cells as a function of luciferin type and concentration.** Dose-response curves for cephalofurimazine (CFz; blue), fluorofurimazine (FFz; green), and furimazine (Fz; orange) in undifferentiated N2A cells expressing CaBLAM. Luminescence expressed as relative light units (RLU) was measured from a microplate reader. Points shown are average ± SEM (N = 3 technical replicates from 2 different transfections. The area under the curve (AUC) over the 20-minute time course in the presence of substrate was averaged across the six wells. The dose-response curves for CFz and FFz were fit using the combination of a Hill plot and straight line, and for Fz a Hill plot. Fit parameters were for CFz: $EC_{50}$ = 7.0 μM, $n_H$ = 0.86, $R^2$ = 0.97. FFz: $EC_{50}$ = 8.6 μM, $n_H$ = 1.33, $R^2$ = 0.99. Fz Hill: $EC_{50}$ = 0.80 μM, $n_H$ = 1.10, $R^2$ = 0.96).

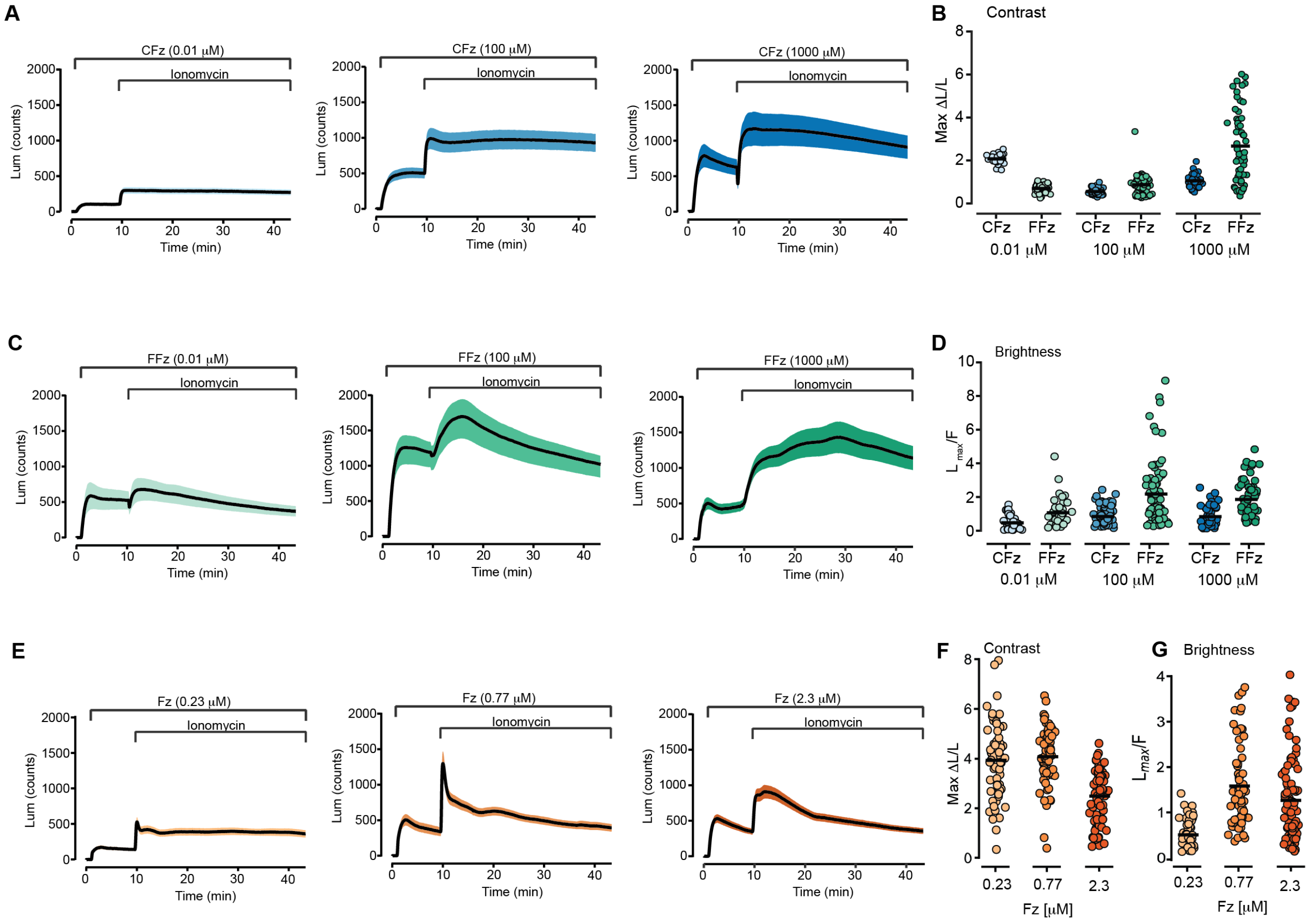

**Supplementary Figure 8. Contrast and brightness estimates for CaBLAM for different luciferins at different concentrations.** Time courses of luminescence for CFz (A, blue), FFz (C, green) and Fz (E, orange) at different concentrations applied alone and together with ionomycin in N2a cells expressing CaBLAM. Contrast (Maximum ΔL/L) and brightness ($L_{max}$/F) are shown for CFz (B, blue), FFz (D, green) and Fz (F, orange). Dark current was sampled for 60 s before adding the luciferin, and at 10 min Ionomycin was added for a final concentration of 2 μM. Data were sampled at 10 Hz with an EMCCD camera. Luminescence time courses for A) CFz at 0.01 μM (n = 39 cells), 100 μM (n = 62 cells), and 1000 μM (n = 51 cells); for C) FFz at 0.01 μM (n = 43 cells), 100 μM (n = 79 cells), and 1000 μM (n = 59 cells); and for E) Fz at 0.23 μM (n = 59 cells), 0.77 μM (n = 63 cells), and 2.3 μM (n = 85 cells). B, D, F, G Contrast and Brightness for each cell for each luciferin concentration. Points shown represent measurements from individual cells together with mean values (black bars).

**Other Supplementary Information**

**Supp Movie 1** — BL imaging of CaBLAM-expressing HeLa cells stimulated with l-histamine.

**Supp Movie 2** — BL imaging of CaBLAM-expressing rat hippocampal neurons with electrical stimulation

**Supp Movie 3** — Example movie of CaBLAM imaging for ~10 minutes (sped-up by 10x) of NDNF neurons in an awake mouse. The following processing steps were performed: A dark image was first subtracted from all images, after which a highly smoothed version of each image (convolved with a disk with a radius of 48 pixels) was then subtracted from each image to remove the signal background. Each pixel was then smoothed in time with a 10-point gaussian kernel. The pixel values are scaled to the range of the data after these processing steps.

**Supp Movie 4** — Infrared movie of head-fixed zebrafish larva illustrating tail flick movement

**Supp Movie 5** — Pixelwise standard deviation for zebrafish CaBLAM time series recorded with machine vision camera

**Supp Movie 6** — Intensified high-speed camera movie of CaBLAM BL in a head-fixed 4 dpf *Tg(elavl3)* zebrafish larva in 1:1000 vivazine

**Supp Data File 1** — peptide sequences of SSLuc, related OLucs, and intermediates, CaBLAM

**Supp Data File 2** — primer list
