## Supplementary Data File 1 (peptide sequences) for "CaBLAM! A high-contrast bioluminescent Ca^2+^ indicator derived from an engineered *Oplophorus gracilirostris* luciferase"

>SSLuc
MVFTLEDFVGDWEQIAAYNLDQVLEQGGVSSVLQTLAVSVTPIQRIVRSGENGLKIDIHVIIPYEGLSADQMA
HIEEVFKVVYPVDDHHFKVIMEYGTLVIDGVTPNMLNYFGRPYEGIAVFDGKKITVTGTLWNGNKIIDERLIT
PEGSMLFRVTINGVTGYRLLKKISN
>GeNL_SS (mNGd10C-GF-d5N_SSLuc)
MVSKGEEDNMASLPATHELHIFGSINGVDFDMVGQGTGNPNDGYEELNLKSTKGDLQFSPWILVPHIGYGFHQ
YLPYPDGMSPFQAAMVDGSGYQVHRTMQFEDGASLTVNYRYTYEGSHIKGEAQVKGTGFPADGPVMTNSLTAA
DWCRSKKTYPNDKTIISTFKWSYTTGNGKRYRSTARTTYTFAKPMAANYLKNQPMYVFRKTELKHSKTELNFK
EWQKAFTGFEDFVGDWEQIAAYNLDQVLEQGGVSSVLQTLAVSVTPIQRIVRSGENGLKIDIHVIIPYEGLSA
DQMAHIEEVFKVVYPVDDHHFKVIMEYGTLVIDGVTPNMLNYFGRPYEGIAVFDGKKITVTGTLWNGNKIIDE
RLITPEGSMLFRVTINGVTGYRLLKKISN
>CaBLAM
MVSKGEEDNMASLPATHELHIFGSINGVDFDMVGQGTGNPNDGYEELNLKSTKGDLQFSPWILVPHIGYGFHQ
YLPYPDGMSPFQAAMVDGSGYQVHRTMQFEDGASLTVNYRYTYEGSHIKGEAQVKGTGFPADGPVMTNSLTAA
DWCRSKKTYPNDKTIISTFKWSYTTGNGKRYRSTARTTYTFAKPMAANYLKNQPMYVFRKTELKHSKTELNFK
EWQKAFTGFEDFVGDWEQTAAYNLDQVLEQGGVSSVLQTLAVSVTPIQRIVRSGENGLKIDIHVIIPYEGLSA
DQMAHIEEVFKVVYPVDDHHFKVIMEYGTLVIDGVTPNMLNYFGRPYEGIAVFDGKKITVTGTLWNGNKIIDE
RLITPEGSMLFRVTINSASSDSSRRKWNKTGHAVRAIGRLSSGGSGGSGGSGGSGGSNLPDQLTEEQIAEFKE
EFSLFDKDGDGTITTKELGTVMRSLGQNPTEAELQDMINEVDADGDGTIDFPEFLTMMARKMKYRDTEEEIRE
AFGVFDKDGIGYISAAELRHVMTNLGEKLTDEEVDEMIREADIDGDGQVNYEEFVQMMTAKSPAVTGYRLLEE
ISN
>CaBLAM_294W
MVSKGEEDNMASLPATHELHIFGSINGVDFDMVGQGTGNPNDGYEELNLKSTKGDLQFSPWILVPHIGYGFHQ
YLPYPDGMSPFQAAMVDGSGYQVHRTMQFEDGASLTVNYRYTYEGSHIKGEAQVKGTGFPADGPVMTNSLTAA
DWCRSKKTYPNDKTIISTFKWSYTTGNGKRYRSTARTTYTFAKPMAANYLKNQPMYVFRKTELKHSKTELNFK
EWQKAFTGFEDFVGDWEQTAAYNLDQVLEQGGVSSVLQTLAVSVTPIQRIVRSGENGLKIDIHVIIPYEGLSA
DQMAHIEEVFKVVYPVDDHHFKVIMEYGTLVIDGVTPNMLNYFGRPYEGIAVFDGKKITVTGTLWNGNKIIDE
RLITPEGSMLFRVTINSASSDSSRRKWNKTGHAVRAIGRLSSGGSGGSGGSGGSGGSNLPDQLTEEQIAEFKE
EFSLFDKDGDGTITTKELGTVMRSLGQNPTEAELQDMINEVDADGDGTIDFPEFLTMMARKMKYRDTEEEIRE
AFGVFDKDGWGYISAAELRHVMTNLGEKLTDEEVDEMIREADIDGDGQVNYEEFVQMMTAKSPAVTGYRLLEE
ISN
>CaBLAM_332W
MVSKGEEDNMASLPATHELHIFGSINGVDFDMVGQGTGNPNDGYEELNLKSTKGDLQFSPWILVPHIGYGFHQ
YLPYPDGMSPFQAAMVDGSGYQVHRTMQFEDGASLTVNYRYTYEGSHIKGEAQVKGTGFPADGPVMTNSLTAA
DWCRSKKTYPNDKTIISTFKWSYTTGNGKRYRSTARTTYTFAKPMAANYLKNQPMYVFRKTELKHSKTELNFK
EWQKAFTGFEDFVGDWEQTAAYNLDQVLEQGGVSSVLQTLAVSVTPIQRIVRSGENGLKIDIHVIIPYEGLSA
DQMAHIEEVFKVVYPVDDHHFKVIMEYGTLVIDGVTPNMLNYFGRPYEGIAVFDGKKITVTGTLWNGNKIIDE
RLITPEGSMLFRVTINSASSDSSRRKWNKTGHAVRAIGRLSSGGSGGSGGSGGSGGSNLPDQLTEEQIAEFKE
EFSLFDKDGDGTITTKELGTVMRSLGQNPTEAELQDMINEVDADGDGTIDFPEFLTMMARKMKYRDTEEEIRE
AFGVFDKDGNGYISAAELRHVMTNLGEKLTDEEVDEMIREADIDGDGWVNYEEFVQMMTAKSPAVTGYRLLEE
ISN
>eKL9h
MVFTLEDFVGDWEQTAAYNLDQVLEQGGVSSVLQTLAVSVTPIQRIVRSGENGLKIDIHVIIPYEGLSADQMA
HIEEVFKVVYPVDDHHFKVIMEYGTLVIDGVTPNMLNYFGRPYEGIAVFDGKKITVTGTLWNGNKIIDERLIT
PDGSMLFRVTINGVSGWRLFEKISN
>eKAZ-L9
MVFTLADFVGDWQQTAGYNQDQVLEQGGLSSVFQTLGVSVTPIQKIVLSGENGLKIDIHVIIPYEGLSGDQMG
HIEMIFKVVYPVDDHHFKIIMEYGTLVIDGVTPNMIDYFGRPYPGIAVFDGKQITVTGTLWNGNKIIDERLIN
PDGSLLFRVTINGVTGWRLCENILA
>eKAZ-L6
MVFTLADFVGDWQQTAGYNQDQVLEQGGLSSLFQALGVSVTPIQKIVLSGENGLKIDIHVIIPYEGLSGFQMG
LIEMIFKVVYPVDDHHFKIIMEYGTLVIDGVTPNMIDYFGRPYPGIAVFDGKQITVTGTLWNGNKIIDERLIN
PDGSLLFRVTINGVTGWRLCENILA

```
>NanoBit::pep86
MVFTLEDFVGDWEQTAAYNLDQVLEQGGVSSLLQNLAVSVTPIQRIVRSGENALKIDIHVIIPYEGLSADQMA
QIEEVFKVVYPVDDHHFKVILPYGTLVIDGVTPNMLNYFGRPYEGIAVFDGKKITVTGTLWNGNKIIDERLIT
PDGSMLFRVTINGVSGWRLFKKIS
>NanoLuc
MVFTLEDFVGDWRQTAGYNLDQVLEQGGVSSLFQNLGVSVTPIQRIVLSGENGLKIDIHVIIPYEGLSGDQMG
QIEKIFKVVYPVDDHHFKVILHYGTLVIDGVTPNMIDYFGRPYEGIAVFDGKKITVTGTLWNGNKIIDERLIN
PDGSLLFRVTINGVTGWRLCERILA
>eKAZ
MVFTLADFVGDWQQTAGYNQDQVLEQGGLSSLFQALGVSVTPIQKIVLSGENGLKIDIHVIIPYEGLSGFQMG
LIEMIFKVVYPVDDHHFKIILHYGTLVIDGVTPNMIDYFGRPYPGIAVFDGKQITVTGTLWNGNKIIDERLIN
PDGSLLFRVTINGVTGWRLCENILA
>OLuc
MVFTLADFVGDWQQTAGYNQDQVLEQGGLSSLFQALGVSVTPIQKVVLSGENGLKADIHVIIPYEGLSGFQMG
LIEMIFKVVYPVDDHHFKIILHYGTLVIDGVTPNMIDYFGRPYPGIAVFDGKQITVTGTLWNGNKIYDERLIN
PDGSLLFRVTINGVTGWRLCENILA
```
