## Supplementary Data File 2 (primer list) for "CaBLAM! A high-contrast bioluminescent Ca^2+^ indicator derived from an engineered *Oplophorus gracilirostris* luciferase"

| Name | Sequence | Primer usage notes |
|---|---|---|
| mNGdel9C-eKAZ-R | CAG CCA GGG TGA AAA CAT CGG TAA AGG CCT TTT GCC AC | for fusions to eKAZ N-term |
| mFrdel9C-eKAZ-R | CAG CCA GGG TGA AAA CGG AGT GGC GGC CCT CG | for fusions to eKAZ N-term |
| mNGdel7N-eKAZ-F | GCG AAA ACA TCC TGG CTG ATA ACA TGG CCT CTC TCC CAG C | for fusions to eKAZ N-term |
| mFrdel10N-eKAZ-F | GCG AAA ACA TCC TGG CTG CTA TCA TCA AGG AGT TCA TGC G | for fusions to eKAZ N-term |
| eKAZ-N-F | GTT TTC ACC CTG GCT GAC TTC G | for N-terminal FP fusion constructs |
| eKAZ-mNGd9N-R | GGGAGAGAGGCCATAGGAGGAGAGATCTTTTTAAACAGACGCCAACCG | construction of eKAZ-PP-Δ9N_mNG |
| mNGd9N-PP-F | cctcctatggcctctctcccagcg | construction of eKAZ-PP-Δ9N_mNG |
| mNG_pNCST-F | gaaaacctgtacttccagggtatggtgagcaagggcgag | construction of eKAZ-PP-Δ9N_mNG |
| eKAZ-C-R | AGC CAG GAT GTT TTC GCA CAG | construction of eKAZ-PP-Δ9N_mNG |
| eKAZ-pNCST-F | gaaaacctgtacttccagggtgttttcaccctggctgacttcg | construction of mNGΔ7C-PP-Δ5N_eKAZ |
| eKAZ-mNGd7C-F | ctttaccgatgtgatgcctcctgttttcaccctggctgacttcg | construction of mNGΔ7C-PP-Δ5N_eKAZ |
| mNGd7C-PP-R | AGGAGGCATCACATCGGTAAAGGCCTTTTGC | construction of mNGΔ7C-PP-Δ5N_eKAZ |
| mNG_pNCST-R | GCAGCCGGATCAAACGTCACTTGTACAGCTCGTCCATGCC | construction of mNGΔ7C-PP-Δ5N_eKAZ |
| mNG-eK-link3X-R | GTC AGC CAG GGT GAA AAC ASB ASB ASB GGT AAA GGC CTT TTG CCA CTC C | linker optimization |
| mNG-eK-link2X-R | GTC AGC CAG GGT GAA AAC ASB ASB GGT AAA GGC CTT TTG CCA CTC C | linker optimization |
| mNG-eK-link1X-R | GTC AGC CAG GGT GAA AAC ASB GGT AAA GGC CTT TTG CCA CTC C | linker optimization |
| mNG-eK-link0X-R | GTC AGC CAG GGT GAA AAC ASB AAA GGC CTT TTG CCA CTC CTT G | linker optimization |
| eKAZ-d7C-0-F | caaaaggcctttaccgatgtgatg GTTTTCACCCTGGCTGACTTCG | linker optimization |
| eKAZ-d8C-F | caaaaggcctttaccgatgtg GTTTTCACCCTGGCTGACTTCG | linker optimization |
| eKAZ-d7C-P-F | caaaaggcctttaccgatgtgatg CCT GTTTTCACCCTGGCTGACTTCG | linker optimization |
| eKAZ-d7C-PPP-F | caaaaggcctttaccgatgtgatg CCTCCACCT GTTTTCACCCTGGCTGACTTCG | linker optimization |
| mNG-linklib-R | ATC GGT AAA GGC CTT TTG CCA CTC | linker optimization |
| mNG-2XPPX2-eK-F | GCAAAAGGCCTTTACCGATGTGATG VMKCCTCCTVMK GTTTTCACCCTGGCTGACTTCG | linker optimization |
| mNG-1XPPX2-eK-F | GCAAAAGGCCTTTACCGATGTG VMKCCTCCTVMK GTTTTCACCCTGGCTGACTTCG | linker optimization |
| mNG-0XPPX2-eK-F | GCAAAAGGCCTTTACCGAT VMKCCTCCTVMK GTTTTCACCCTGGCTGACTTCG | linker optimization |
| mNG-2XPPX1-eK-F | GCAAAAGGCCTTTACCGATGTGATG VMKCCTCCTVMK TTCACCCTGGCTGACTTCGTTG | linker optimization |
| mNG-1XPPX1-eK-F | GCAAAAGGCCTTTACCGATGTG VMKCCTCCTVMK TTCACCCTGGCTGACTTCGTTG | linker optimization |
| mNG-0XPPX1-eK-F | GCAAAAGGCCTTTACCGAT VMKCCTCCTVMK TTCACCCTGGCTGACTTCGTTG | linker optimization |
| mNG-2XPPX0-eK-F | GCAAAAGGCCTTTACCGATGTGATG VMKCCTCCTVMK ACCCTGGCTGACTTCGTTGGTG | linker optimization |
| mNG-1XPPX0-eK-F | GCAAAAGGCCTTTACCGATGTG VMKCCTCCTVMK ACCCTGGCTGACTTCGTTGGTG | linker optimization |
| mNG-0XPPX0-eK-F | GCAAAAGGCCTTTACCGAT VMKCCTCCTVMK ACCCTGGCTGACTTCGTTGGTG | linker optimization |
| PP-eKL9-F | CCTCCTGTTTTCACCCTGGC | linker optimization |
| eKL9-pNCST-F | gaaaacctgtacttccagggt GTTTTCACCCTGGCTGACTTC | linker optimization |
| eKAZ-F | GTT TTC ACC CTG GCT GAC TTC GTT G | longer N-terminal primer for eKAZ |
| eKS11-pNCST-F | gaaaacctgtacttccagggt GTTTTCACCCTGGCTGACTTCG | Construction of eKL9-S11 |
| eKS11-pNCST-R | GCA GCC GGA TCA AAC GTC A A GAG ATC TTT TTA AAC AGA CGC CAA CCG | Construction of eKL9-S11 |
| eKL9h-pNCST-F | GAA AAC CTG TAC TTC CAG GGT ATG GAG GAC TTC GTT GGT GAC TGG | Construction of eKL9h |
| eK-56-R | GTCGATTTTCAGACCGTTTTCAC | site-directed mutagenesis |
| eK-92-3-F | GACCACCACTTCAAAATCATC WKgVMK TACGGTACCCTGGTTATCGACG | site-directed mutagenesis |
| eK-92-3-R | GATGATTTTGAAGTGGTGGTCGTC | site-directed mutagenesis |
| eK-56-F | GAAAACGGTCTGAAAATCGAC RNC CACGTTATCATCCCGTACG | site-directed mutagenesis |
| eKL6E2-72-F | ATCGAAGAGATCTTCAAAGTTGTTTACC | site-directed mutagenesis |
| eKL6E2-30-R | AGAAGACAGACCACCCTGTTCCAG | site-directed mutagenesis |
| eKL6E2-30-34-F | GAACAGGGTGGTCTGTCTTCT VDKctccagRMK CTGGCTGTTTCTGTGACTCCG | site-directed mutagenesis |
| eKL6E2-68-72-R | CAA CTT TGA AGA TCT CTT CGA TMW GAG CCA TCT GAW MAG CAG ACA GAC CTT CGT ACG G | site-directed mutagenesis |
| eKL9h-22-R | CACAGAAGACACACCACCCTGTTCTRHAACCTGGTCCAGGTTGTAAGC | site-directed mutagenesis |

| | | |
|---|---|---|
| eKL9h-33-F | CAGGGTGGTGTGTCTTCTGTGCTCVAKACGCTGGCTGTTTCTGTGAC | site-directed mutagenesis |
| eKL9h-56-R | CTTCGTACGGGATGATAACGTGMNNGTCGATTTTCAGACCGTTTTCAC | site-directed mutagenesis |
| eKL9h-56-F | CACGTTATCATCCCGTACGAAG | site-directed mutagenesis |
| eKL9h-18-R | GTTGTAAGCAGCGGTCTGCTC | site-directed mutagenesis |
| eKL9h-18-19X-F | GCAGACCGCTGCTTACAAC NNKVWK CAGGTTCTGGAACAGGGTGGTG | site-directed mutagenesis |
| eKL9h-162X-R | tcagttagagatcttttcaaacagMNBCCAACCGGACACACCGTTG | site-directed mutagenesis |
| eKL9h-162-F | CTGTTTGAAAAGATCTCTAACTGACGTTTGATC | site-directed mutagenesis |
| d5n_ekl9h-F | GAGGACTTCGTTGGTGACTGGG | site-directed mutagenesis |
| mNG_eK_pep_LLKKISN_8-R | ACCAACGAAGTCTTCAAATCCGGTAAAGG | site-directed mutagenesis |
| mNG_eK_pep_LLKKISN_9A-F | GGATTTGAAGACTTCGTTGGTGCTTGGGAACAGACCGCTGCTTACAAC | site-directed mutagenesis |
| mNG_eK_pep_LLKKISN_9E-F | GGATTTGAAGACTTCGTTGGTGAATGGGAACAGACCGCTGCTTACAAC | site-directed mutagenesis |
| mNG_eK_pep_LLKKISN_9F-F | GGATTTGAAGACTTCGTTGGTTTCTGGGAACAGACCGCTGCTTACAAC | site-directed mutagenesis |
| mNG_eK_pep_LLKKISN_9G-F | GGATTTGAAGACTTCGTTGGTGGTTGGGAACAGACCGCTGCTTACAAC | site-directed mutagenesis |
| mNG_eK_pep_LLKKISN_9H-F | GGATTTGAAGACTTCGTTGGTCATTGGGAACAGACCGCTGCTTACAAC | site-directed mutagenesis |
| mNG_eK_pep_LLKKISN_9I-F | GGATTTGAAGACTTCGTTGGTATCTGGGAACAGACCGCTGCTTACAAC | site-directed mutagenesis |
| mNG_eK_pep_LLKKISN_9K-F | GGATTTGAAGACTTCGTTGGTAAGTGGGAACAGACCGCTGCTTACAAC | site-directed mutagenesis |
| mNG_eK_pep_LLKKISN_9L-F | GGATTTGAAGACTTCGTTGGTCTGTGGGAACAGACCGCTGCTTACAAC | site-directed mutagenesis |
| mNG_eK_pep_LLKKISN_9M-F | GGATTTGAAGACTTCGTTGGTATGTGGGAACAGACCGCTGCTTACAAC | site-directed mutagenesis |
| mNG_eK_pep_LLKKISN_9N-F | GGATTTGAAGACTTCGTTGGTAACTGGGAACAGACCGCTGCTTACAAC | site-directed mutagenesis |
| mNG_eK_pep_LLKKISN_9P-F | GGATTTGAAGACTTCGTTGGTCCTTGGGAACAGACCGCTGCTTACAAC | site-directed mutagenesis |
| mNG_eK_pep_LLKKISN_9Q-F | GGATTTGAAGACTTCGTTGGTCAGTGGGAACAGACCGCTGCTTACAAC | site-directed mutagenesis |
| mNG_eK_pep_LLKKISN_9R-F | GGATTTGAAGACTTCGTTGGTCGTTGGGAACAGACCGCTGCTTACAAC | site-directed mutagenesis |
| mNG_eK_pep_LLKKISN_9S-F | GGATTTGAAGACTTCGTTGGTTCTTGGGAACAGACCGCTGCTTACAAC | site-directed mutagenesis |
| mNG_eK_pep_LLKKISN_9T-F | GGATTTGAAGACTTCGTTGGTACCTGGGAACAGACCGCTGCTTACAAC | site-directed mutagenesis |
| mNG_eK_pep_LLKKISN_9V-F | GGATTTGAAGACTTCGTTGGTGTTTGGGAACAGACCGCTGCTTACAAC | site-directed mutagenesis |
| mNG_eK_pep_LLKKISN_9W-F | GGATTTGAAGACTTCGTTGGTTGGTGGGAACAGACCGCTGCTTACAAC | site-directed mutagenesis |
| mNG_eK_pep_LLKKISN_9Y-F | GGATTTGAAGACTTCGTTGGTTACTGGGAACAGACCGCTGCTTACAAC | site-directed mutagenesis |
| mNG_eK_pep_LLKKISN_9-R | GTCACCAACGAAGTCTTCAAATCCGGTAAAG | site-directed mutagenesis |
| mNG_eK_pep_LLKKISN_10A-F | GATTTGAAGACTTCGTTGGTGACGCTGAACAGACCGCTGCTTACAACCTGG | site-directed mutagenesis |
| mNG_eK_pep_LLKKISN_10D-F | GATTTGAAGACTTCGTTGGTGACGACGAACAGACCGCTGCTTACAACCTGG | site-directed mutagenesis |
| mNG_eK_pep_LLKKISN_10E-F | GATTTGAAGACTTCGTTGGTGACGAAGAACAGACCGCTGCTTACAACCTGG | site-directed mutagenesis |
| mNG_eK_pep_LLKKISN_10F-F | GATTTGAAGACTTCGTTGGTGACTTCGAACAGACCGCTGCTTACAACCTGG | site-directed mutagenesis |
| mNG_eK_pep_LLKKISN_10G-F | GATTTGAAGACTTCGTTGGTGACGGTGAACAGACCGCTGCTTACAACCTGG | site-directed mutagenesis |
| mNG_eK_pep_LLKKISN_10H-F | GATTTGAAGACTTCGTTGGTGACCATGAACAGACCGCTGCTTACAACCTGG | site-directed mutagenesis |
| mNG_eK_pep_LLKKISN_10I-F | GATTTGAAGACTTCGTTGGTGACATCGAACAGACCGCTGCTTACAACCTGG | site-directed mutagenesis |
| mNG_eK_pep_LLKKISN_10K-F | GATTTGAAGACTTCGTTGGTGACAAGGAACAGACCGCTGCTTACAACCTGG | site-directed mutagenesis |
| mNG_eK_pep_LLKKISN_10L-F | GATTTGAAGACTTCGTTGGTGACCTGGAACAGACCGCTGCTTACAACCTGG | site-directed mutagenesis |
| mNG_eK_pep_LLKKISN_10M-F | GATTTGAAGACTTCGTTGGTGACATGGAACAGACCGCTGCTTACAACCTGG | site-directed mutagenesis |
| mNG_eK_pep_LLKKISN_10N-F | GATTTGAAGACTTCGTTGGTGACAACGAACAGACCGCTGCTTACAACCTGG | site-directed mutagenesis |
| mNG_eK_pep_LLKKISN_10P-F | GATTTGAAGACTTCGTTGGTGACCCTGAACAGACCGCTGCTTACAACCTGG | site-directed mutagenesis |
| mNG_eK_pep_LLKKISN_10Q-F | GATTTGAAGACTTCGTTGGTGACCAGGAACAGACCGCTGCTTACAACCTGG | site-directed mutagenesis |
| mNG_eK_pep_LLKKISN_10R-F | GATTTGAAGACTTCGTTGGTGACCGTGAACAGACCGCTGCTTACAACCTGG | site-directed mutagenesis |
| mNG_eK_pep_LLKKISN_10S-F | GATTTGAAGACTTCGTTGGTGACTCTGAACAGACCGCTGCTTACAACCTGG | site-directed mutagenesis |
| mNG_eK_pep_LLKKISN_10T-F | GATTTGAAGACTTCGTTGGTGACACCGAACAGACCGCTGCTTACAACCTGG | site-directed mutagenesis |
| mNG_eK_pep_LLKKISN_10V-F | GATTTGAAGACTTCGTTGGTGACGTTGAACAGACCGCTGCTTACAACCTGG | site-directed mutagenesis |
| mNG_eK_pep_LLKKISN_10W-F | GATTTGAAGACTTCGTTGGTGACTGGGAACAGACCGCTGCTTACAACCTGG | site-directed mutagenesis |
| mNG_eK_pep_LLKKISN_10Y-F | GATTTGAAGACTTCGTTGGTGACTACGAACAGACCGCTGCTTACAACCTGG | site-directed mutagenesis |

| | | |
|---|---|---|
| mNG_eK_pep_LLKKISN_12-R | CTGTTCCCAGTCACCAACGAAGTCTTC | site-directed mutagenesis |
| mNG_eK_pep_LLKKISN_13A-F | GTTGGTGACTGGGAACAGGCTGCTGCTTACAACCTGGACCAGGTTC | site-directed mutagenesis |
| mNG_eK_pep_LLKKISN_13D-F | GTTGGTGACTGGGAACAGGACGCTGCTTACAACCTGGACCAGGTTC | site-directed mutagenesis |
| mNG_eK_pep_LLKKISN_13E-F | GTTGGTGACTGGGAACAGGAAGCTGCTTACAACCTGGACCAGGTTC | site-directed mutagenesis |
| mNG_eK_pep_LLKKISN_13G-F | GTTGGTGACTGGGAACAGGGTGCTGCTTACAACCTGGACCAGGTTC | site-directed mutagenesis |
| mNG_eK_pep_LLKKISN_13H-F | GTTGGTGACTGGGAACAGCATGCTGCTTACAACCTGGACCAGGTTC | site-directed mutagenesis |
| mNG_eK_pep_LLKKISN_13I-F | GTTGGTGACTGGGAACAGATCGCTGCTTACAACCTGGACCAGGTTC | site-directed mutagenesis |
| mNG_eK_pep_LLKKISN_13K-F | GTTGGTGACTGGGAACAGAAGGCTGCTTACAACCTGGACCAGGTTC | site-directed mutagenesis |
| mNG_eK_pep_LLKKISN_13L-F | GTTGGTGACTGGGAACAGCTGGCTGCTTACAACCTGGACCAGGTTC | site-directed mutagenesis |
| mNG_eK_pep_LLKKISN_13M-F | GTTGGTGACTGGGAACAGATGGCTGCTTACAACCTGGACCAGGTTC | site-directed mutagenesis |
| mNG_eK_pep_LLKKISN_13N-F | GTTGGTGACTGGGAACAGAACGCTGCTTACAACCTGGACCAGGTTC | site-directed mutagenesis |
| mNG_eK_pep_LLKKISN_13P-F | GTTGGTGACTGGGAACAGCCTGCTGCTTACAACCTGGACCAGGTTC | site-directed mutagenesis |
| mNG_eK_pep_LLKKISN_13Q-F | GTTGGTGACTGGGAACAGCAGGCTGCTTACAACCTGGACCAGGTTC | site-directed mutagenesis |
| mNG_eK_pep_LLKKISN_13R-F | GTTGGTGACTGGGAACAGCGTGCTGCTTACAACCTGGACCAGGTTC | site-directed mutagenesis |
| mNG_eK_pep_LLKKISN_13S-F | GTTGGTGACTGGGAACAGTCTGCTGCTTACAACCTGGACCAGGTTC | site-directed mutagenesis |
| mNG_eK_pep_LLKKISN_13V-F | GTTGGTGACTGGGAACAGGTTGCTGCTTACAACCTGGACCAGGTTC | site-directed mutagenesis |
| mNG_eK_pep_LLKKISN_13W-F | GTTGGTGACTGGGAACAGTGGGCTGCTTACAACCTGGACCAGGTTC | site-directed mutagenesis |
| mNG_eK_pep_LLKKISN_13Y-F | GTTGGTGACTGGGAACAGTACGCTGCTTACAACCTGGACCAGGTTC | site-directed mutagenesis |
| mNG_eK_pep_LLKKISN_145-R | AGGGGTGATCAGACGTTCGTCG | site-directed mutagenesis |
| mNG_eK_pep_LLKKISN_146A-F | GAACGTCTGATCACCCCTGCTGGTTCTATGCTGTTCCGTGTTACCATCAAC | site-directed mutagenesis |
| mNG_eK_pep_LLKKISN_146E-F | GAACGTCTGATCACCCCTGAAGGTTCTATGCTGTTCCGTGTTACCATCAAC | site-directed mutagenesis |
| mNG_eK_pep_LLKKISN_146F-F | GAACGTCTGATCACCCCTTTCGGTTCTATGCTGTTCCGTGTTACCATCAAC | site-directed mutagenesis |
| mNG_eK_pep_LLKKISN_146G-F | GAACGTCTGATCACCCCTGGTGGTTCTATGCTGTTCCGTGTTACCATCAAC | site-directed mutagenesis |
| mNG_eK_pep_LLKKISN_146H-F | GAACGTCTGATCACCCCTCATGGTTCTATGCTGTTCCGTGTTACCATCAAC | site-directed mutagenesis |
| mNG_eK_pep_LLKKISN_146I-F | GAACGTCTGATCACCCCTATCGGTTCTATGCTGTTCCGTGTTACCATCAAC | site-directed mutagenesis |
| mNG_eK_pep_LLKKISN_146K-F | GAACGTCTGATCACCCCTAAGGGTTCTATGCTGTTCCGTGTTACCATCAAC | site-directed mutagenesis |
| mNG_eK_pep_LLKKISN_146L-F | GAACGTCTGATCACCCCTCTGGGTTCTATGCTGTTCCGTGTTACCATCAAC | site-directed mutagenesis |
| mNG_eK_pep_LLKKISN_146M-F | GAACGTCTGATCACCCCTATGGGTTCTATGCTGTTCCGTGTTACCATCAAC | site-directed mutagenesis |
| mNG_eK_pep_LLKKISN_146N-F | GAACGTCTGATCACCCCTAACGGTTCTATGCTGTTCCGTGTTACCATCAAC | site-directed mutagenesis |
| mNG_eK_pep_LLKKISN_146P-F | GAACGTCTGATCACCCCTCCTGGTTCTATGCTGTTCCGTGTTACCATCAAC | site-directed mutagenesis |
| mNG_eK_pep_LLKKISN_146Q-F | GAACGTCTGATCACCCCTCAGGGTTCTATGCTGTTCCGTGTTACCATCAAC | site-directed mutagenesis |
| mNG_eK_pep_LLKKISN_146R-F | GAACGTCTGATCACCCCTCGTGGTTCTATGCTGTTCCGTGTTACCATCAAC | site-directed mutagenesis |
| mNG_eK_pep_LLKKISN_146S-F | GAACGTCTGATCACCCCTTCTGGTTCTATGCTGTTCCGTGTTACCATCAAC | site-directed mutagenesis |
| mNG_eK_pep_LLKKISN_146T-F | GAACGTCTGATCACCCCTACCGGTTCTATGCTGTTCCGTGTTACCATCAAC | site-directed mutagenesis |
| mNG_eK_pep_LLKKISN_146V-F | GAACGTCTGATCACCCCTGTTGGTTCTATGCTGTTCCGTGTTACCATCAAC | site-directed mutagenesis |
| mNG_eK_pep_LLKKISN_146W-F | GAACGTCTGATCACCCCTTGGGGTTCTATGCTGTTCCGTGTTACCATCAAC | site-directed mutagenesis |
| mNG_eK_pep_LLKKISN_146Y-F | GAACGTCTGATCACCCCTTACGGTTCTATGCTGTTCCGTGTTACCATCAAC | site-directed mutagenesis |
| mNG_eK_pep_LLKKISN_13F-F | GTTGGTGACTGGGAACAGTTCGCTGCTTACAACCTGGACCAGGTTC | site-directed mutagenesis |
| mNG_eK_pep_LLKKISN_9AF-F | GGCCCGGGATCCACCGGTCGCCACCATGTTCGGTGCTATCCAGCTGGAC | site-directed mutagenesis |
| mNG_eKpep_LLKKISN_9AF-F | CGGATTTGAAGACTTCGTTGGTGCTTTCGAACAGACCGCTGCTTACAACCTG | site-directed mutagenesis |
| mNG_eKpep_LLKKISN_9LF-F | CGGATTTGAAGACTTCGTTGGTGTGTTCGAACAGACCGCTGCTTACAACCTG | site-directed mutagenesis |
| mNG_eKpep_LLKKISN_9YF-F | CGGATTTGAAGACTTCGTTGGTTACTTCGAACAGACCGCTGCTTACAACCTG | site-directed mutagenesis |
| eK_ISN_L18I-R | CCTGTTCCAGAACCTGGTCgatGTTGTAAGCAGCGGTCTGTTCCC | site-directed mutagenesis |
| eK_ISN_L18V-R | CCTGTTCCAGAACCTGGTCaacGTTGTAAGCAGCGGTCTGTTCCC | site-directed mutagenesis |
| ek_ISN_19-F | GACCAGGTTCTGGAACAGGGTG | site-directed mutagenesis |
| eK_ISN_L22I-R | GAAGAAACACCACCCTGTTCgatAACCTGGTCCAGGTTGTAAGCAG | site-directed mutagenesis |
| eK_ISN_L22V-R | GAAGAAACACCACCCTGTTCaacAACCTGGTCCAGGTTGTAAGCAG | site-directed mutagenesis |

| | | |
|---|---|---|
| ek_ISN_23-F | GAACAGGGTGGTGTTTCTTCTGTTCTG | site-directed mutagenesis |
| eK_ISN_L52I-R | GGATGATAACATGGATGTCGATCTTgatACCGTTTTCACCAGAACGAACGATAC | site-directed mutagenesis |
| eK_ISN_L52V-R | GGATGATAACATGGATGTCGATCTTaacACCGTTTTCACCAGAACGAACGATAC | site-directed mutagenesis |
| ek_ISN_53-F | AAGATCGACATCCATGTTATCATCCCTTACG | site-directed mutagenesis |
| eK_ISN_L65A-R | GCCATCTGGTCAGCAGAagcACCTTCGTAAGGGATGATAACATGGATGTC | site-directed mutagenesis |
| ek_ISN_66-F | TCTGCTGACCAGATGGCTCATATCG | site-directed mutagenesis |
| eK_ISN_L97I-R | GGGTAACACCGTCGATAACgatGGTACCGTATTCCATGATAACCTTGAAATGATG | site-directed mutagenesis |
| eK_ISN_L97V-R | GGGTAACACCGTCGATAACaacGGTACCGTATTCCATGATAACCTTGAAATGATG | site-directed mutagenesis |
| ek_ISN_98-F | GTTATCGACGGTGTTACCCCTAACATG | site-directed mutagenesis |
| eK_ISN_L131I-R | CGATGATCTTGTTACCGTTCCAgatGGTACCGGTAACGGTGATCTTCTTACC | site-directed mutagenesis |
| ek_ISN_131-F | TGGAACGGTAACAAGATCATCGACG | site-directed mutagenesis |
| eK_ISN_L142Q-R | GAACCGTCAGGGGTGATctgACGTTCGTCGATGATCTTGTTACCGTTC | site-directed mutagenesis |
| ek_ISN_143-F | ATCACCCCTGACGGTTCTATGCTG | site-directed mutagenesis |
| eK_ISN_L65F-R | GCCATCTGGTCAGCAGAgaaACCTTCGTAAGGGATGATAACATGGATGTC | site-directed mutagenesis |
| mNG_eK_IE_22F | GAAGAAACACCACCCTGTTCttcAACCTGGTCCAGGTTGTAAGCAG | site-directed mutagenesis |
| mNG_eK_IE_22M | GAAGAAACACCACCCTGTTCatgAACCTGGTCCAGGTTGTAAGCAG | site-directed mutagenesis |
| mNG_eK_IE_22P | GAAGAAACACCACCCTGTTCcctAACCTGGTCCAGGTTGTAAGCAG | site-directed mutagenesis |
| mNG_eK_IE_22T | GAAGAAACACCACCCTGTTCaccAACCTGGTCCAGGTTGTAAGCAG | site-directed mutagenesis |
| mNG_eK_IE_22A | GAAGAAACACCACCCTGTTCgctAACCTGGTCCAGGTTGTAAGCAG | site-directed mutagenesis |
| mNG_eK_IE_22Y | GAAGAAACACCACCCTGTTCtacAACCTGGTCCAGGTTGTAAGCAG | site-directed mutagenesis |
| mNG_eK_IE_22H | GAAGAAACACCACCCTGTTCcatAACCTGGTCCAGGTTGTAAGCAG | site-directed mutagenesis |
| mNG_eK_IE_22Q | GAAGAAACACCACCCTGTTCcagAACCTGGTCCAGGTTGTAAGCAG | site-directed mutagenesis |
| mNG_eK_IE_22N | GAAGAAACACCACCCTGTTCaacAACCTGGTCCAGGTTGTAAGCAG | site-directed mutagenesis |
| mNG_eK_IE_22K | GAAGAAACACCACCCTGTTCaagAACCTGGTCCAGGTTGTAAGCAG | site-directed mutagenesis |
| mNG_eK_IE_22D | GAAGAAACACCACCCTGTTCgacAACCTGGTCCAGGTTGTAAGCAG | site-directed mutagenesis |
| mNG_eK_IE_22E | GAAGAAACACCACCCTGTTCgaaAACCTGGTCCAGGTTGTAAGCAG | site-directed mutagenesis |
| mNG_eK_IE_22W | GAAGAAACACCACCCTGTTCtggAACCTGGTCCAGGTTGTAAGCAG | site-directed mutagenesis |
| mNG_eK_IE_22R | GAAGAAACACCACCCTGTTCcgtAACCTGGTCCAGGTTGTAAGCAG | site-directed mutagenesis |
| mNG_eK_IE_22S | GAAGAAACACCACCCTGTTCtctAACCTGGTCCAGGTTGTAAGCAG | site-directed mutagenesis |
| mNG_eK_IE_22G | GAAGAAACACCACCCTGTTCggtAACCTGGTCCAGGTTGTAAGCAG | site-directed mutagenesis |
| mNG_eK_IE_22G | GAAGAAACACCACCCTGTTCggtAACCTGGTCCAGGTTGTAAGCAG | site-directed mutagenesis |
| mNG_eK-ISN_21-R | AACCTGGTCCAGGTTGTAAGCAGC | site-directed mutagenesis |
| mNG_eK-ISN_22A-F | CTTACAACCTGGACCAGGTTGCTGAACAGGGTGGTGTTTCTTCTGTTCTGC | site-directed mutagenesis |
| mNG_eK-ISN_22D-F | CTTACAACCTGGACCAGGTTGATGAACAGGGTGGTGTTTCTTCTGTTCTGC | site-directed mutagenesis |
| mNG_eK-ISN_22K-F | CTTACAACCTGGACCAGGTTAAAGAACAGGGTGGTGTTTCTTCTGTTCTGC | site-directed mutagenesis |
| mNG_eK-ISN_22N-F | CTTACAACCTGGACCAGGTTAATGAACAGGGTGGTGTTTCTTCTGTTCTGC | site-directed mutagenesis |
| mNG_eK-ISN_22Q-F | CTTACAACCTGGACCAGGTTCAGGAACAGGGTGGTGTTTCTTCTGTTCTGC | site-directed mutagenesis |
| mNG_eK-ISN_22W-F | CTTACAACCTGGACCAGGTTTGGGAACAGGGTGGTGTTTCTTCTGTTCTGC | site-directed mutagenesis |
| mNG_eK-ISN_22Y-F | CTTACAACCTGGACCAGGTTTATGAACAGGGTGGTGTTTCTTCTGTTCTGC | site-directed mutagenesis |
| eK_Y158W -R | GTTAGAGATCTTCTTCAGCAGACGCCAACCGGTAACACCGTTGATGGTAAC | site-directed mutagenesis |
| eK_159 -F | CGTCTGCTGAAGAAGATCTCTAACTGACG | site-directed mutagenesis |
| ek_TGYfL-R | GATCAAACGTCAAGATTTCTTCAACAGGAAGTAACCGGTAACACCGTTGATGGTAAC | SSLuc and CaBLAM C-terminal peptide optimization |
| ek_TGYlL-R | GATCAAACGTCAAGATTTCTTCAACAGCAGGTAACCGGTAACACCGTTGATGGTAAC | SSLuc and CaBLAM C-terminal peptide optimization |
| ek_TGYiL-R | GATCAAACGTCAAGATTTCTTCAACAGGATGTAACCGGTAACACCGTTGATGGTAAC | SSLuc and CaBLAM C-terminal peptide optimization |
| ek_TGYmL-R | GATCAAACGTCAAGATTTCTTCAACAGCATGTAACCGGTAACACCGTTGATGGTAAC | SSLuc and CaBLAM C-terminal peptide optimization |
| ek_TGYvL-R | GATCAAACGTCAAGATTTCTTCAACAGAACGTAACCGGTAACACCGTTGATGGTAAC | SSLuc and CaBLAM C-terminal peptide optimization |
| ek_TGYpL-R | GATCAAACGTCAAGATTTCTTCAACAGAGGGTAACCGGTAACACCGTTGATGGTAAC | SSLuc and CaBLAM C-terminal peptide optimization |

| | | |
|---|---|---|
| ek_TGYtL-R | GATCAAACGTCAAGATTTCTTCAACAGGGTGTAACCGGTAACACCGTTGATGGTAAC | SSLuc and CaBLAM C-terminal peptide optimization |
| ek_TGYaL-R | GATCAAACGTCAAGATTTCTTCAACAGAGCGTAACCGGTAACACCGTTGATGGTAAC | SSLuc and CaBLAM C-terminal peptide optimization |
| ek_TGYyL-R | GATCAAACGTCAAGATTTCTTCAACAGGTAGTAACCGGTAACACCGTTGATGGTAAC | SSLuc and CaBLAM C-terminal peptide optimization |
| ek_TGYhL-R | GATCAAACGTCAAGATTTCTTCAACAGATGGTAACCGGTAACACCGTTGATGGTAAC | SSLuc and CaBLAM C-terminal peptide optimization |
| ek_TGYqL-R | GATCAAACGTCAAGATTTCTTCAACAGCTGGTAACCGGTAACACCGTTGATGGTAAC | SSLuc and CaBLAM C-terminal peptide optimization |
| ek_TGYnL-R | GATCAAACGTCAAGATTTCTTCAACAGGTTGTAACCGGTAACACCGTTGATGGTAAC | SSLuc and CaBLAM C-terminal peptide optimization |
| ek_TGYkL-R | GATCAAACGTCAAGATTTCTTCAACAGCTTGTAACCGGTAACACCGTTGATGGTAAC | SSLuc and CaBLAM C-terminal peptide optimization |
| ek_TGYdL-R | GATCAAACGTCAAGATTTCTTCAACAGGTCGTAACCGGTAACACCGTTGATGGTAAC | SSLuc and CaBLAM C-terminal peptide optimization |
| ek_TGYeL-R | GATCAAACGTCAAGATTTCTTCAACAGTTCGTAACCGGTAACACCGTTGATGGTAAC | SSLuc and CaBLAM C-terminal peptide optimization |
| ek_TGYwL-R | GATCAAACGTCAAGATTTCTTCAACAGCCAGTAACCGGTAACACCGTTGATGGTAAC | SSLuc and CaBLAM C-terminal peptide optimization |
| ek_TGYsL-R | GATCAAACGTCAAGATTTCTTCAACAGAGAGTAACCGGTAACACCGTTGATGGTAAC | SSLuc and CaBLAM C-terminal peptide optimization |
| ek_TGYgL-R | GATCAAACGTCAAGATTTCTTCAACAGACCGTAACCGGTAACACCGTTGATGGTAAC | SSLuc and CaBLAM C-terminal peptide optimization |
| ek_KS_RLIRL-F | ctgttgaagaaatcttgacgtttgatccggctgc | SSLuc and CaBLAM C-terminal peptide optimization |
| eK_LLKKISN-R | GCAGCCGGATCAAACGTCAGTTAGAGATCTTCTTCAGCAGCTTGTAAC | SSLuc and CaBLAM C-terminal peptide optimization |
| ek_pep_CEKISN-R | CCGTCTGTGCGAAAAGATCTCTAACTGAcGtTTGATCCGGCTGC | SSLuc and CaBLAM C-terminal peptide optimization |
| ek_pep_CKKS-R | GGTTACCGTCTGTGCAAGAAGTCTTGAcGtTTGATCCGGCTGC | SSLuc and CaBLAM C-terminal peptide optimization |
| ek_pep_IEKIS-R | GGTTACCGTCTGATCGAAAAGATCTCTTGAcGtTTGATCCGGCTGC | SSLuc and CaBLAM C-terminal peptide optimization |
| ek_pep_IKKS-R | CGGTTACCGTCTGATCAAGAAGTCTTGAcGtTTGATCCGGCTGC | SSLuc and CaBLAM C-terminal peptide optimization |
| ek_pep_LEEIS-R | GTTACCGTCTGCTGGAAGAAATCTCTTGAcGtTTGATCCGGCTGC | SSLuc and CaBLAM C-terminal peptide optimization |
| ek_pep_LEKIS-R | GTTACCGTCTGCTGGAAAAGATCTCTTGAcGtTTGATCCGGCTGC | SSLuc and CaBLAM C-terminal peptide optimization |
| ek_pep_LEKISN-R | CCGTCTGCTGGAAAAGATCTCTAACTGAcGtTTGATCCGGCTGC | SSLuc and CaBLAM C-terminal peptide optimization |
| ek_pep_LEKS-R | GGTTACCGTCTGCTGGAAAAGTCTTGAcGtTTGATCCGGCTGC | SSLuc and CaBLAM C-terminal peptide optimization |
| ek_pep_LEKSS-R | GTTACCGTCTGCTGGAAAAGTCTTCTTGAcGtTTGATCCGGCTGC | SSLuc and CaBLAM C-terminal peptide optimization |
| ek_pep_LERILA-R | CTGCTGGAACGTATCCTGGCTTGAcGtTTGATCCGGCTGC | SSLuc and CaBLAM C-terminal peptide optimization |
| ek_pep_LKKIS-R | GTTACCGTCTGCTGAAGAAGATCTCTTGAcGtTTGATCCGGCTGC | SSLuc and CaBLAM C-terminal peptide optimization |
| ek_pep_VEKIS-R | GGTTACCGTCTGGTTGAAAAGATCTCTTGAcGtTTGATCCGGCTGC | SSLuc and CaBLAM C-terminal peptide optimization |
| ek_pep_VKKS-R | CGGTTACCGTCTGGTTAAGAAGTCTTGAcGtTTGATCCGGCTGC | SSLuc and CaBLAM C-terminal peptide optimization |
| mNG_eK_APD3A6-LLEEISN-C1-F | cagatccgctagcgctACCGGTCGCCACCATGGTGAGCAAGGGCGAGGAG | SSLuc and CaBLAM C-terminal peptide optimization |
| mNG_eK_APD3A6-LLEEISN-C1-R | GAGATCTGAGTCCGGATTACTTGTACTTAGTTAGAGATTTCTTCGAGCAGACGGTAAC | SSLuc and CaBLAM C-terminal peptide optimization |
| ek9h_GCamp6_RS20-R | TGAGCTCAGCCGACCTATAGCTC | GECI construction and directed evolution |
| NLP ekGCamp7c-R | CTCTTCAGTCAGTTGGTCCGGCAGGTTGGATCCACCAGAACCTCCGGATC | GECI construction and directed evolution |
| NLP ekGCamp7c_C' | GGAGGTTCTGGTGGATCCAACCTGCCGGACCAACTGACTGAAGAGCAGATCG | GECI construction and directed evolution |
| ekCaMPa6-R | CAA ATG TGG TAT GGC TGA TTA TGA TCA GTT ATC AAA GGA TCT CCT CGA ACA GTC | GECI construction and directed evolution |
| mNG_eK_3xGGS-R | GCT ACC GCC CGA GCC TCC ACT ACC GCC TGA GCT CAG CCG ACC TAT AGC TC | GECI construction and directed evolution |
| GF-d5N_eKL9h-F | GGATTTGAAGACTTCGTTGGTGACTGG | GECI construction and directed evolution |
| mNG_d10C-GF-eKL9h-R | CACCAACGAAGTCTTCAAATCCGGTAAAGTCCTTTTGCCACTCCTTG | GECI construction and directed evolution |
| mNGd10C_d5NeKL9h_5xGGS-R | GCTGGTCAGGCAGGTTGGATCCACCAGAACCTCCGGATC | GECI construction and directed evolution |
| ekCamp_RLLKKS-R | AGA TTT CTT CAA CAG ACG GTA ACC GGT AAC AGC AG | GECI construction and directed evolution |
| L9hyb-CaMP-GG114-F | GAGGACTTCGTTGGTGACTGGG | GECI construction and directed evolution |
| CaMP_APD-F | GTTACCATCAACTCTGCTTCTTCTGACTCTTC | GECI construction and directed evolution |
| CaMP_PEP_end-R | GCAGCCGGATCAAACGTCACAG | GECI construction and directed evolution |
| CaMP_APD_gPv-F | gttaactaTgaGGAATTCGTTCAGATGATGACCGCTAAGTC | GECI construction and directed evolution |
| CaMP_APD_gPv-R | CATCATCTGAACGAATTCCTCATAGTTAACTGGACCGTCACCGTCGATGTCAG | GECI construction and directed evolution |
| CaMP_APDs_1-F | AACCTGCCTGACCAGCTGAC | GECI construction and directed evolution |
| CaMP_APDs-A6_136-R | ACCGTCACCGTCGATGTCAGC | GECI construction and directed evolution |
| CaMP_APDs-A6_Q137P-F | CATCGACGGTGACGGTCCAGTTAACTACGAAGAATTCGTTCAGATGATGACCG | GECI construction and directed evolution |
| CaMP_APD2-A4_98-R | ACCGTCCTTGTCGAAAACACGG | GECI construction and directed evolution |

| | | |
|---|---|---|
| CaMP_APD4-A4_98-R | ACCGTCCTTGTCAGCAACACG | GECI construction and directed evolution |
| CaMP_APD2-A4_N99I-F | GTGTTTTCGACAAGGACGGTATCGGTTACATCTCTGCTGCTGAACTGC | GECI construction and directed evolution |
| CaMP_APD4-A4_N99I-F | GTTGCTGACAAGGACGGTATCGGTTACATCTCTGCTGCTGAACTGC | GECI construction and directed evolution |
| APD_rs20-R | AGAAGACAGACGACCGAAAGCACG | GECI construction and directed evolution |
| APD3A6-F | GTTCCGTGTTACCATCAACTCTGCTTC | GECI construction and directed evolution |
| APD_EIS-R | GCAGCCGGATCAAACGTCAAGAGATTTCTTCGAGCAGACGGTAAC | GECI construction and directed evolution |
| APD_ISN-R | GCAGCCGGATCAAACGTCAGTTAGAGATTTCTTCGAGCAGACGG | GECI construction and directed evolution |
| APD3A6_ISN_pCAG-R | CCTGCACCTGAGGAGTGCGGCCGCttaGTTAGAGATTTCTTCGAGCAGACGG | GECI construction and directed evolution |
| APD3_A6_LEEISN_calmodulin-F | AACCTGCCTGACCAGCTGAC | GECI construction and directed evolution |
| APD3_A6_LEEISN_calmodulin-R | GTCAGCTGGTCAGGCAGGTT | GECI construction and directed evolution |
| APD3A6-F | AACCTGCCTGACCAGCTGAC | GECI construction and directed evolution |
| APD3A6_pNCST-R | GCAGCCGGATCAAACGTCACTTAGCGGTCATCATCTGAACGAATTCTTC | GECI construction and directed evolution |
| CaBLAM_APD3-F | TCTGCTTCTTCTGACTCTTCTAAGCGTC | GECI construction and directed evolution |
| APD3A6 CaM-R | CTTAGCGGTCATCATCTGAACGAATTCTTC | GECI construction and directed evolution |
| eK-ISN -R | CTTTGTTAGCAGCCGGATCAAaCgtcaGTTAGAGATCTTCTTCAGCAGACGGTAACC | GECI linker optimization |
| eK RVTIN-R | GTTGATGGTAACACGGAACAGCATAGAAC | GECI linker optimization |
| eK SASS-F | CTGTTCCGTGTTACCATCAACTCTGCTTCTTCTGACTCATCACGTCG | GECI linker optimization |
| eK TAKSPA-R | AGCAGGAGACTTCGCTGTCATC | GECI linker optimization |
| SPA eK-F | CAGCGAAGTCTCCTGCTGTTACCGGTTACCGTCTGCTCG | GECI linker optimization |
| eK EDFV -F | GAGGACTTCGTTGGTGACTGGGAG | GECI linker optimization |
| eK KKISN -R | GTTAGAGATCTTCTTCAGCAGACGGTAACC | GECI linker optimization |
| eK RVTIN SASS-R | AGAAGAAGCAGAGTTGATGGTAACACGGAACAGCATAGAAC | GECI linker optimization |
| APD3A6_S151_F-R | GTAACCGGTAACAGCAGGGAACTTAGCGGTCATCATCTGAACGAATTCTTC | GECI linker optimization |
| APD3A6_S151_L-R | GTAACCGGTAACAGCAGGCAGCTTAGCGGTCATCATCTGAACGAATTCTTC | GECI linker optimization |
| APD3A6_S151_I-R | GTAACCGGTAACAGCAGGGATCTTAGCGGTCATCATCTGAACGAATTCTTC | GECI linker optimization |
| APD3A6_S151_M-R | GTAACCGGTAACAGCAGGCATCTTAGCGGTCATCATCTGAACGAATTCTTC | GECI linker optimization |
| APD3A6_S151_V-R | GTAACCGGTAACAGCAGGAACCTTAGCGGTCATCATCTGAACGAATTCTTC | GECI linker optimization |
| APD3A6_S151_P-R | GTAACCGGTAACAGCAGGAGGCTTAGCGGTCATCATCTGAACGAATTCTTC | GECI linker optimization |
| APD3A6_S151_T-R | GTAACCGGTAACAGCAGGGGTCTTAGCGGTCATCATCTGAACGAATTCTTC | GECI linker optimization |
| APD3A6_S151_A-R | GTAACCGGTAACAGCAGGAGCCTTAGCGGTCATCATCTGAACGAATTCTTC | GECI linker optimization |
| APD3A6_S151_Y-R | GTAACCGGTAACAGCAGGGTACTTAGCGGTCATCATCTGAACGAATTCTTC | GECI linker optimization |
| APD3A6_S151_H-R | GTAACCGGTAACAGCAGGATGCTTAGCGGTCATCATCTGAACGAATTCTTC | GECI linker optimization |
| APD3A6_S151_Q-R | GTAACCGGTAACAGCAGGCTGCTTAGCGGTCATCATCTGAACGAATTCTTC | GECI linker optimization |
| APD3A6_S151_N-R | GTAACCGGTAACAGCAGGGTTCTTAGCGGTCATCATCTGAACGAATTCTTC | GECI linker optimization |
| APD3A6_S151_K-R | GTAACCGGTAACAGCAGGCTTCTTAGCGGTCATCATCTGAACGAATTCTTC | GECI linker optimization |
| APD3A6_S151_D-R | GTAACCGGTAACAGCAGGGTCCTTAGCGGTCATCATCTGAACGAATTCTTC | GECI linker optimization |
| APD3A6_S151_E-R | GTAACCGGTAACAGCAGGTTCCTTAGCGGTCATCATCTGAACGAATTCTTC | GECI linker optimization |
| APD3A6_S151_W-R | GTAACCGGTAACAGCAGGCCACTTAGCGGTCATCATCTGAACGAATTCTTC | GECI linker optimization |
| APD3A6_S151_R-R | GTAACCGGTAACAGCAGGACGCTTAGCGGTCATCATCTGAACGAATTCTTC | GECI linker optimization |
| APD3A6_S151_G-R | GTAACCGGTAACAGCAGGACCCTTAGCGGTCATCATCTGAACGAATTCTTC | GECI linker optimization |
| APD3A6_A153_F-R | CAGACGGTAACCGGTAACGAAAGGAGACTTAGCGGTCATCATCTGAAC | GECI linker optimization |
| APD3A6_A153_L-R | CAGACGGTAACCGGTAACCAGAGGAGACTTAGCGGTCATCATCTGAAC | GECI linker optimization |
| APD3A6_A153_I-R | CAGACGGTAACCGGTAACGATAGGAGACTTAGCGGTCATCATCTGAAC | GECI linker optimization |
| APD3A6_A153_M-R | CAGACGGTAACCGGTAACCATAGGAGACTTAGCGGTCATCATCTGAAC | GECI linker optimization |
| APD3A6_A153_V-R | CAGACGGTAACCGGTAACAACAGGAGACTTAGCGGTCATCATCTGAAC | GECI linker optimization |
| APD3A6_A153_P-R | CAGACGGTAACCGGTAACAGGAGGAGACTTAGCGGTCATCATCTGAAC | GECI linker optimization |
| APD3A6_A153_T-R | CAGACGGTAACCGGTAACGGTAGGAGACTTAGCGGTCATCATCTGAAC | GECI linker optimization |

| | | |
|---|---|---|
| APD3A6_A153_Y-R | CAGACGGTAACCGGTAACGTAAGGAGACTTAGCGGTCATCATCTGAAC | GECI linker optimization |
| APD3A6_A153_H-R | CAGACGGTAACCGGTAACATGAGGAGACTTAGCGGTCATCATCTGAAC | GECI linker optimization |
| APD3A6_A153_Q-R | CAGACGGTAACCGGTAACCTGAGGAGACTTAGCGGTCATCATCTGAAC | GECI linker optimization |
| APD3A6_A153_N-R | CAGACGGTAACCGGTAACGTTAGGAGACTTAGCGGTCATCATCTGAAC | GECI linker optimization |
| APD3A6_A153_K-R | CAGACGGTAACCGGTAACCTTAGGAGACTTAGCGGTCATCATCTGAAC | GECI linker optimization |
| APD3A6_A153_D-R | CAGACGGTAACCGGTAACGTCAGGAGACTTAGCGGTCATCATCTGAAC | GECI linker optimization |
| APD3A6_A153_E-R | CAGACGGTAACCGGTAACTTCAGGAGACTTAGCGGTCATCATCTGAAC | GECI linker optimization |
| APD3A6_A153_W-R | CAGACGGTAACCGGTAACCCAAGGAGACTTAGCGGTCATCATCTGAAC | GECI linker optimization |
| APD3A6_A153_R-R | CAGACGGTAACCGGTAACACGAGGAGACTTAGCGGTCATCATCTGAAC | GECI linker optimization |
| APD3A6_A153_S-R | CAGACGGTAACCGGTAACAGAAGGAGACTTAGCGGTCATCATCTGAAC | GECI linker optimization |
| APD3A6_A153_G-R | CAGACGGTAACCGGTAACACCAGGAGACTTAGCGGTCATCATCTGAAC | GECI linker optimization |
| APD3A6_152-F | CCTGCTGTTACCGGTTACCGTC | GECI linker optimization |
| APD3A6_154-F | GTTACCGGTTACCGTCTGCTCG | GECI linker optimization |
| APD3A6_S151L_A153T-R | CAGACGGTAACCGGTAACGGTAGGCAGCTTAGCGGTCATCATCTGAACGAATTCTTC | GECI linker optimization |
| APD3A6_S151L_A153Y-R | CAGACGGTAACCGGTAACGTAAGGCAGCTTAGCGGTCATCATCTGAACGAATTCTTC | GECI linker optimization |
| APD3A6_S151L_A153S-R | CAGACGGTAACCGGTAACAGAAGGCAGCTTAGCGGTCATCATCTGAACGAATTCTTC | GECI linker optimization |
| APD3A6_S151L_A153G-R | CAGACGGTAACCGGTAACACCAGGCAGCTTAGCGGTCATCATCTGAACGAATTCTTC | GECI linker optimization |
| APD3A6_S151Q_A153T-R | CAGACGGTAACCGGTAACGGTAGGCTGCTTAGCGGTCATCATCTGAACGAATTCTTC | GECI linker optimization |
| APD3A6_S151Q_A153Y-R | CAGACGGTAACCGGTAACGTAAGGCTGCTTAGCGGTCATCATCTGAACGAATTCTTC | GECI linker optimization |
| APD3A6_S151Q_A153S-R | CAGACGGTAACCGGTAACAGAAGGCTGCTTAGCGGTCATCATCTGAACGAATTCTTC | GECI linker optimization |
| APD3A6_S151Q_A153G-R | CAGACGGTAACCGGTAACACCAGGCTGCTTAGCGGTCATCATCTGAACGAATTCTTC | GECI linker optimization |
| APD3A6_S151T_A153T-R | CAGACGGTAACCGGTAACGGTAGGGGTCTTAGCGGTCATCATCTGAACGAATTCTTC | GECI linker optimization |
| APD3A6_S151T_A153Y-R | CAGACGGTAACCGGTAACGTAAGGGGTCTTAGCGGTCATCATCTGAACGAATTCTTC | GECI linker optimization |
| APD3A6_S151T_A153S-R | CAGACGGTAACCGGTAACAGAAGGGGTCTTAGCGGTCATCATCTGAACGAATTCTTC | GECI linker optimization |
| APD3A6_S151T_A153G-R | CAGACGGTAACCGGTAACACCAGGGGTCTTAGCGGTCATCATCTGAACGAATTCTTC | GECI linker optimization |
| APD3A6_S151K_A153T-R | CAGACGGTAACCGGTAACGGTAGGCTTCTTAGCGGTCATCATCTGAACGAATTCTTC | GECI linker optimization |
| APD3A6_S151K_A153Y-R | CAGACGGTAACCGGTAACGTAAGGCTTCTTAGCGGTCATCATCTGAACGAATTCTTC | GECI linker optimization |
| APD3A6_S151K_A153S-R | CAGACGGTAACCGGTAACAGAAGGCTTCTTAGCGGTCATCATCTGAACGAATTCTTC | GECI linker optimization |
| ek_ISN_pC1-R | CTAGATCCGGTGGATCCTTACTttaGTTAGAGATTTCTTCGAGCAGACGGTAAC | mammalian vector construction |
| APD3A6_S151K_A153G-R | CAGACGGTAACCGGTAACACCAGGCTTCTTAGCGGTCATCATCTGAACGAATTCTTC | mammalian vector construction |
| APD3A6_pC1-R | GAGATCTGAGTCCGGATTACTTGTACTTACTTAGCGGTCATCATCTGAACGAATTCTTC | mammalian vector construction |
| ISN-IE_Lifeact-R | CTGATTATGATCTAGAGTCGCGGCCGCTTAGTTAGAGATCTTCTTCAGCAGACGGTAAC | mammalian vector construction |
| LifeAct-F | GGAGGAGGGGGATCCAccggtcgccaccATGGTGAGCAAGGGCGAGGAG | mammalian vector construction |
| TOMM20-FP_ins-F | GGCGGTAGCGGGGATCCACCGGTCGCCACCATGGTGAGCAAGGGCGAGG | mammalian vector construction |
| ISN-IE_H2B-C-R | GCCTCCGGAGCCTCCAGATCTGAGTCCGGAGTTAGAGATCTTCTTCAGCAGACGGTAAC | mammalian vector construction |
| ISN-IE_CytERM-R | CTGATTATGATCTAGAGTCGCGGCCGCTTAGTTAGAGATCTTCTTCAGCAGACGGTAAC | mammalian vector construction |
